# Species-specific bacterial associations emerge from stochastically assembled microbiomes in Northeastern American fireflies

**DOI:** 10.1101/2025.08.19.670969

**Authors:** Benoît Béchade, Sarah E. Lower, Sierra R. Nichols, Tanner J. Dabbert, Alison Ravenscraft

## Abstract

Many insects harbor diverse microbial communities (“microbiomes”) that can profoundly influence the biology of their host. Yet, for most insect taxa, the relative roles of stochastic events versus deterministic forces in shaping these communities remain unknown. We investigated microbiome assembly in fireflies (*Coleoptera*: *Lampyridae*) using 16S rRNA gene amplicon sequencing and quantitative PCR from 12 species and species groups collected across the Northeastern United States. We hypothesized that firefly microbiomes are predominantly structured by stochastic ecological processes, such as random microbial exposures. Consistent with this, we observed elevated normalized stochasticity ratio (NST) values for most bacteria, coupled with high intraspecific variability in bacterial abundance and composition. However, microbiomes were more similar among closely related firefly hosts, and several bacterial taxa, including endosymbionts and unusually prevalent mollicute strains, showed lower NST values, species-specific associations, and patterns linked to geography, season, and sex. These results reveal that deterministic processes—likely involving host-driven microbial filtering, microbe-microbe interactions, and reliable transmission modes—act alongside stochastic forces to shape firefly-microbe associations. By disentangling these ecological drivers in a diverse, understudied beetle lineage, our study advances understanding of the ecological and evolutionary processes structuring microbiomes in the wild.

## Introduction

Eukaryotes are constantly exposed to a plethora of microbes in their environment but only a small fraction of those microbes associate with a host in recurrent symbioses (McFall-Ngai et al. 2013). As in any community of organisms, the assembly of microbial symbiont communities (“microbiomes”) associated with eukaryotic hosts is shaped by stochastic and deterministic processes. Stochastic forces include random spatiotemporal variations, microbial dispersal events, and priority effects (i.e., order of microbial colonizers in hosts) (Jones et al. 2022; Meadow et al. 2013; Sloan et al. 2006; Zhou and Ning 2017), whereas deterministic forces involve microbe-microbe interactions (e.g., competition, facilitation) and host/habitat filtering— within-host conditions or microbial traits that favor the establishment of certain microbes (Coyte et al. 2021; Mazel et al. 2018; Oliphant et al. 2019). However, not all microbial symbionts are the same. Specialized symbionts form species-specific associations (i.e., one microbial strain for one host species) and evolve adaptations to a host-associated lifestyle (Kwong and Moran 2015), such as reliable transmission modes (Michalik and Szklarzewicz 2025). They can influence host physiology and ecology, either in beneficial or harmful ways (Bäumler and Fang 2013; Dillon and Dillon 2003; Hammer 2024), and their establishment is mainly driven by deterministic processes (Bäumler and Fang 2013; Mazel et al. 2018). Conversely, symbiont communities with lower host specialization, typically including microbes acquired either from the environment or from physical contact with other hosts, provide more flexibility to the host but rely on more hazardous community assembly events, where stochastic processes are predominant (Perreau and Moran 2022). To understand how symbiosis between microbes and eukaryotes emerge and function as stable associations, it is crucial to identify the importance of each of these ecological processes and their interplay.

Fireflies (*Coleoptera*: *Lampyridae*) comprise >2,600 species worldwide (Keller 2025), of which 179 have been described in the US and Canada (Fallon et al. 2021; FireflyAtlas.org 2025). Seminal work using culture-based assays has shown that these insects commonly harbor bacteria from the *Mollicutes* class (Hackett et al. 1992; Stevens et al. 1997; Tully et al. 1989; 1994; Williamson et al. 1990), suggesting a role of deterministic processes in developing stable microbe-host associations. This discovery was recently supported through sequencing-based studies which hinted at possible species-specific associations between a few firefly species and certain bacterial genera (Fallon et al. 2018; Green et al. 2021). Other microbes have also been sampled from fireflies, including common insect endosymbionts (i.e., symbionts residing inside the body or cells of another organism) (Jeong et al. 2009; Jeyaprakash and Hoy 2000) and likely environment-derived bacteria (Green et al. 2021; Zhao et al. 2023), suggesting a role of stochastic processes. However, a more extensive characterization of the firefly microbiome across degrees of relatedness, species lifestyles, and environmental factors is required to better comprehend the nature of firefly-bacteria associations and identify the contribution of ecological forces to their microbiome assembly.

While the biology of a large proportion of firefly species is uncharacterized, these insects exhibit diverse life histories and ecologies (e.g., Faust 2017; Lloyd 2018) that can affect, or be affected by, associated microbial communities in deterministic ways. First, as holometabolous insects that include larval, pupal, and adult stages, the ecology and physiology of individual fireflies change drastically throughout their lifetime, potentially resulting in different microbiomes at each developmental stage. In addition, while the adult lifespan is only a few weeks in most firefly species, it is much longer in some, such as the winter firefly (*Photinus corruscus*) which can overwinter as adults for over 10 months (Deyrup et al. 2017; Rooney and Lewis 2000). Age may be a key factor impacting resistance to bacterial pathogen infections in fireflies (Lower et al. 2023), but little is known about its influence on the firefly microbiome in the wild. Second, while almost all firefly larvae are predators of soft invertebrates like snails (Buschman 1984a; 1984b; 1988; Fu and Benno Meyer-Rochow 2013; Hess 1920; Vaz et al. 2021; 2020; Yang et al. 2024), adult diets are diverse across species and may impose drastically divergent physiochemical conditions upon microbial gut colonizers. For instance, members of the *Photuris versicolor* species group prey on other non-photurine fireflies (Lewis et al. 2011; Lloyd 1965), whereas adult *Pn. corruscus* consume sap and nectar (Rooney and Lewis 2000). Most other firefly species, including *Lucidota* spp., *Pyractomena* spp., and most *Photinus* spp., are assumed not to eat as adults in the wild (Lloyd 2005; Wing 1989; but see Faust 2017; Faust and Faust 2014; Lloyd 1998; Othman et al. 2018). Third, while bioluminescence is used by all firefly species at the larval stage as a predator avoidance mechanism (Oba et al. 2020; Powell et al. 2022), some species have lost the ability to emit light as adults over evolutionary time (Martin et al. 2017; Stanger-Hall and Lloyd 2015; Stanger-Hall et al. 2007; Zaragoza-Caballero et al. 2023). Bacterial symbionts are often implicated in bioluminescence of invertebrates (Cassells et al. 2024; Jones and Nishiguchi 2004), but nothing is known about microbial involvement in firefly bioluminescence, nor about how microbiomes change between hosts that emit light as adult and those that do not. With all these specificities, fireflies provide a rich system to understand interactions between the microbiota and host ecologies, yet little is known about specialization of microbes to these insects.

Here, focusing on field-collected fireflies from 12 common species and species groups from the Northeastern United States (Figure 1A and 1B; Figure S1; Table S1), we investigate the degree of host-symbiont specificity and the contribution of diverse factors in structuring firefly microbiomes, such as host species relatedness, diet, bioluminescence, season, geography, urbanization, sex, life stage, and intraindividual tissue localization. Through a combination of quantitative (q)PCR and amplicon sequencing of the bacterial 16S rRNA gene (Figure S2), we tested the hypothesis that most bacteria composing firefly microbiomes are non-specific and these communities are mainly shaped by stochastic forces, except for host species-restricted bacterial strains from the *Mollicutes* which establish through deterministic processes. We predict that while the relative abundance of most bacteria and total bacterial titers vary in fireflies, most firefly species host their own mollicute taxon. This study, focusing on a beetle group that includes differing degrees of symbiont specialization, provides an important contrast to other beetles and paves the way for future research on bacterial transmission modes and influence of microbial symbionts on firefly biology.

**Figure 1:**
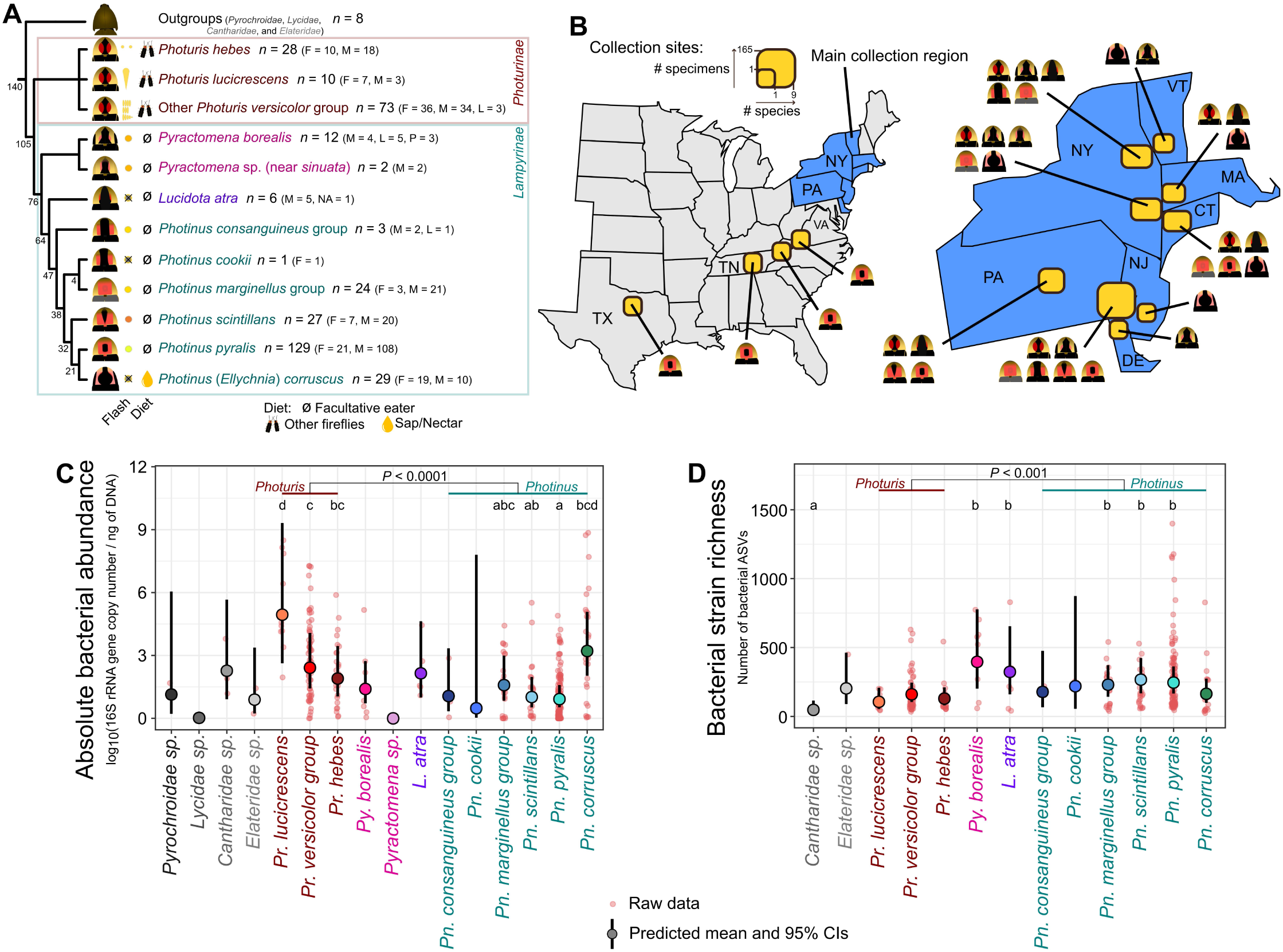
Sampling overview and firefly microbiome abundance and diversity across species. **A**. Species sampled and analyzed in the present study. The phylogenetic tree is adapted from Catalán et al. (2022) and Martin et al. (2019). Numbers at each node indicate estimated divergence time from Catalán et al. (2025), Höhna et al. (2024), and Powell (2022) in millions of years. The cartoons next to the tree branches are simplified representations of the typical markings visible on the pronotum of different firefly species. Symbols next to these pronotal depictions correspond to the most common male flash patterns for the species or species group, with the frequency of the flashes for *Photuris* species, and whether a firefly is bioluminescent (colored flash) or not (crossed out). The next symbols correspond to adult diets, with “ø” for species that are not known to eat as adults, other fireflies for predatory diet, and a yellow droplet for sap and nectar drinkers. The number of specimens collected and processed is shown next to species’ names, with “F” for females, “L” for larvae, “M” for males, and “P” for pupae. **B**. Map of the central and eastern United States and Northeastern states showing the location of sampling sites. While a few *Pn. pyralis* specimens were collected in Tennessee, Texas, and Virginia, most fireflies were collected in Northeastern states (blue color). The yellow rectangles give the number (“#”) of species (width) and specimens (height) collected in the region and the cartoons of firefly pronotum indicate which species were collected, corresponding to the cartoons from panel A. Specimen information can be found in Table S1. **C**. Normalized absolute bacterial abundance across all firefly and related insect samples. Values are expressed as the log_10_ of the number of 16S rRNA gene copies in a sample, measured through qPCR, minus the number of 16S rRNA gene copy in the DNA extraction blank sample from the same batch, and then divided by the total DNA concentration in the sample, which was measured via Qubit. Least-square means (emmeans) post-hoc tests on generalized linear mixed models (GLMMs), were used for statistical testing of differences between genera and species, and controlled for sex, day of year, month of capture, time of capture, longitude at capture, local sampling point, and whether the specimen was dissected or not. Lowercase English letters indicate the significance of the difference across firefly species. Letters for species that are not significantly different from any other species (i.e., *P* > 0.05: “abcd”) were removed. Balls and error bars correspond to predicted average and 95% confidence intervals obtained from the GLMM. Dots represent raw data points. **D**. Bacterial richness, expressed as the number of bacterial ASVs, across firefly species. The *Photinus* microbiome contains, on average, a higher bacterial strain richness than that of *Photuris* (emmeans post-hoc tests on GLMMs, controlling for genus, sex, local sampling point, day of year, year of capture, and sequencing type). See description of panel D for details on graph components. All alpha diversity metrics can be found in Table S4 and Figure S4.

## Results

### 1. The microbiome of fireflies is mostly structured by stochastic processes, except when bacteria from the *Mollicutes* are included

To assess the relative importance of deterministic *vs.* stochastic factors in structuring firefly bacterial communities, we calculated and compared the normalized stochastic ratio (NST; Ning et al. 2019) across species that had more than two specimens in our dataset. We found that the microbiome of all nine firefly species tested had an NST index below 0.5 (average = 0.27, SD = 0.10, median = 0.26; Figure 2; Table S2), indicating that their microbiome is structured by deterministic processes. To infer which bacterial group may be responsible for deterministic community assembly, we next removed all the *Mollicutes* from our datasets, and found that the NST significantly increased for five out of nine firefly species (Wilcoxon signed-rank test for these five species: *W* < 2.3 x 10^4^, *P* < 0.05), reaching index values above 0.5 (for these five species: average = 0.62, SD = 0.09, median = 0.58). This suggests that for several firefly species, while stochastic factors are involved in structuring most of the bacterial community, deterministic processes shape the relative abundance of mollicutes, influencing the whole microbiome. Interestingly, this was not the case for members of the *Photinus marginellus* group, for which removing bacteria known as endosymbionts (i.e., symbionts residing inside the body or cells of another organism) of arthropods (i.e., *Candidatus* Hepatincola, *Rickettsia*, *Rickettsiella*, *Wolbachia*) instead significantly increased the NST index, reaching a value of 0.56 (Wilcoxon signed-rank test: *W* = 370, *P* < 0.001). This indicates that for this firefly species group, deterministic processes shape the abundance of endosymbionts, not mollicutes.

**Figure 2:**
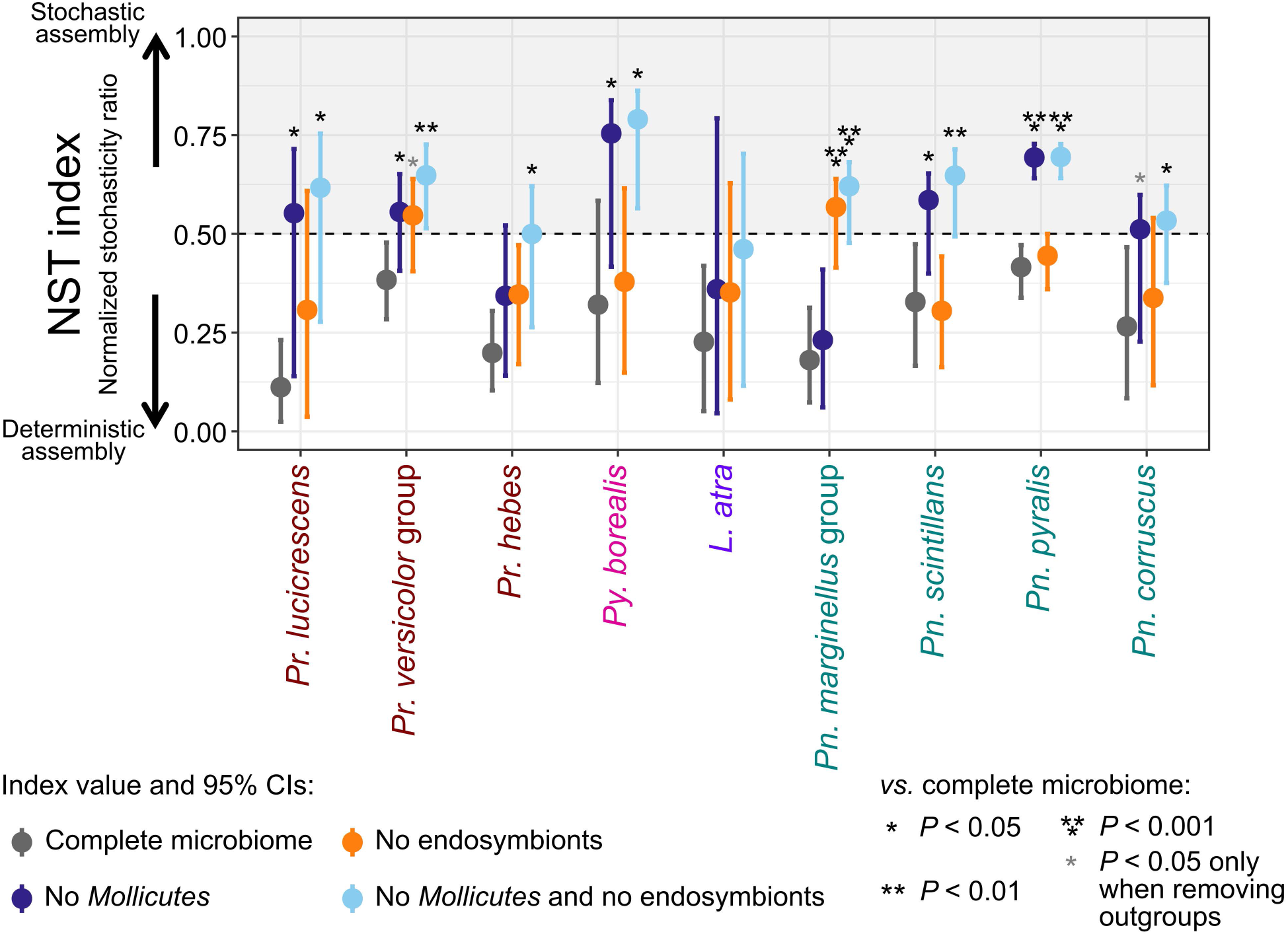
Normalized stochasticity ratio (NST) across firefly species. NST index values higher than 0.5 indicate a higher importance of stochastic forces governing bacterial community assembly, whereas values lower than 0.5 point to deterministic forces as the main drivers of community assembly. For each firefly species or species group the balls correspond to the index value and the error bars are 95% confidence intervals obtained from bootstrapping. Dark grey color is for the complete, untouched microbiome, dark blue corresponds to microbiomes where all the mollicutes were removed from the dataset, orange is for microbiomes where putative endosymbionts (*Candidatus* Hepatincola, *Rickettsia*, *Rickettsiella*, and *Wolbachia*) were removed, and light blue corresponds to microbiome datasets where both mollicutes and endosymbionts were removed. The asterisks depict, for each species, significantly different NST values between the complete microbiome and either of the modified microbiome datasets. Two-tailed Wilcoxon signed-rank statistical tests were used to test the significance of these differences. Asterisks in light grey are for tests run on datasets where outlier data points (i.e., values higher or lower than 1.5-fold of the interquartile range) were removed. Values and statistical test results can be found in Table S2.

### 2. The microbiome of fireflies is phylogenetically structured

#### 2.a. Firefly genera differ in their bacterial density and diversity

Using qPCR, we found that *Photuris* fireflies had significantly higher bacterial densities than *Photinus* fireflies (emmeans: *z*-ratio = 4.64, *P* < 0.0001; Table S3), despite high intraspecific variability (Figure 1C; Figure S3). The *Pyractomena* and *Lucidota* genera were much less represented in our dataset and members of these groups appear to have intermediate but variable bacterial levels. When looking within firefly genera, we found that *Pn. corruscus* differed from other *Photinus* species in that specimens from this long-lived adult, day-active species had three orders of magnitude higher bacterial titers, reaching levels close to those of *Pr. lucicrescens*. The difference between *Pn. corruscus* and other *Photinus* species was, however, only significant when comparing bacterial titers of *Pn. corruscus* to *Pn. pyralis* (emmeans: *z*-ratio = 3.67, *P* < 0.05). This indicates that, with the exception of *Pn. corruscus*, members of the *Photinus* genus tend to be infected with low to extremely low bacterial densities in the wild, while *Photuris* fireflies have moderately high to very high bacterial titers.

In terms of bacterial richness, despite having relatively low bacterial densities, fireflies from the *Photinus* genus harbored significantly more bacterial ASVs than *Photuris* fireflies (emmeans: *z*-ratio = 4.38, *P* < 0.001; Figure 1D; Figure S4; Table S3 and S4), suggesting that fireflies with higher bacterial densities tend to harbor fewer bacterial species.

#### 2.b. Bacterial composition differs across firefly subfamilies, genera, and species

Following rarefying and aggregation at the bacterial taxon level, we tested differences across microbiome compositions. The microbiome of Northeastern fireflies differed between the *Lampyrinae* and *Photurinae* subfamilies (PERMANOVA: *R^2^* = 0.07, *F*_1,264_ = 22.95, *P* < 0.001; Figure 3A; Table S3) and across genera (nested within subfamilies, PERMANOVA: *R*^2^ = 0.02, *F*_2,264_ = 3.87, *P* < 0.001; Figure 3B). Despite high intraspecific variability, bacterial composition also differed significantly across firefly species (nested within genera, within subfamilies, PERMANOVA: *R*^2^ = 0.09, *F*_7,264_ = 4.18, *P* < 0.001; Figure 3C–3F). To assess whether firefly microbiomes were more divergent in species that were less genetically related, we examined how bacterial community composition varied with degree of host relatedness. We found a strong positive correlation between pairwise microbiome dissimilarities and host genetic divergence, with closely related firefly species having more similar microbiome composition and abundance (partial Mantel test controlling for geographic distance: *r* = 0.37, *P* < 0.001; Figure S5), further indicating phylogenetically structured microbiomes in fireflies.

**Figure 3:**
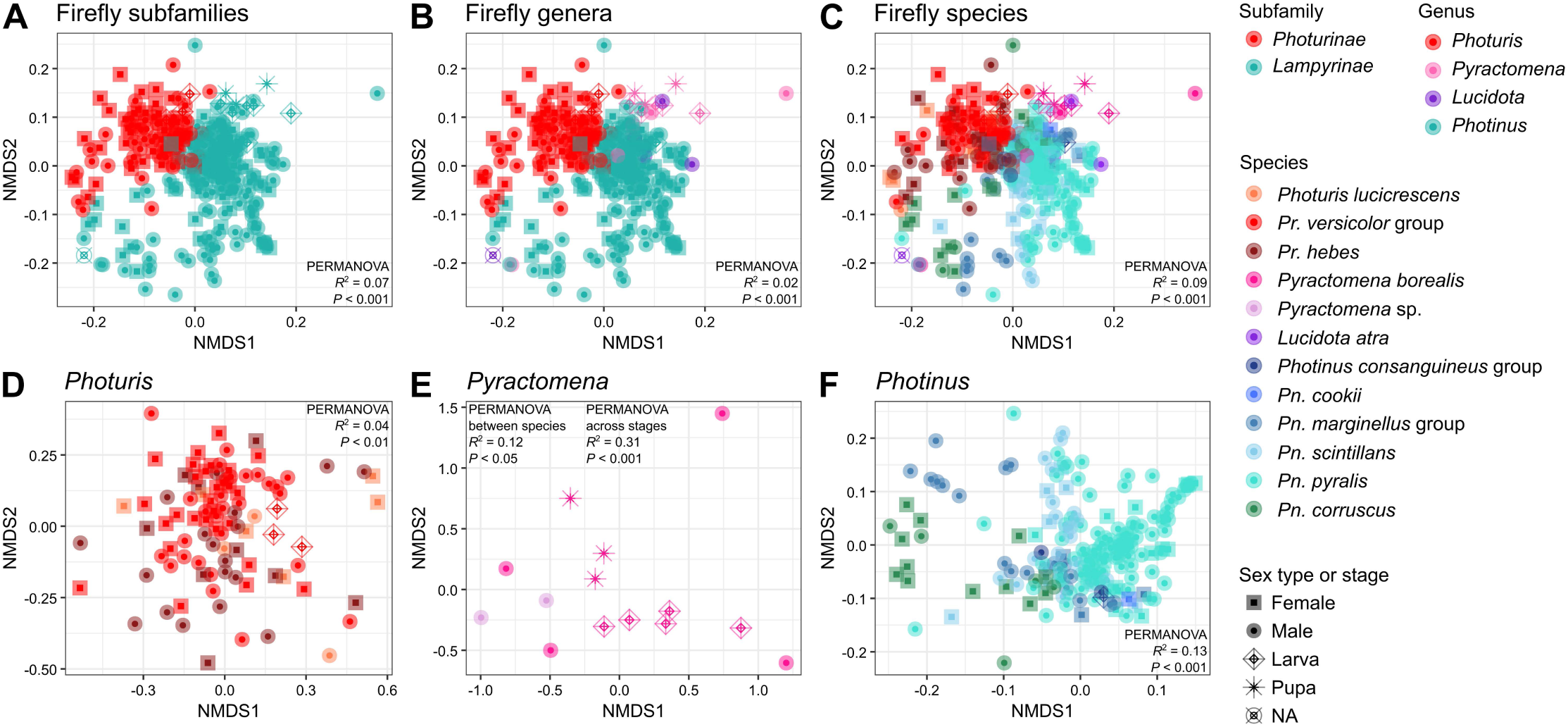
Microbiome divergence across fireflies at different taxonomic levels. **A**, **B** and **C**. Non-metric multidimensional scaling (NMDS) analyses showing separation of the bacterial communities based on composition and relative abundance at the firefly subfamily, genus, and species levels. Four outgroup specimens were removed from these analyses. **D**, **E**, and **F**. NMDS analyses showing variation in the microbiome across species within three firefly genera. All plots are derived from Bray-Curtis distances calculated on non-rarefied, proportional (i.e., compositional) relative abundance of taxon-aggregated bacteria in firefly specimens. Permutational multivariate analyses of variance (PERMANOVAs) were used for statistical testing of group similarities. Nested PERMANOVAs were used for B and C, with genus nested within subfamily and species nested within genus within subfamily, respectively. In panels D and F, the species term was the main predictor in the PERMANOVA models. PERMANOVAs were computed on read depths rarefied at 30,000 reads, except in E where it was computed on non-rarefied data to be able to compare the two *Pyractomena* species (*Pyractomena* sp. near *sinuata* specimens are filtered out by rarefying) or three developmental stages.

#### 2.c. Firefly species that include individuals with bigger size tend to harbor higher bacterial densities

We investigated whether three key characteristics of firefly species—the use of bioluminescence as a mating signal by adults, adult diet, and adult body size—influenced firefly microbiomes when taking into account fireflies’ degree of relatedness. Through phylogenetic generalized least square models, we found that bacterial density, richness, and composition did not significantly change between bioluminescent and non-bioluminescent fireflies and between species eating and not eating at the adult stage (Figure S6; Supplementary Text 1; Table S3). We also found that bigger firefly species tend to harbor a higher bacterial density and a lower bacterial richness than smaller species (Figure S7; Supplementary Text 1). These results show that firefly microbiomes do not appear to be influenced by mating signals and adult diet, but that species’ body size may contribute.

### 3. Firefly microbiomes are often dominated by a few bacterial taxa with signs of species-specific associations

The most abundant bacterial order associated with all fireflies in our amplicon sequencing dataset was *Entomoplasmatales* (class *Mollicutes*), representing 27.3% of all rarefied bacterial reads from fireflies. Highly represented bacterial orders in fireflies also included *Burkholderiales* (9.5% of all rarefied reads), *Xanthomonadales* (8.6%), *Pseudomonadales* (8.3%), *Enterobacterales* (7.7%), and *Rickettsiales* (7.0%). Compared to non-lampyrid elateroid beetles, only a few of the abundant firefly-associated bacterial taxa were similar (Figure S8; Supplementary Text 1).

#### 3.a. Fireflies harbor a diversity of mollicute taxa with prevalent and abundant strains and a few endosymbionts

The fireflies we collected harbored strains from almost all the mollicute taxa that have been sampled from fireflies in the past (seven out of eight; Figure 4; Figure S9; Supplementary Text 1). Both *Spiroplasma ixodetis* and *Williamsoniiplasma lucivorax* species were represented with a high strain diversity in our firefly specimens, with at least 20 and 30 ASVs each, respectively. Despite this train diversity, within several mollicute genera and species, one or two ASVs could be very abundant and prevalent in fireflies, some of which have only been sampled from firefly bodies to date and thus appear endemic to fireflies. While the high abundance of mollicutes associated with fireflies is unusual for an insect microbiome, we note that several prevalent and abundant bacteria in these insects are known to be endosymbionts in other arthropods (Figure 5A; Figure S10). These include bacteria from the *Rickettsia*, *Rickettsiella*, *Wolbachia*, and occasionally *Serratia* and *Spiroplasma* genera (Russell et al. 2013; Werren et al. 2008).

**Figure 4:**
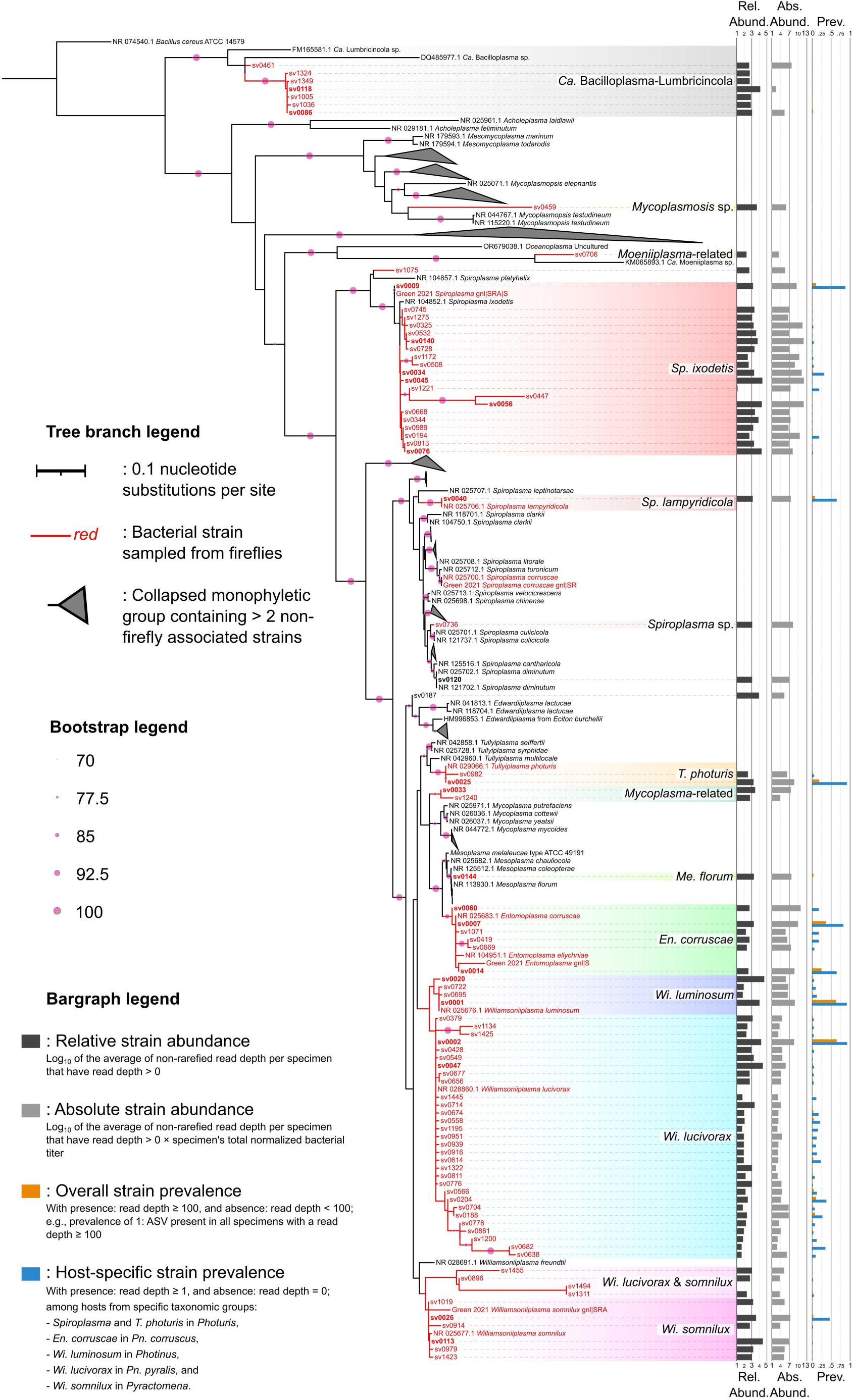
Phylogenetic strain diversity of the *Mollicutes* associated with fireflies. The tree was obtained from a maximum likelihood phylogenetic analysis on the 16S rRNA gene, rooted with a *Bacillus cereus* sequence (NR_074540.1), and contains 230 leaves of which 88 are 16S rRNA amplicon sequences from the present study, four are amplicon sequences from another firefly microbiome study (Green et al. 2021), and four are amplicon sequences from another study on insect-associated mollicutes (Funaro et al. 2011). All the other leaves are full length 16S rRNA gene sequences and were obtained either through NCBI searches to acquire reference sequences of mollicute species or through NCBI BLASTn runs to find the most similar sequences to our amplicon sequences. Red branches contain bacterial ASVs sampled from fireflies (from the present and previous studies), only focusing on our 1,500 most abundant ASVs (SV identifier number lower than 1,501). ASV labels in bold are among the most abundant in our dataset (SV identifier number lower than 151). Gray triangles correspond to collapsed monophyletic groups containing more than two non-firefly associated leaves. Bootstrap values above 70 are depicted with a pink dot on the tree nodes. Genus and species names are from Gupta et al. (2018, 2019). A detailed version of the tree with uncollapsed leaf labels is shown in Figure S9. Bargraphs on the right represent ASV average relative abundance (“Rel. Abund.”) when present (dark gray), average absolute abundance (“Abs. Abund.”) when present (lighter grey), and prevalence (“Prev.”) both across all specimens in our dataset (gold) and within specific firefly groups (blue). Only specimens’ whole body was considered for these calculations.

**Figure 5:**
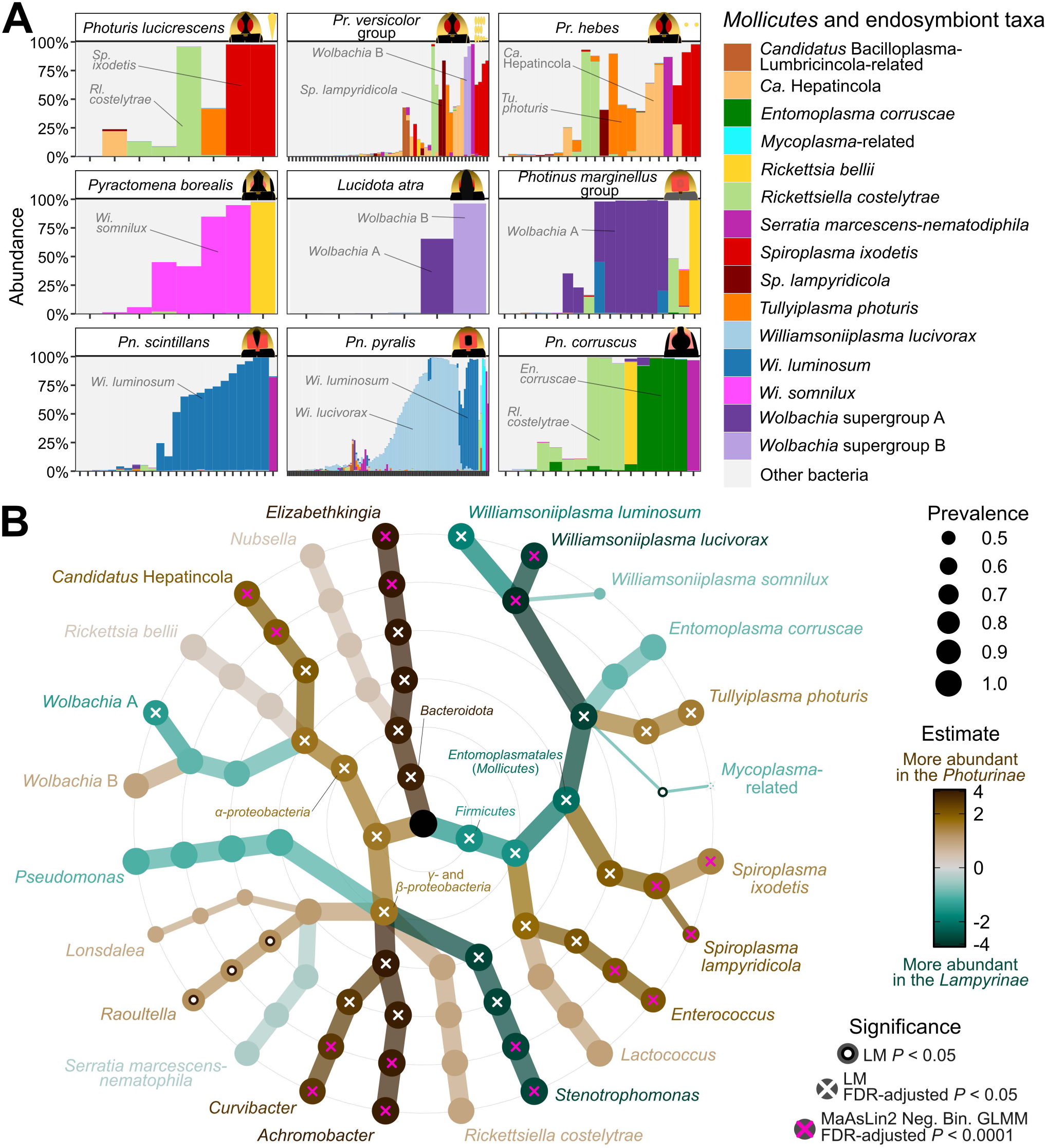
Bacterial composition and differential abundance. **A**. Composition of bacterial taxa across firefly species and species groups with more than three specimens in our dataset post rarefying. Each vertical bar (x-axis) represents one firefly specimen. The relative rarefied abundance (y-axis) of the most abundant mollicutes and putative endosymbionts is shown as different colors whereas the abundance of all other bacteria is in light grey and can be seen in Figure S12. **B**. Differential bacterial abundance at multiple taxonomic levels across adult fireflies. Only firefly species with more than one specimen in our dataset, post rarefying, and only bacterial taxa with rarefied abundance > 20,000 reads in one or more samples were used in this analysis. Moving outwards from the center, light gray rings show phylum, class, order, family, and genus (sensu SILVA database); bacterial “taxa” are found on the outermost ring of the tree. The size of the balls depicts prevalence of a clade across samples (proportion of firefly specimens in which it was detected). Results from three statistical tests comparing bacterial abundances in *Lampyrinae vs. Photurinae* subfamilies are displayed. Following the replacement of zeros by half of the minimum value (to reduce zero-inflation and improve model fitting) and log_2_ transformation, a linear model (“LM”) was run on all bacteria at every taxonomic rank using the *microViz* R package. White circles and crosses on balls indicate significantly differential abundance with unadjusted and false discovery rate (“FDR”)-adjusted, respectively, at *P* < 0.05. Estimates of bacterial differential abundance from this LM are displayed with shades of brown (darker: higher abundance in *Photurinae*) and sea green color (darker: higher abundance in *Lampyrinae*). At both the bacterial genus and taxon levels, differential abundance was additionally tested through *MaAsLin2* multiple generalized linear mixed models (“GLMM”) with negative binomial (“Neg. Bin.”) distribution and controlling for date of capture, geographic origin, and sex. Pink crosses show the bacterial taxa that were found to be significantly differentially abundant at FDR-adjusted *P* < 0.001.

#### 3.b. *Stenotrophomonas* and *Williamsoniiplasma* often dominate lampyrine microbiomes

Following filtering of bacterial taxa with low abundance (keeping only taxa with rarefied abundance > 20,000 reads in one or more samples) in our amplicon sequencing dataset, we found that the *Stenotrophomonas* (*Xanthomonadales*) and *Williamsoniiplasma* (*Mollicutes*) genera were characteristic members of the microbiome of fireflies from the *Lampyrinae* subfamily (Figure 5B), each detected in more than 98% of lampyrine fireflies (colonizing > 57% of lampyrine specimens with more than 1% relative abundance). Compared to fireflies from the *Photurinae* subfamily, lampyrine fireflies were significantly more associated with bacteria from these two genera (MaAsLin2: coef > 3.30, FDR *P*-adj < 0.0001; Table S5). While *Stenotrophomonas* could also be found in *Photuris* fireflies, albeit typically at low abundance, members of the *Williamsoniiplasma* genus were very rare in *Photuris* species (Figure 5A and 5B; Figure S11 and S12), making this bacterial genus nearly endemic to the lampyrine microbiome.

Three *Williamsoniiplasma* taxa were commonly detected in *Photinus* and *Pyractomena* fireflies with differing levels of endemicity (Figure 5A; Figure S11). *Wi. lucivorax* was only found in *Pn. pyralis*, as the most dominant and prevalent bacterium after *Stenotrophomonas*, infecting over 69% of individuals with more than 1% relative abundance, and being significantly more abundant in this firefly species than in any other (MaAsLin2: coef > 4.50, FDR *P*-adj < 0.0001; Figure S11; Table S5). In contrast, *Wi. luminosum* was detected in both *Pn. pyralis* and *Pn. scintillans*, as well as in two individuals from the *Pn. marginellus* group (Figure 5A; Figure S11), thus appearing less host-restricted than *Wi. lucivorax*. Yet, *Wi. luminosum* was significantly associated with *Pn. scintillans* (*vs.* all firefly species but *Pn. marginellus* group: MaAsLin2: coef > 1.55, FDR *P*-adj < 0.05; Figure S11), being found in all *Pn. scintillans* specimens and 64% of *Pn. scintillans* specimens with 1% relative abundance threshold (Figure 5A). Another species from the *Williamsoniiplasma* genus, *Wi. somnilux*, was significantly more abundant in *Pyractomena borealis* than in other firefly genera (MaAsLin2: coef > 8.04, FDR *P*-adj < 0.0001; Table S5) and showed endemicity patterns similar to those of *Wi. lucivorax* (Figure 5A; Figure S11). When using the 1% relative abundance threshold, *Wi. somnilux* was present in five out of eight specimens (62%) from this firefly species, including both adults and juveniles, and was the most abundant bacterial taxon (Figure 5A). We also found that *Pn. corruscus* fireflies were significantly associated with another member of the *Mollicutes*, the *En. corruscae* taxon (MaAsLin2: coef > 4.30, P-adj < 0.001; Figure 5A; Figure S11), being the most abundant bacterial taxon in all *Pn. corruscus* fireflies, found at more than 1% relative abundance in nine out of 16 individuals (56%), and showing patterns of endemicity as well.

In contrast, four other lampyrine species and species groups rarely harbored mollicutes. Instead, in the *Pn. marginellus* group, *Wolbachia* (*Rickettsiales*) from the supergroup A was found to be a dominant bacterium (colonizing > 47% of specimens with more than 1% relative abundance).

#### 3.c. Photurine fireflies commonly harbor a variety of bacteria, including a few mollicutes

Contrasting with the lampyrines, members of the *Photurinae* were significantly more associated with abundant bacteria (filtered taxa with > 20,000 reads in one or more samples) from the *Achromobacter* (*Burkholderiales*), *Ca.* Hepatincola (*Rickettsiales*), *Curvibacter* (*Burkholderiales*), *Elizabethkingia* (*Flavobacteriales*), *Enterococcus* (*Lactobacillales*), and *Spiroplasma* (*Mollicutes*) genera (MaAsLin2: coef > 1.87, FDR *P*-adj < 0.0001; Figure 5B). Despite a lower relative abundance, *Leucobacter* bacteria also appeared significantly associated with photurine fireflies (MaAsLin2: coef = 2.97, FDR *P*-adj < 0.0001; Figure S11 and S12; Table S5). Among the spiroplasmas, two taxa were significantly more abundant in photurine fireflies: *Sp. ixodetis* and *Sp. lampyridicola* (found in 14% and 7% of *Photuris* specimens at 1% relative abundance threshold, respectively) (MaAsLin2: coef > 3.25, FDR *P*-adj < 0.0001). Another mollicute species, *Tullyiplasma photuris*, was abundant in photurine fireflies, though at lower degrees (MaAsLin2: coef = 0.69, FDR *P*-adj < 0.05; Figure 5A; Figure S11), found in 13% of *Photuris* specimens at 1% relative abundance threshold.

#### 3.d. Bacteria positively and negatively co-occur in fireflies

From our co-occurrence analysis, we found that some bacteria, including endosymbionts (e.g., *Rickettsia bellii*, *Wolbachia*) and potential pathogens (e.g., *Serratia marcescens-nematodiphila*), were more likely to be found on their own in fireflies (Figure S13). We also found some positive co-occurrence patterns within firefly species, including *Pseudomonas*, *Stenotrophomonas*, and *Rl. costelytrae* in both *Pn. marginellus* group and *Pn. pyralis*, and *Achromobacter*, *Curvibacter*, and *Elizabethkingia* in *Photuris* fireflies. This suggests that some bacteria are more likely to occur alone whereas others may be part of a consortium in fireflies.

### 4. Within-species variation can be partially explained by the region, season, environment, sex, and developmental stage

#### 4.a. The abundance of some, but not all, bacteria vary across geographic locations

To assess whether geography was a main factor influencing changes in firefly microbiomes, we first used partial Mantel tests on microbiome dissimilarity matrices and geographic distance, while taking into account species’ phylogenetic distances. Through this test, we found a weak positive correlation between pairwise microbiome dissimilarities based on relative bacterial abundance and geographic distance (partial Mantel test controlling for genetic distances: *r* = 0.05, *P* < 0.05; Table S3), indicating that regardless of host species identity, fireflies separated by longer spatial distances tend to harbor more dissimilar microbial communities. Because of substantial intra-specific variation in bacterial density and community composition, we next investigated within species microbiome variation for species that were collected in multiple regions.

At the level of the whole bacterial community, *Pn. corruscus* microbiomes did not appear to differ across regions (PERMANOVA: *R*^2^ = 0.40, *F*_5,10_ = 1.33, *P* = 0.11; Figure S14; Table S3). Nevertheless, and despite a low sample size (*n* = 16) and potential confounding factors (albeit being controlled as random factors in our GLMMs and MaAsLin2 models), we found regional differences in terms of absolute bacterial abundance (Supplementary Text 1), and at the level of individual bacteria (Figure 6A; Table S6). *En. corruscae*, which was a characteristic bacterial taxon of *Pn. corruscus*, may have preferentially been associated with specimens from northern latitudes, since it was less abundant in specimens collected in southwestern Connecticut (swCT) and South New Jersey compared to specimens collected in northwestern CT (nwCT), southeastern New York (seNY), and southwestern Vermont (swVT) (MaAsLin2: coef > 4.57, FDR *P*-adj < 0.01).

**Figure 6:**
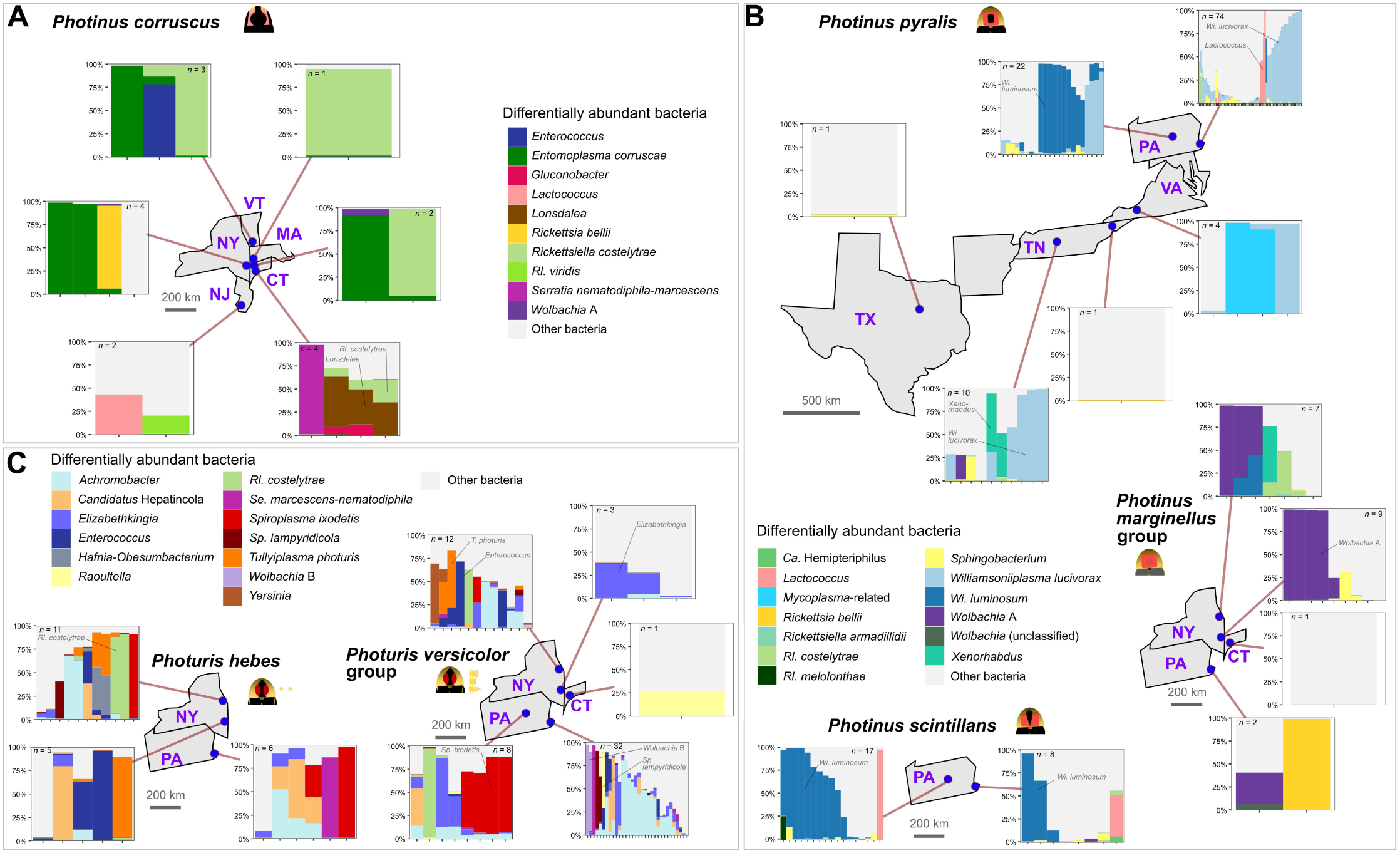
Regional differences in bacterial composition. Bacterial composition, with a focus on the most differentially abundant bacterial taxa across regions. Firefly species are grouped by microbiome similarity, with *Photinus corruscus* in **A**, *Pn. pyralis*, *Pn. marginellus* group, and *Pn. scintillans* in **B**, and *Photuris hebes* and unidentified species from the *Pr. versicolor* group in **C**. Bacterial taxon composition and abundance per species per region was obtained from rarefied read depth. The number of specimens is given within each bar plot. In all panels, “CT”: Connecticut, “PA”: Pennsylvania, “MA”: Massachusetts, “NJ”: New Jersey, “NY”: New York, “TN”: Tennessee, “TX”: Texas, “VA”: Virginia, and “VT”: Vermont US states. A full list of differentially abundant bacterial taxa across regions can be found in Table S6.

*Pn. pyralis* microbiomes showed significant variation in composition across regions (PERMANOVA: *R*^2^ = 0.13, *F*_5,106_ = 3.15, *P* < 0.001; Figure S14; Table S3), despite relatively stable absolute bacterial abundances (emmeans: *z*-ratio < 1.76, *P* > 0.07). The most divergent microbiomes were from *Pn. pyralis* specimens collected in central Pennsylvania (cPA) (pairwise adonis: *R*^2^ > 0.08, *F* > 2.99, FDR *P*-adj. < 0.05; Figure 6B). Likely contributing to the singularity of cPA *Pn. pyralis* microbiomes, *Wi. luminosum* was mostly found in individuals collected in this region (MaAsLin2: coef > 3.64, FDR *P*-adj < 0.05; Table S6), where its prevalence at 1% relative abundance threshold was 68% (64% for *Wi. lucivorax* in these specimens) *vs.* 4% in *Pn. pyralis* from all other regions (71% for *Wi. lucivorax* in these other specimens). For *Wi. lucivorax*, relative abundance was high in specimens collected in several states. *Pn. pyralis* specimens from western Virginia also had a divergent microbiome compared to specimens from other regions, with compositions differing from those of cPA and southeastern PA specimens (pairwise adonis: *R*^2^ = 0.03, *F* = 2.51, FDR *P*-adj. < 0.05), and unusually dominated by bacteria from the *Mycoplasma*-related group (Figure 6B). For both *Pn. marginellus* group and *Pn. scintillans*, we found limited evidence of regional specificities in the microbiome (PERMANOVA: *R*^2^ < 0.17, *F* < 1.67, *P* > 0.11; Figure 6B; Figure S14), although total bacterial density was higher in *Pn. scintillans* collected in cPA compared to sePA (emmeans: *z*-ratio = 2.03, *P* < 0.05).

Regarding the *Photuris* genus, we found strong microbiome differences across regions for the unidentified species of the *Pr. versicolor* group (PERMANOVA: *R*^2^ = 0.13, *F*_4,51_ = 1.83, *P* < 0.001; Figure S14; Table S3), but not for *Pr. hebes* (PERMANOVA: *R*^2^ = 0.11, *F*_2,19_ = 1.18, *P* = 0.24). Members of the bacterial taxon *Sp. ixodetis* were common and abundant in specimens collected in cPA, but were very rare in specimens from other regions (MaAsLin2: coef > 5.58, FDR *P*-adj < 0.001; Table S6). We also identified regional specificities among other common bacterial associates of *Photuris* fireflies, with *T. photuris* being more abundant in northeastern NY (neNY) and seNY compared to sePA for *Pr. hebes* (MaAsLin2: coef > 5.88, FDR *P*-adj < 0.0001), and neNY compared to sePA in unidentified species from the *Pr. versicolor* group (MaAsLin2: coef > 3.53, FDR *P*-adj < 0.05).

#### 4.b. Seasonal, but not annual, factors influence within-species microbiome variation

We tested whether seasonality, which is linked to firefly adult age, influenced firefly microbiome. When focusing on all species in our dataset, we found that bacterial density varied throughout day of year (GLMM Wald test: *Χ*^2^ = 124.38, *P* < 0.0001; Table S3), but this appeared to be the case mostly for some firefly species. Several species, including *Pn. scintillans*, unidentified *Pr. versicolor* group, and *Py. borealis*, tended to have higher bacterial titers later in the season (emmeans: *z*-ratio > 2.63, *P* < 0.05). We also found that bacterial composition varied throughout the year and across species (PERMANOVA with species and regions as covariates, focusing on the interaction between day of year and species: *R*^2^ = 0.05, *F*_14,186_ = 1.20, *P* < 0.05). We next focused our analyses only on *Pn. corruscus* specimens given their long adult lifespan (up to 10 months *vs.* less than three months usually for other species) and since we collected this species throughout the year.

*Pn. corruscus* specimens collected in the months of April, May, and June had higher bacterial titers than those collected in November (emmeans: *z*-ratio > 3.55, *P* < 0.01; Figure 7A; Table S3). When looking at individual bacterial taxa, we found that specimens collected in May and April were typically associated with high to very high relative abundance of *En. corruscae*, but this bacterial taxon was barely detectable in *Pn. corruscus* specimens collected in February and November (April and May *vs.* February and November, MaAsLin2: coef > 4.89, FDR *P*-adj < 0.01; Figure 7B; Table S7). Supporting these results across months, we found a strong association between the collection day of year used as a quadratic continuous predictor in our LMMs and both total bacterial titers (GLMM Wald tests: *Χ*^2^ > 35.81, *P* < 0.0001), and abundance of *En. corruscae* (LMM Wald tests: *Χ*^2^ > 18.47, *P* < 0.0001), in *Pn. corruscus*, with a peak in late May (Figure 7A and 7B). Thanks to citizen science data collected from iNaturalist (iNaturalist community 2025), we found that during the years and in the states of collection, the reproductive activity of *Pn. corruscus* spanned from early April to mid/late June, peaking during the month of May (Figure 7C). This corresponds to the reproductive seasons observed for this species in other states and years (Faust 2012; Rooney and Lewis 2000; Smedley et al. 2017), and implies a potential link between *En. corruscae* abundance and either its host reproduction or seasonal increased food intake (Rooney and Lewis 2000).

**Figure 7:**
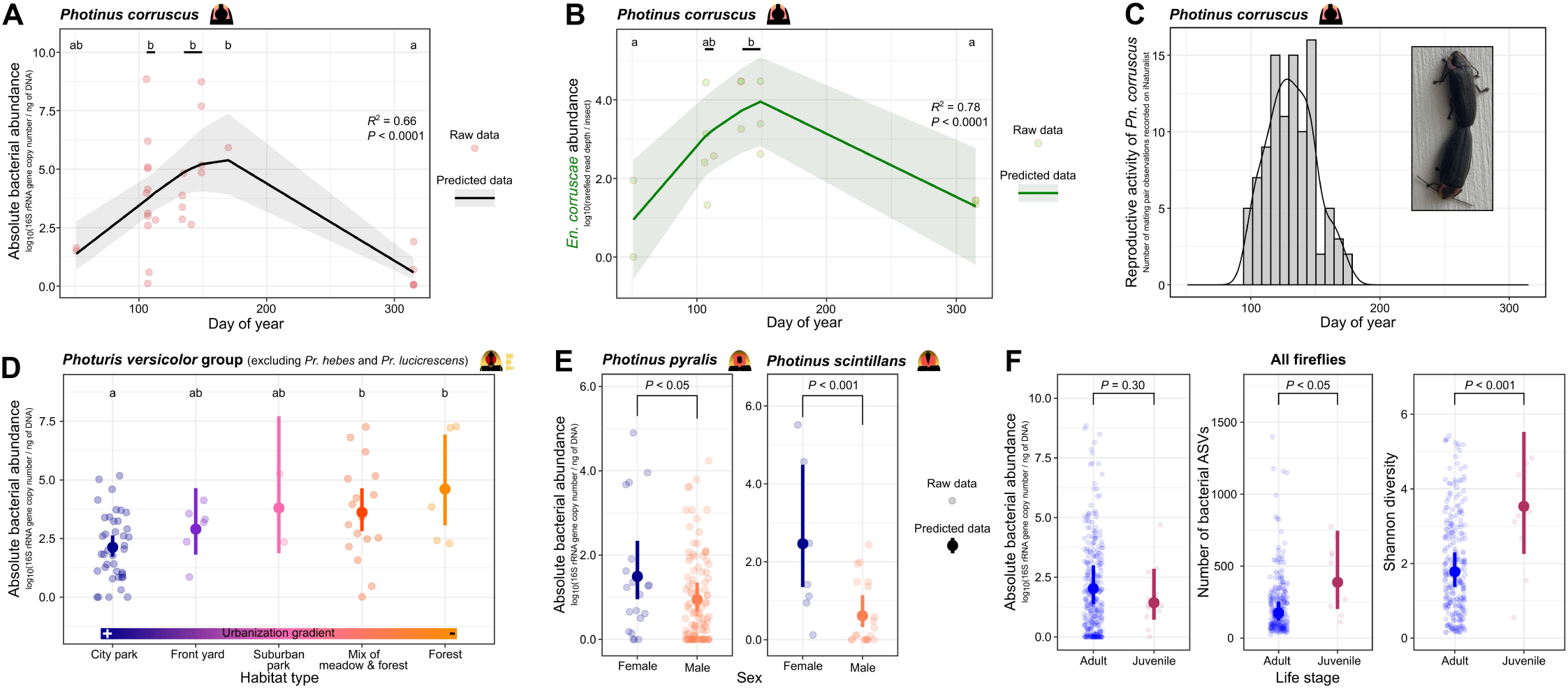
Seasonal, environmental, and life history factors influence the microbiome of fireflies. **A**. Normalized absolute bacterial abundance in *Photinus corruscus* peaks at the end of spring. A generalized linear mixed model (GLMM) was used to test whether bacterial titer is influenced by day of year, used as a quadratic term, in *Pn. corruscus*, when controlling for sex, latitude, year of capture, and whether the specimen was dissected or not. Change in bacterial titers over the months for this species was also tested with least-square means (emmeans) post hoc test on a GLMM controlling for the same factors. Significant differences between February, April, May, June, and November are shown with lowercase English letters. **B**. Abundance of *Entomoplasma corruscae* in *Pn. corruscus* also peaks at the end of spring. *En. corruscae* abundance is expressed as the log_10_ of rarefied read depth. A GLMM was used to test whether *En. corruscae* abundance is influenced by day of year, used as a quadratic term, in *Pn. corruscus*, when controlling for sex, region of capture, year of capture, and specimen identifier. Lowercase English letters indicate significant differences between months, as measured in the previous panel. **C**. Reproductive activity in *Pn. corruscus* also peaks in the spring. The bars depict the number of mating pairs observed per week on iNaturalist for this species in Connecticut, New Jersey, New York, Pennsylvania, and Vermont, in 2021, 2022, and 2023. The photograph of a type 1 copulation event (Wing, 1984) was taken in New York state in 2021 by Benoît Béchade. **D**. For unidentified species from the *Photuris versicolor* group, the highest bacterial densities are found in specimens from the least urbanized habitats. A GLMM was used to test whether bacterial titer is influenced by habitat type in these species, when controlling for sex, latitude, year and time of capture, day of year, specimen identifier, and whether the specimen was dissected or not. Lowercase English letters indicate significant differences across habitat types. **E**. Bacterial absolute abundance is higher in females than in males of *Pn. pyralis* and *Pn. scintillans*. A GLMM was used to test whether bacterial titer is different between females and males in firefly species (i.e., through the interaction term sex:species), when controlling for the factors as in the previous panel. **F**. Firefly juveniles harbor a higher bacterial diversity than adults, but the two life stages do not differ in their absolute bacterial abundance. GLMMs were used to test whether bacterial titer, bacterial richness, and the bacterial Shannon diversity index are different between juveniles (including larvae and pupae) and adults, using firefly genus as a covariate and controlling for region, day of year, and year of capture. For panels A, D, E, and F, bacterial titer is expressed as the log_10_ of the number of 16S rRNA gene copies in a sample, measured through qPCR, minus the number of 16S rRNA gene copy in the DNA extraction blank sample from the same batch, and then divided by the total DNA concentration in the sample, which was measured via Qubit. For all panels except C, predicted data (curve for A and B and balls for D, E, and F) with 95% confidence interval (grey shade for A, B, and C, and error bars for D, E, and F) were obtained from GLMMs.

To assess microbiome turnover across years, we collected *Pn. pyralis* specimens in the same location, at approximately the same time of year, for three consecutive years. We found that the overall microbiome composition of *Pn. pyralis* changed only marginally over the years (PERMANOVA: *R*^2^ = 0.06, *F*_2,53_ = 1.76, *P* = 0.051; Table S3). However, at the level of individual bacterial taxa, while the abundance of *Wi. lucivorax* remained stable from 2021 to 2023, that of *T. photuris* and *Wi. luminosum* increased and that of *Pseudoclavibacter* and *Tumebacillus* dropped (MaAsLin2: coef < -1.96, FDR *P*-adj < 0.05; Figure S15; Table S7). This shows that the overall composition of bacteria associated with fireflies may not drastically change over the years, although the abundance of a few bacteria may fluctuate.

#### 4.c. The microbiome of *Photuris versicolor* group fireflies changes across habitat types

The interaction between species and habitat types (with different urbanization levels) significantly affected both bacterial composition and density (PERMANOVA with species and regions as covariates: *R*^2^ = 0.05, *F*_14,235_ = 1.30, *P* < 0.001; GLMM Wald test: *Χ*^2^ = 143.6, *P* < 0.0001; Table S3), indicating that different species’ microbiomes vary in their response to divergent habitat types. In unidentified species from the *Pr. versicolor* group, we found that absolute bacterial abundance was significantly lower for specimens collected in the city than in the forest or in a mix of forest and meadow (emmeans: *z*-ratio > 3.17, *P* < 0.05; Figure 7D). Through MaAsLin2 multiple GLMMs controlling for potential confounding factors, we found that *Rickettsiella costelytrae* may have been a key influence on bacterial titers over the urbanization gradient since it was more abundant in specimens from the forest than in the city (MaAsLin2: coef = 5.72, FDR *P*-adj < 0.01; Figure S16; Table S7). These results show that the type of habitat used by *Photuris* fireflies may influence the abundance and composition of their associated bacteria.

#### 4.d. *Photinus* females harbor a denser bacterial community than males

Using qPCR, we found that females of both *Pn. pyralis* and *Pn. scintillans* had a significantly higher bacterial titer than their male conspecifics (emmeans: *z*-ratio > 2.03, *P* < 0.05; Figure 7E; Table S3). In contrast, in *Photinus* species, the Evar index of evenness was higher for males compared to females (emmeans: *z*-ratio = 3.75, *P* < 0.01; Figure S17), suggesting that bacterial strains in male microbiomes were represented in more equal proportions, while female microbiomes tended to be dominated by fewer bacteria. Nevertheless, firefly microbiome composition did not significantly differ between sexes, with the only exception of *Pr. hebes* (pairwise adonis: *R*^2^ = 0.09, *F* = 1.97, FDR *P*-adj < 0.05).

#### 4.e. Juveniles harbor a more diverse bacterial community than adults

We collected a few larvae from three different firefly genera, and pupae from the *Pyractomena* genus. When comparing adults to juveniles (i.e., including both larvae and pupae) from all firefly species, while we found no difference in terms of absolute bacterial abundance (GLMM Wald test: *Χ*^2^ = 1.08, *P* = 0.30; Figure 7F; Table S3), there was a significant difference in their microbiome composition (PERMANOVA with life stage nested within species within genus within subfamily: *R*^2^ = 0.01, *F*_3,261_ = 1.35, *P* < 0.05), a higher bacterial richness (GLMM Wald test on number of ASVs with firefly genus used as a covariate: *Χ*^2^ = 4.99, *P* < 0.05), and a higher bacterial diversity (GLMM Wald test on Shannon diversity with firefly genus used as a covariate: *Χ*^2^ = 7.40, *P* < 0.001) in juveniles compared to adults. These results appear to be driven by the difference between larvae and adults (Figure S18; Supplementary Text 1).

The single larva from the *Photinus* genus, identified through DNA barcoding as from the *Pn. consanguineus* group (Figure S1), did not appear to have a different bacterial composition compared to adults, and did not harbor any *Williamsoniiplasma* spp. (Figure S19). In contrast, *Pr. versicolor* group larvae had a different bacterial composition compared to adults (PERMANOVA: *R*^2^ = 0.03, *F*_1,84_ = 2.22, *P* < 0.01; Table S3), including a significantly higher abundance of unidentified *Rhizobiales* (MaAsLin2: coef = 8.92, FDR *P*-adj < 0.05; Table S7). Notably, one larva specimen from this species group had a higher bacterial titer compared to the two other larvae and harbored *Sp. ixodetis* with high relative abundance. All three *Photuris* larvae analyzed hosted *Leucobacter* in high relative abundance, which was also commonly found in adults (Figure S11 and S12).

Juveniles from the *Py. borealis* species did not have a significantly different bacterial community as a whole when compared to adult males from the same species (PERMANOVA: *R*^2^ = 0.25, *F*_1,6_ = 1.99, *P* = 0.11; Figure S19; Table S3). Nevertheless, and despite our low sample size, significant differences in the abundance of single bacteria across larvae, pupae, and adult males could be noted. The mollicute *Wi. somnilux*, which was commonly detected in *Py. Borealis* fireflies, associated with adults and larvae at similar abundances (MaAsLin2: coef = 0.14, FDR *P*-adj = 1; Figure S20; Table S7), but was present in lower abundance in pupae (MaAsLin2: coef = 7.92, FDR *P*-adj < 0.01). Other bacterial genera, including *Rhodococcus*, had a significantly higher abundance in *Pyractomena* juveniles compared with adult males (MaAsLin2: coef > 1.99, FDR *P*-adj < 0.05; Figure S19). These results indicate that juveniles harbor a more diverse microbiome, and some of the most prevalent and abundant adult-associated bacteria are also abundant in larvae and detected in pupae.

### 5. The tissue localization of bacteria varies across firefly species

We investigated preferential colonization of bacteria in specific tissues of three firefly species and species groups through dissections and tissue separations (Table S8). We found a significant difference in overall bacterial composition across firefly tissues, but only for unidentified species from the *Pr. versicolor* group, between their lantern and gut tissues (pairwise adonis: *R*^2^ = 0.12, *F* = 2.36, FDR *P*-adj < 0.01; Figure S21; Table S3), indicative of major tissue differences only in *Photuris* fireflies.

#### 5.a. *Pn. corruscus* tissues do not differ in bacterial titer but have a few differentially abundant bacterial taxa

Bacterial densities were relatively high throughout the body of *Pn. corruscus* (Figure 8A). In terms of composition, *En. corruscae* commonly colonized all the main tissues of both males and females, suggesting that most *Pn. corruscus* tissues are colonized by bacteria in high abundance, including prevalent mollicutes.

**Figure 8:**
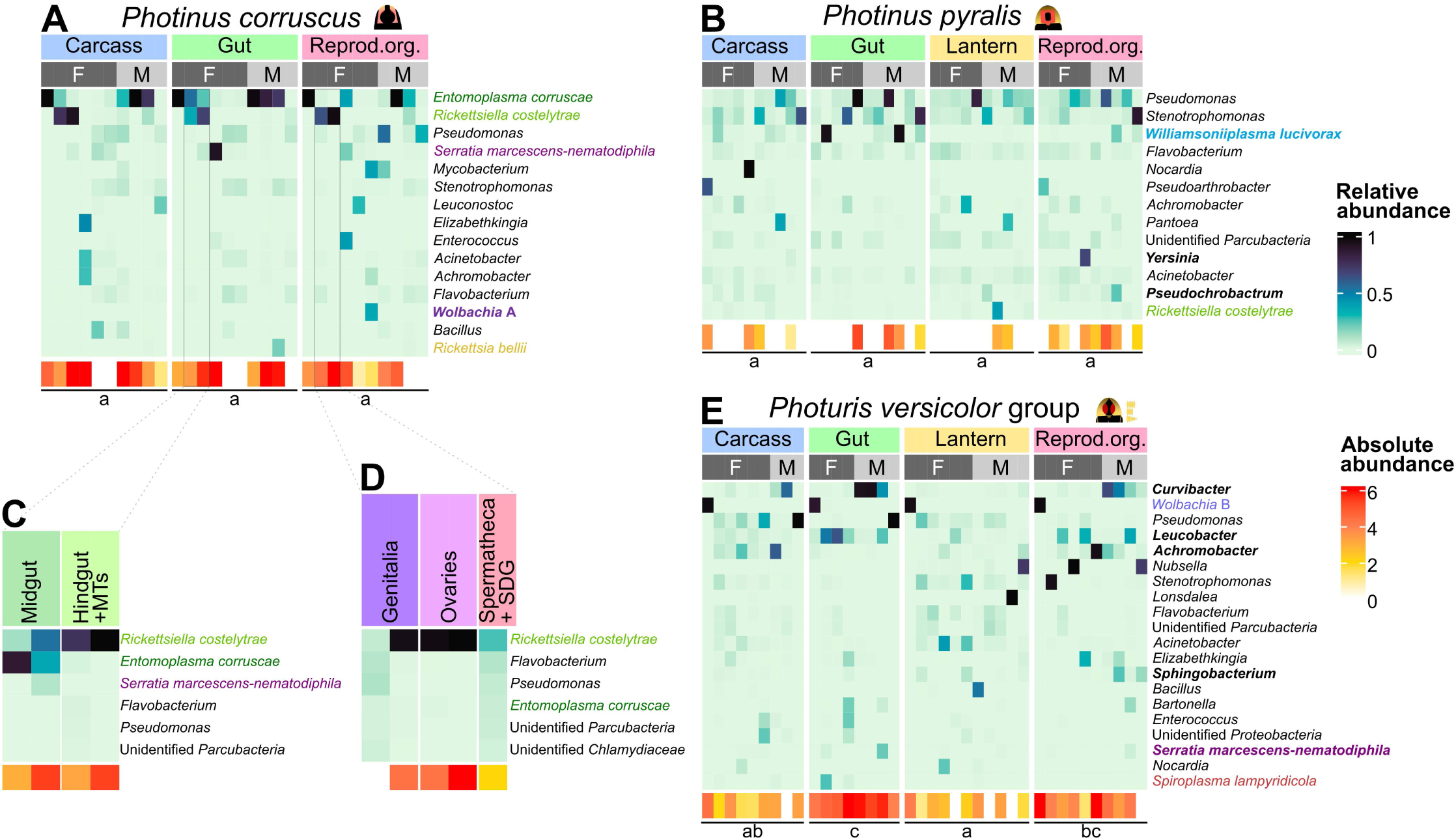
Microbiome changes across tissues in three firefly species. **A**, **B**, and **E**. Bacterial community composition and abundance across the carcass, gut, reproductive organs (“Reprod. Org.”), and lantern in *Photinus corruscus*, *Pn. pyralis*, and unidentified species from the *Photuris versicolor* group. Blue shaded heatmaps show the relative abundance of the most abundant bacterial taxa for each tissue per specimen. The abundance was calculated from non-rarefied, proportional (i.e., compositional) transformations of read depth relative abundance. Bacterial taxa with bold fonts are significantly differentially abundant between at least two tissues in the species, as tested through *MaAsLin2* multiple generalized linear mixed models (GLMMs), using the cumulative sum scaling normalization method, controlling for geographic location and date of sampling, and using the FDR-adjusted *P* < 0.05 as significance threshold. All the differentially abundant bacterial taxa across tissues can be found in our Table S9. Red shaded heatmaps show non-normalized absolute bacterial abundance, expressed as the log_10_ of the number of 16S rRNA gene copies in a sample, measured through qPCR, minus the number of 16S rRNA gene copy in the dissection blank sample, minus the number of 16S rRNA gene copy in the DNA extraction blank sample from the same batch. A linear mixed model (LMM) was used to compare bacterial titer across tissues in these species, by using the species:tissue interaction term as the fixed effect, controlling for sex, latitude, day of year, year of capture, and temperature. For each firefly species, lowercase English letters indicate significant difference in absolute bacterial titers across tissues, as obtained from emmeans post-hoc tests on the LMM. **C** and **D**. Total absolute bacterial abundance and relative abundance of the six most abundant bacteria in separated gut compartments and reproductive organs of two *Pn. corruscus* female specimens. “MTs”: Malpighian tubules and “SDG”: spermatophore-digesting gland. More detailed heatmaps and dissection pictures can be found in Figure S22–S24, and dissection information can be found in Table S8.

Two females had their gut segments and reproductive organs separated through dissection. The mollicute *En. corruscae* was not found in ovaries of these females, but was present in other reproductive organs, and was very abundant in the midgut (Figure 8A, 8C, and 8D; Figure S22). In contrast, *Rl. costelytrae* was very abundant in *Pn. corruscus*’ carcass, hindgut, and ovaries and was also detected in the genitalia, spermatheca and spermatophore-digesting gland, and midgut of these two individuals, matching infection patterns typical of symbionts residing inside host tissues (i.e., endosymbionts; Perlmutter and Bordenstein 2020). Mollicutes and endosymbionts may thus slightly differ in their tissue localization in *Pn. corruscus*, *En. corruscae* being mainly found in the midgut but also infecting reproductive organs, and endosymbionts such as *Rickettsiella* colonizing most tissues, including ovaries.

#### 5.b. Bacteria variably colonize the tissues of *Pn. pyralis* but with a potential primary gut localization of *Williamsoniiplasma*

While there were no significant differences in bacterial titers across *Pn. pyralis* dissected tissues (emmeans: *t*-ratio < 1.93, *P* > 0.22; Figure 8B), the gut and reproductive organs could occasionally be infected with high titers. When present, *Wi. lucivorax* was found to be common and relatively abundant in the gut, and while it was sometimes detected in the carcass and reproductive organs of males, it appeared virtually absent from the lantern (MaAsLin2: coef > 3.79, FDR *P*-adj < 0.05; Figure 8B; Figure S23; Table S9). Despite being much less abundant overall than its sister species, the abundance of *Wi. luminosum* was also reduced in the lantern compared to the gut (MaAsLin2: coef = 2.40, FDR *P*-adj < 0.001). This suggests that, as for *Pn. corruscus*, mollicutes may have a primary gut localization in *Pn. pyralis*, but with lower bacterial titers overall in this firefly species.

#### 5.c. Specimens from the *Photuris versicolor* group harbor high density gut bacteria

While bacterial density in unidentified species from the *Pr. versicolor* group whole body was relatively high (Figure 1C; Figure S3), it peaked in the gut, especially compared to the lantern and carcass (emmeans: *t*-ratio > 2.85, *P* < 0.05; Figure 8E; Figure S24; Table S3). All individuals of these species had a consistently high bacterial titer in the gut, regardless of sex, collection date, and location. Reproductive organs also had significantly higher bacterial titers than the lantern (emmeans: *t*-ratio = 2.71, *P* < 0.05).

When separating the unidentified *Pr. versicolor* group specimens by sex, we found that bacteria from the *Curvibacter* genus dominated the microbiome of male guts and reproductive organs compared to female guts and other male tissues (male gut vs. all other tissues except male reproductive organs: MaAsLin2: coef > 4.19, FDR *P*-adj < 0.05; Figure 8E; Table S9). Otherwise, the gut of both females and males included a high abundance of *Leucobacter* in the gut and reproductive organs compared to the carcass and the lantern (MaAsLin2: coef > 2.93, FDR *P*-adj < 0.05). Notably, one female harbored *Wolbachia* from the supergroup B in all its tissues and another one had a high abundance of *Sp. lampyridicola* in its gut (Figure 8E). *Photuris* fireflies thus harbor high density gut bacteria with compositions that seem to differ between males and females.

## Discussion

We studied the factors shaping firefly microbiomes through a comprehensive characterization of bacterial communities in several Northeastern US firefly species. Echoing previous research (Green et al. 2021; Hackett et al. 1992), we detected a high variation in bacterial composition and abundance within firefly species (Figure 1C and 5A; Figure S3, S12, and S19). However, we also reveal that the bacterial community of most firefly species is unique (Figure 3C–3F), implying that each host species preferentially associates with its own set of bacteria. This high species-specificity combined with intraspecific variability could be explained by the interplay between stochastic and deterministic mechanisms. This was supported by our analysis of normalized stochasticity ratios (NST), showing that either mollicutes or endosymbionts (or both in some *Photuris* species) were acquired by fireflies through deterministic factors, but when these symbionts were not included in the microbiome the rest of these firefly-associated bacterial communities were mainly structured through stochastic forces (Figure 2). These results highlight the importance of partitioning microbiome analyses based on microbial incidence (e.g., *Mollicutes*) and known functional groups (e.g., endosymbionts), as different ecological processes dictate their acquisition and maintenance by hosts.

### Ecological mechanisms structuring firefly microbiota

Mollicutes and some endosymbionts may be acquired and maintained through deterministic forces, such as habitat/host filtering mechanisms, which include some component of the host physiology favoring the acquisition, transmission, and establishment of only specific microbes (Mazel et al. 2018; Michalik and Szklarzewicz 2025; Ohbayashi et al. 2015; Sachs et al. 2004), and bacteria-driven exclusion or facilitation of the establishment of other bacteria (Alizon et al. 2013; Zélé et al. 2018). Other bacteria composing firefly microbiomes may largely be free-living bacteria acquired from the environment, and their acquisition may depend on stochastic mechanisms, such as random exposure events experienced individually by each firefly (Jones et al. 2022; Zhou and Ning 2017). We further discuss these mechanisms and how they contribute to structuring the firefly microbiome below.

First, some of our results support the idea that variability in firefly microbiomes is due to stochastic community assembly mechanisms, including the random acquisition of various free-living microbes from the environment, as suggested before (Green et al. 2021). For example, we found that, for several firefly species, the prevalence of individual bacterial taxa changed across specimens collected in different geographic locations (e.g., *Lactococcus* in *Photinus* species; Figure 6A–6B). A similar variation was observed in *Photuris* microbiomes across habitats with different levels of urbanization (Figure 7D; Figure S16). Such intraspecific variability of microbiome composition has been described in a wide range of host groups like butterflies or mammals (Ley et al. 2008; Ravenscraft et al. 2019). Acquisition of several bacteria from environmental pools, which differ in their composition across geographic locations and habitat types (Hosen et al. 2017; Malard et al. 2019; Martiny et al. 2006; Zheng et al. 2024), thus appears probable in fireflies.

Second, deterministic habitat filtering mechanisms imposed by host physiological conditions are likely to participate in shaping the establishment of characteristic members of the bacterial communities. Supporting this, we found that, despite being collected at the same locations, times of day, and seasons, sympatric firefly species (Faust 2017; Lloyd 2018; Martin et al. 2024; Stanger-Hall and Lloyd 2015) did not share similar microbiomes, whether they were closely related or not (Figure 3A–3F). As previously pointed out by Hackett et al. (1992), even fireflies from the *Photuris versicolor* group, which consume *Photinus* and *Pyractomena* fireflies, did not harbor any of the *Williamsoniiplasma* bacteria characteristic of their prey (Figure 5A and 5B). In addition, several firefly-associated bacterial taxa from the *Mollicutes* may be endemic to fireflies since they have only been found in firefly tissues to date (Figure 4; Figure S8 and S11). These results are indicative of host restriction, suggesting that fireflies are not entirely permeable to the microbes they are exposed to, and their microbiome composition cannot be fully explained by passive colonization of environmental bacteria. Unknown habitat filtering mechanisms within the host are thus likely to contribute to the assembly of firefly microbiomes, as is the case in other organisms (Ohbayashi et al. 2015; Sachs et al. 2004).

Vertical transmission can be considered as a special case of deterministic habitat/host filtering mechanism, where only specific symbionts are transmitted from parents to offspring and conserved by the host line throughout generations. Many insects associating with lineage-specific, primary symbionts have been shown to rely on strict vertical transmission to preserve key beneficial functions from the symbiont (Baumann 2005; McCutcheon et al. 2009; Sacchi et al. 2000). Yet, several endosymbionts that are common across arthropods, such as *Rickettsia*, *Rickettsiella*, *Spiroplasma*, and *Wolbachia* practice a mix of vertical and horizontal transmission with no strict host specificity (Gu et al. 2023; Liu et al. 2024; Michalik and Szklarzewicz 2025; Perreau and Moran 2022). Here, we confirmed the endosymbiotic nature of *Rickettsiella* and *Wolbachia* in fireflies, which infected several tissues, including reproductive organs (Figure 8A– 8E). While our NST analysis suggests that the acquisition of these endosymbionts was deterministic in *Pn. marginellus* group and unidentified species from the *Pr. versicolor* group (Figure 2), these endosymbionts appeared with a more random distribution in other firefly species (Figure 5A), suggesting a stochastic influence, perhaps conditioned by horizontal jumps across hosts. Although we detected endosymbionts and mollicutes in female reproductive organs (Figure 8A), it is still unclear whether they are vertically transmitted to firefly offspring.

Third, bacteria-driven exclusion or facilitation is another deterministic process that could structure firefly-associated bacterial communities. In insect microbiomes, some bacteria have to compete to establish in a host (Itoh et al. 2019), while others need other bacterial species to be present (Carpenter et al. 2021). We found that when they were detected in fireflies, *Rickettsia* and *Wolbachia* were each likely to dominate bacterial communities and rarely co-occurred with other bacteria (Figure S13). This result could be a consequence of potential competitive exclusion driven by these known endosymbionts (Caragata et al. 2013). However, we cannot reject the possibility that these endosymbionts were present at such high densities in firefly tissues that the sequencing depth was not high enough to detect other co-occurring bacteria (Gofton et al. 2015). We note that *Rickettsia bellii* and *Wolbachia* supergroup B usually had a very high absolute bacterial abundance in specimens (Figure S19 and S24), supporting this latter possibility. We also found patterns of positive co-occurrence among bacteria associated with a few firefly species, which could suggest facilitation. Chief among them is the *Pseudomonas*-*Stenotrophomonas* duo found in several *Photinus* fireflies (Figure S13), also known to cooperate during infections of vertebrates (McDaniel et al. 2023; 2020). In *Photuris* fireflies, *Curvibacter*, *Achromobacter*, and *Elizabethkingia* tend to co-occur in the reproductive organs (Figure 8E) and in the gut (Green et al. 2021), suggesting acquisition of this trio either from sexual transmission or from food intake. Future work should investigate whether these bacteria are acquired together by chance (Carpenter et al. 2021), or truly facilitate each other’s establishment in firefly hosts (Halliday et al. 2024; Zélé et al. 2018).

It thus appears that, as in many animals (Chandler et al. 2011; Furman et al. 2020; Hendrycks et al. 2025; Oliphant et al. 2019), fireflies acquire their microbiomes through a combination of stochastic and deterministic elements. Mechanisms involved likely include the acquisition of environmental bacteria, habitat filtering mechanisms—including potential reliable transmission modes, partner choice processes to select specific bacterial strains, and/or parasitic associations where parasites circumvent barriers to establishment—and possible competition and facilitation among bacterial strains. Future studies should test the contribution of each of these stochastic and deterministic mechanisms in structuring firefly microbiota.

### Which factors influence the association with bacterial symbionts?

As a group, fireflies offer diverse ecologies across species and throughout their lifetime. We investigated the influence of several firefly traits, such as sex, dietary ecology, and life stage, on the composition and abundance of housed bacteria. Notably, we did not find convincing evidence directly linking adult bioluminescence to bacterial symbiosis, since bioluminescent firefly species had similar bacterial loads and composition as non-bioluminescent ones (Figure S6) and the lantern of two common species housed either low or variable bacterial density with seemingly random bacterial composition, except in the rare occasion of infections with an endosymbiont colonizing all tissues including the lantern (Figure 8B and 8E; Figure S23 and S24).

Interspecific variation in the animal microbiome is often influenced by diet (Degregori et al. 2024; Karasov and Douglas 2013). Here, we found that bacterial density and composition changed between fireflies known to eat versus those supposed not to eat as adults (Figure S6). However, since diet and phylogeny are confounded in our study, this result was not well supported when accounting for host phylogenetic relationship, which was responsible for a large portion of microbiome (dis)similarity among firefly species (Figure 3A–3C; Figure S5). Notable microbiome divergences observed between closely related firefly species could, nevertheless, be attributed to species’ differing ecology, perhaps including dietary differences. For example, *Pn. pyralis* and *Pn. corruscus*, which are two closely related species (Catalan et al. 2022) with overlapping geographic range—the former not seemingly eating as adults and having a short adult lifespan from May to August, and the latter consuming plant sap and nectar and living from November to May as adults—harbored divergent bacterial densities and compositions (Figure 1C and 3F). In-depth dietary analyses should be performed to formally address the contribution of adult diet to microbial abundance and composition in fireflies, while controlling for host relatedness.

Our results suggest that some of the intraspecific variation in bacterial abundance in *Pn. corruscus* could be explained by seasonal effects. We found that the abundance of the mollicute *Entomoplasma corruscae* associated with these fireflies increased in the spring (Figure 7B), when reproductive activity peaks for this firefly species in the Northeastern US (Figure 7C). Given that this bacterium was sometimes abundant in the midgut, but could also be detected in the reproductive organs and carcass (Figure 8A and 8D; Figure S22), several possible modes of acquisition may promote the proliferation of *En. corruscae* in the spring. First, *Pn. corruscus* fireflies may drastically increase food intake during the reproductive season (Rooney and Lewis 2000), hence acquiring large quantities of this bacterium from their food sources, thus implying an environmental acquisition mode. But it is also possible that *En. corruscae* is vertically transmitted from parents to offspring, although this was not supported in a recent study analyzing the bacterial composition of six eggs (Green et al. 2021). Finally, as seen in the association between aphids and three facultative endosymbionts (Moran and Dunbar 2006), *En. corruscae* could potentially be sexually transmitted (i.e., horizontal transmission), and its abundance in adults could be directly proportional to the number of copulation events they experience.

As holometabolous insects, fireflies have two dissociated, active life stages with drastically divergent ecologies, translating into distinct periods of exposure to, and interaction with, microbes (Hammer and Moran 2019; Manthey et al. 2023): one at the larval stage, after eggs hatch, and one at the adult stage, following pupation. While the ecology, physiology, and behaviors of adult fireflies have been well documented, earlier developmental stages (e.g., eggs, larvae, pre-pupae, and pupae) remain understudied, largely due to the cryptic lifestyles of these early stages (e.g., soil dwelling) and difficulties to rear them throughout their development (Riley et al. 2021). However, it is crucial to study larval microbiomes, since fireflies spend most of their life in this stage. Here, we found that firefly larvae from the Northeastern US harbored a more diverse bacterial community compared to adults (Figure 7F; Figure S18), with diversity index values and a few bacterial genera similar to those from a study on the gut microbiota of distantly related *Aquatica leii* firefly larvae from China (Zhao et al. 2023). When focusing on key bacterial groups in fireflies, although our single *Pn. consanguineus* group larva did not harbor any mollicute in high abundance, we found that all five late stage *Py. borealis* larvae, sampled in different locations (northern Delaware, eastern New York, southeastern Pennsylvania), harbored moderate to high abundance of the mollicute *Wi. somnilux* (Figure S19 and S20). While the high diversity of larva-associated bacterial communities could easily be explained by environmental acquisition of diverse bacteria from the microbe-rich habitats firefly larvae occur in, such as soil (Torsvik et al. 1990), the presence of the same mollicute species in *Py. borealis* larvae collected from locations separated by hundreds of kilometers further supports our finding that deterministic mechanisms promote the acquisition and persistence of species-specific mollicutes in fireflies. However, it is still unknown whether this mollicute species is acquired from the specific environment of *Py. borealis* larvae. Transmission experiments should be performed in this firefly species to determine the bacterial mode of transmission.

Another mystery lies in the fate of symbionts through metamorphosis to adulthood in holometabolous insects. Here, we found that one *Py. borealis* pupa (which was approaching emergence) out of three had detectable, yet low, levels of *Wi. somnilux* (Figure S19 and S20). This bacterium has originally been sampled multiple times in the pupal gut of *Py. angulata* from Maryland (Hackett et al. 1992; Williamson et al. 1990). It thus appears that, at least in *Pyractomena* fireflies, larvae, pupae, and adults associate with species-specific mollicutes, which occasionally appear to persist in the gut during pupation. A drastic reduction in abundance of larva-associated microbes is common during the metamorphosis of holometabolous insects (Hammer and Moran 2019), but the retention of symbionts through pupation has been demonstrated in a few insects before. For example, the fruit fly *Drosophila melanogaster* can harbor high abundance of the mollicute *Spiroplasma* in the gut throughout its lifetime, including during pupation (Anbutsu and Fukatsu 2003). The possible retention of species-specific mollicutes through pupation is yet another indication of the tight, non-random association between these bacteria and their firefly hosts.

### Insights into the evolutionary history of firefly-associated mollicutes

While the transmission mode of mollicutes in fireflies is still uncertain, we found support that these bacteria non-randomly associate with firefly hosts. Although this is not a prerequisite for being a symbiont specialist (Bourguignon et al. 2018; Cabuslay et al. 2024; Koch et al. 2013), several ancient symbionts are so dependent on their host that they have co-diversified with it (García-Lozano et al. 2024; Hu et al. 2023; Kikuchi et al. 2009; Lo et al. 2003; Zhang et al. 2023), forming patterns of co-phylogeny where the symbiont phylogeny tightly matches that of their host. Our phylogenetic analyses (Figure 4; Figure S9), along with previous phylogenomic results (Gupta et al. 2019; Lo et al. 2018), suggest limited evidence for co-cladogenesis between the *Mollicutes* and fireflies, since mollicutes infecting closely related firefly species are rarely each others’ closest relatives. For example, the closely related fireflies *Pn. corruscus* and *Pn. pyralis* (Catalan et al. 2022) commonly harbored *En. corruscae* and *Wi. lucivorax*, respectively, classified in two distinct bacterial genera (Gupta et al. 2019). These two bacterial species are phylogenetically separated by other bacterial genera with very different lifestyles (e.g., members of the *Mycoplasma* are vertebrate pathogens; Fischer et al. 2012; Waites and Talkington 2004). Even within the mollicute genera that infect fireflies, the preferred host can vary. For example, the *Williamsoniiplasma* genus includes species associated with either lampyrine fireflies or tiger beetles (Tully et al. 1998), which last shared a common ancestor over 300 million years ago (McKenna et al. 2019). Thus, it appears unlikely that mollicute symbionts are the result of ancient domestication in a common host ancestor and conserved through firefly evolution to extant lineages.

This incongruence between host and symbiont phylogenies could have two explanations. First, species from the *Mollicutes* may be generalists and associated with multiple host species. These bacteria could switch hosts on multiple occasions over evolutionary time scales, a process that has been documented in other insect-symbiont systems (Sudakaran et al. 2017). While most *Williamsoniiplasma* species appear to specifically associate with one lampyrine species or genus (Figure 5A), the *Wi. luminosum* species was detected at moderate to high abundance in three *Photinus* firefly species (Figure 5A; Figure S11; Fallon et al. 2018; Williamson et al. 1990), suggesting a lower level of host specificity for this mollicute species. This is also supported by *En. ellychniae*, a bacterial strain with a 16S rRNA amplicon sequence almost identical to *En. corruscae* (Figure 4), being able to infect *Pn. corruscus* as well as tabanid flies (Wedincamp et al. 1996); and by members of the *Spiroplasma* genus, with single *Spiroplasma* species being detected in different insect species (Hackett et al. 1992; 1996; Stevens et al. 1997; Tully et al. 1995). Host switching could conceivably be facilitated by interspecific sexual interactions, a behavior that has been documented among *Photinus* fireflies (Vergara et al. 2025). Future laboratory transplantation experiments could help discern the extent of species-specificity in firefly-mollicute associations.

A second possible explanation for the discrepancy in firefly-mollicute phylogenies is that some of these associations may have evolved recently, leading to patterns of co-phylogeny that may only be detectable at lower taxonomic levels, within single mollicute genera, across species or strains. The *Entomoplasma* and *Williamsoniiplasma* genera, for example, may each include additional firefly-associated symbiont species that have to be discovered from yet unscreened firefly species. Given that *Williamsoniiplasma* species can be endemic to single firefly host species (e.g., *Wi. lucivorax* to *Pn. pyralis*), unscreened lampyrine species could plausibly harbor new endemic mollicute taxa. The microbiome of more firefly species should be characterized to determine whether they harbor new mollicute species, sisters to the ones that have already been identified.

## Conclusion

Through qPCR, deep amplicon sequencing, dissections, phylogenetics, and bioinformatic analyses, we characterized the bacterial communities of fireflies from two common subfamilies in the Northeastern US and evaluated the influence of different factors on bacterial community structure and composition. We found that most firefly species harbor a unique, yet variable microbiome, structured by a combination of stochastic and deterministic forces, and often dominated by members of species-specific mollicutes. Although the intraspecific variation observed in firefly microbiomes likely derives from a combination of stochastic process and geographic, environmental, and host-specific factors, like life stage, age, and sex, the non-random preferential associations with members of the *Mollicutes* raises the possibility that host-specific physiochemical conditions (i.e., habitat filtering mechanisms) contribute to the maintenance and specificity of these bacteria. While the transmission mode of these bacteria is still unknown, given their high abundance when present, mollicutes may play a role in the biology of firefly hosts (Supplementary Text 1). We also occasionally found insect endosymbionts infecting firefly tissues, with yet undetermined roles, but with a hypothetical potential for reproductive manipulation.

Altogether, our results suggest that fireflies are associated with species-specific bacteria that are not always present, and that could, hypothetically, be facultative symbionts. In the future, functional characterizations of firefly microbiomes through combinations of targeted experimental antibiotic and fitness assays, genomics, and gene expression analyses will be necessary to better comprehend the nature of the microbe-firefly associations. Firefly biology and ecology can be challenging to study, given the short adult lifespans of most species and the difficulty to rear early developmental stages, especially terrestrial species, in laboratory settings. Improvements of existing methodologies (Buschman 1988; Fallon et al. 2018; Fu and Benno Meyer-Rochow 2013; Ho et al. 2010; McLean et al. 1972; Vaz et al. 2020; Yang et al. 2024) could facilitate the study of their microbiome. This research constitutes a major first step towards a better understanding of the ecology of firefly-associated microbial communities. Future research will shed light on how microbes contribute to the fascinating but puzzling ecology and evolution of fireflies, a group that provides pest control services (Fu and Benno Meyer-Rochow 2013) and has high aesthetic value, but is experiencing severe population reductions around the world (Fallon et al. 2018; Lewis et al. 2020).

## Materials and Methods

### 1. Specimen collection and identification

A total of 344 firefly specimens (*Coleoptera*: *Lampyridae*) were collected opportunistically in the continental United States in 2021, 2022, and 2023 (Figure 1A and 1B; Table S1). We captured specimens by hand or with a net, preserved them in > 95% ethanol upon collection, and froze them within the next 72 hours. Firefly specimens were initially identified and sexed via examination of flash patterns, external morphology, and genitalia (Faust 2017; Fender 1970; Green 1956; 1957; McDermott 1967). To verify species identifications, up to 27 specimens from each species and species group were DNA barcoded, for a total of 59 specimens. We extracted DNA as described below and amplified fragments of the universal metazoan cytochrome C oxidase subunit 1 (COI) gene marker using four different primer pairs (Folmer et al. 1994; Geller et al. 2013; Rennstam Rubbmark et al. 2018; Simon et al. 1994; Stanger-Hall et al. 2007). A full description of the barcoding methods is available in Supplementary Text 1 with results shown in Figure S1. DNA barcoding sequences are available as NCBI GenBank files, with accession IDs PV844642-PV844700.

The firefly specimens belonged to two subfamilies, four genera, and 12 species or species groups (Figure 1A). We chose to use species group-level classification for three groups (*Photinus consanguineus*, *Pn. marginellus*, and *Photuris versicolor* groups) because (*i*) identification to the species level in the field is challenging in some firefly groups based solely on external morphological characters and flash patterns (e.g., *Photuris*; Faust 2017), (*ii*) while male genitalia morphology can be diagnostic in some firefly groups (e.g., *Photinus*; Green 1956), dissection to extract the aedeagus may compromise the tissues needed for microbiome analyses, and (*iii*) species relationships cannot be resolved with the COI gene for some groups (e.g. *Photuris*, some *Photinus*, *Pyractomena*; Lower et al. 2017; Stanger-Hall and Lloyd 2015), either due to lack of variation in the COI locus, incomplete lineage sorting, or lack of COI sequence in GenBank.

Among the collected firefly specimens, 332 were adults, including 104 females, 227 males, and one *Lucidota atra* specimen that was not sexed, and 12 were juveniles, with nine larvae and three pupae (Figure 1A). Ten adult beetles from other families were also collected as outgroup specimens. They included three soldier beetles (*Cantharidae*), two tiger beetles (*Cicindelidae*) that were already dead upon collection, three click beetles (*Elateridae*), one net-winged beetle (*Lycidae*), and one fire-colored beetle (*Pyrochroidae*).

### 2. Dissection and DNA extraction

To study the bacterial communities of fireflies, we extracted DNA from the whole body of 321 specimens. First, fully intact specimens were surface sterilized in 10% bleach, > 95% ethanol, and sterile water. We then flash-froze specimens in empty 1.5 mL tubes by dipping them in liquid nitrogen, before grinding them to powder with a sterile pestle. For each DNA extraction batch (10–23 samples each), a DNA extraction blank sample, consisting of an empty tube, was processed simultaneously with the other samples. DNA was subsequently extracted from flash-frozen, ground samples using the Qiagen DNeasy Blood and Tissue kit (Qiagen, Germantown, MD, USA), following the manufacturer’s pretreatment protocol for Gram-positive bacteria.

To study the bacterial community of fireflies across tissues, we randomly selected among our preserved specimens 5–6 adult females and males from each of three well-represented firefly taxa (*Photinus corruscus*, *Pn. pyralis*, and the *Pr. versicolor* group; Table S8). We first surface-sterilized the specimens as described above. To isolate tissues, firefly specimens were then dissected in sterile water using bleach-sterilized fine forceps. Separated tissues were placed in pre-chilled 1.5 mL tubes, either filled with 95–100% ethanol or empty, and stored at -80 °C. During the dissections, we separated the entire gut, the reproductive organs, and the lantern (for bioluminescent species) from the rest of the body, which was used as the fourth main tissue (i.e., “carcass”). To investigate the possibility of endosymbionts (i.e., symbionts residing inside the body or cells of another organism) and/or reproductive manipulators, for a few *Pn. corruscus* and *Pn. pyralis* specimens, we additionally separated some of these main tissues to obtain enlarged fat tissue (“globules”), genitalia, ovaries, hindgut, foregut and midgut, spermatheca, spermatophore-digesting glands, and testes. Dissection blank samples consisted of a firefly leg left in a water droplet on the side of the dissection area. To control for contaminants during dissection, we used a new dissection blank for every one to three dissected specimens. In total, we dissected 33 adult fireflies, resulting in 132 firefly tissue samples and 15 dissection blank samples (Figure S2). We extracted DNA from dissected tissue samples as described above for whole bodies, but without the surface sterilization step.

We obtained a total of 497 DNA samples, including 321 whole-body samples, 132 dissected tissue samples, 15 dissection blanks, and 29 DNA extraction blank samples.

### 3. Bacterial titer and total DNA quantifications

Total bacterial titer per sample was obtained through qPCR (Table S10). CT values were generated with a 7300 Real-Time PCR System (Applied Biosystems, Foster City, CA, USA). We used the V4 region of the 16S rRNA gene as the target, amplified using the bacterial universal 515F (5’-GTGCCAGCMGCCGCGGTAA-3’) and 806R (5’-GGACTACHVGGGTWTCTAAT-3’) primers (Apprill et al. 2015; Caporaso et al. 2011; Parada et al. 2016). To obtain absolute bacterial titers measured in 16S rRNA gene copy numbers, we used, in each qPCR plate, a standard curve built on a ten-fold dilution series of a DNA sample with known 16S rRNA gene copy number. Bacterial titers were then corrected using DNA extraction blank and dissection blank samples for tissue samples. Each sample was run in two or three separate technical replicates, consisting of runs on separate qPCR plates. We calculated the average of blank-corrected titers across plate replicates and used these values in downstream statistical analyses. A detailed description of our qPCR procedure can be found in Supplementary Text 1. To normalize bacterial titers and for our Illumina amplicon sequencing decontamination process (see subsection 5.1), we quantified the concentration of extracted DNA per sample using a Qubit 4 fluorometer (Invitrogen, Waltham, MA, USA) and Qubit dsDNA HS and BR kits (Thermo Fisher Scientific, Waltham, MA, USA), with two to three technical replicates per DNA sample.

### 4. Illumina library preparation and sequencing

A first batch of 20 samples was sequenced at the University of Texas at Arlington Genomic Core Facility in 2021. The library was prepared following Illumina’s two-step amplification strategy using primers 341F (5’-CCTACGGGNGGCWGCAG-3’) and 785R (5’-GACTACHVGGGTATCTAATCC-3’), which amplify the V3-V4 hypervariable region of the 16S rRNA gene (Klindworth et al. 2012), as described in (Ravenscraft et al. 2024). Each sample was bidirectionally sequenced on a 600 cycle paired-end run on an Illumina MiSeq platform (Illumina, San Diego, CA, USA). The second batch, containing 477 samples plus one sequencing blank, was sequenced at the Environmental Sample Preparation and Sequencing Facility at Argonne National Laboratory in 2024 (Table S11). The V4 region of the 16S rRNA gene was PCR-amplified with 515F and 806R primers (Caporaso et al. 2012). Amplicons were sequenced on a 300 cycle paired end run in a NextSeq 2000 instrument (Illumina) using customized sequencing primers and procedures. For both runs, demultiplexed data were then processed in R (R Core Team 2024) for downstream analysis.

### 5. Bioinformatic pre-treatment

#### 5.1. Pre-processing of sequencing data

For the MiSeq sequencing run, priming sites and poor-quality bases were removed from the 5’ and 3’ ends of the sequences using the cutadapt v3.2 program (Martin 2011). This step was not necessary for the NextSeq sequences, which did not include priming sites. Then, for each sequencing run, we used the *DADA2* v1.32.0 R package (Callahan et al. 2016) to remove low quality reads, merge paired ends, and infer the bacterial strains present, generating amplicon sequence variants (ASVs). We merged data from the two sequencing runs, performed *de novo* chimera removal, and ran taxonomic classification through the RDP classifier and with a modified SILVA SSU v138.1 training dataset (Quast et al. 2013; Wang et al. 2007). Further details on the pre-processing procedure are available in Supplementary Text 1. Filtered amplicon sequences were exported in fasta format using the *seqinr* v4.2.36 R package (Charif and Lobry 2007) and are available in our data repository (https://doi.org/10.6084/m9.figshare.28306616). Since ASVs from the 20 samples on the MiSeq run were a longer, partially overlapping region of the 16S rRNA and therefore could not be merged with the NextSeq ASVs, we performed all analyses at higher taxonomic levels (e.g., bacterial genera) and grouped key bacterial taxa phylogenetically (see section 5.2 below).

The ASV count table, the taxonomic classification table, and the metadata information (including bacterial titer obtained from qPCR and DNA concentration per sample) were converted into a phyloseq object that contained 99,979 ASVs, using the *phyloseq* v1.48.0 and *speedyseq* v0.5.3.9021 R packages (McLaren 2020; McMurdie and Holmes 2013). Following length filtering (removing 14,757 ASVs that were not between 240 and 430 bp; i.e., 15% of the sequences), removal of ASVs annotated as chloroplasts, mitochondria, or unidentified at the phylum level (removing 6,778 ASVs and 8% of the remaining sequences), and removal of ASVs only obtained from sequencing blank and extraction blank samples (removing 23,724 ASVs and 31% of the remaining sequences), we used the *decontam* v1.24.0 R package (Davis et al. 2018), with the stringent “either” frequency or prevalence method to discard contaminant ASVs. With this method, 1,555 ASVs were flagged as contaminants and removed (discarding 2% of the remaining sequences; list available in our data repository: https://doi.org/10.6084/m9.figshare.28306616), resulting in 53,165 ASVs being retained in our dataset. While we removed a high number of ASVs classified as common contaminants with this method (e.g., *Klebsiella*, *Pseudomonas*, *Salmonella*, *Staphylococcus*), we note that it discarded an abundant ASV from the *Acinetobacter* genus; similar studies have also sequenced *Acinetobacter* from fireflies and it is possible this is a common environmental colonist of firefly guts (Green et al. 2021). In addition, a list of bacteria that passed our decontamination pipeline but that could conceivably be contaminants, because they were only relatively abundant in specimens that had a very low bacterial titer, is shown in Table S12 with more details in Supplementary Text 1.

Finally, to control for among-sample differences in sequencing depth, our data were rarefied to 30,000 sequences per sample using *phyloseq* (Schloss 2023; 2024; Weiss et al. 2017). Visual inspection of rarefaction curves indicated that observed diversity had plateaued in the samples by this depth. Because rarefying would remove important samples in our dissected tissue dataset, we instead used compositional normalization for tissue comparative analyses, computed with the *microViz* v0.12.4 R package (Barnett et al. 2021). Similarly, rarefied data were not used for our multivariate ordination analyses based on Bray-Curtis distances, since compositional normalization has been shown to be a more robust normalization method for clustering on Bray-Curtis distance matrices (McMurdie and Holmes 2014) and because rarefying caused some species to be removed from our dataset (data from our two *Pyractomena* sp. near *sinuata* whole body specimens did not pass the rarefaction threshold).

#### 5.2. Taxonomic annotation refinement through phylogenetic analyses

We next performed phylogenetic analyses to refine the taxonomic identification of abundant and prevalent bacterial clades in fireflies (*Mollicutes*; Figure S9), common arthropod endosymbionts (*Rickettsiaceae*, *Rickettsiella*, and *Wolbachia*; Figure S10), and insect pathogens (*Serratia*; Figure S10). For the *Mollicutes*, we used the terminologies from (Gupta et al. 2019), although other authors classify most of the firefly-associated non-*Spiroplasma* mollicutes as the *Entomoplasma* genus (Balish et al. 2019; Gasparich and Kuo 2019; Lo et al. 2018; Yan et al. 2024; but see Arahal et al. 2022; Gupta et al. 2018; Gupta and Oren 2020). Because our data came from two sequencing batches that used two different amplicon lengths, hence classifying identical variants as two different ASVs, we used phylogenetic analyses to group together ASVs that were likely from the same microorganism but were classified as different variants due to different amplicon sizes. We use the term “bacterial phylotype” hereafter in this section, approximately equivalent to bacterial species, to refer to this classification, which we performed only for the five aforementioned bacterial clades (*Mollicutes*, *Rickettsiaceae*, *Rickettsiella*, *Serratia*, and *Wolbachia*).

To build these phylogenies, we first selected the 16S rRNA gene amplicon sequence of the most abundant ASVs (those numbered from sv0001 to sv1500, sorted by abundance in our dataset) that were classified as either from the *Mollicutes* class (i.e., *Entomoplasmatales* and *Mycoplasmatales* orders *sensu* SILVA database), from the non-*Anaplasmataceae Rickettsiales* order (i.e., including the *Rickettsiaceae* family, unidentified *Rickettsiales*, and *Candidatus* Hepatincola), or from one of the focal bacterial genera (*Rickettsiella*, *Serratia*, and *Wolbachia*). We used these sequences in NCBI BLASTn runs and downloaded the 16S rRNA gene sequence of the closest matches. We also downloaded the sequences of reference, representative bacterial species from NCBI. For the *Mollicutes*, representative amplicon sequences of each of the firefly-associated bacterial species found in Green et al. (2021) were also used, as well as a few insect-associated mollicute sequences from Funaro et al. (2011). With all these sequences, we produced Clustal Omega (Sievers et al. 2011) multiple sequence alignments in SeaView v5.0.5 (Gouy et al. 2009), which were used to build maximum likelihood phylogenies using RAxML Blackbox (Stamatakis 2014), implemented in the CIPRES Science Gateway (Miller et al. 2010), with 999 bootstrap iterations. All phylogenetic trees were visualized and manually annotated in iTOL (Letunic and Bork 2016). Further details on these phylogenetic methods can be found in Supplementary Text 1.

Finally, we manually edited the *phyloseq* taxonomy table to account for this taxonomic refinement (available as a Data Repository: https://doi.org/10.6084/m9.figshare.28306616). Except for analyses on alpha diversity metrics and specific mentions of analyses ran at the bacterial order or genus levels, all downstream analyses were run on microbiome data aggregated at the genus level, except for the aforementioned groups we refined (*Mollicutes*, *Rickettsiaceae*, *Rickettsiella*, *Serratia*, and *Wolbachia*) which were aggregated at the “phylotype” level (Figure S2). This combination of genus and phylotype level classification is hereafter referred to as “taxon” level. Data was aggregated at the bacterial order, genus, and taxon levels using the *microViz* R package.

### 6. Bioinformatic analyses

#### 6.1. Stochasticity analyses

To infer the relative importance of stochastic *vs.* deterministic processes contributing to microbiome assembly, we calculated the normalized stochasticity ratio (i.e., NST index) for each firefly species that had more than two specimens in our dataset through implementation of null-models using the *NST* R package v3.1.9 (Ning et al. 2019). An index value between 0.5 and 1 suggests that stochastic processes shape community assembly, whereas a value between 0 and 0.5 indicates higher influence of deterministic processes. Next, to infer the contribution of specific bacterial groups, we tested how the removal of either all the mollicutes, all putative endosymbionts (*Ca.* Hepatincola, *Rickettsia*, *Rickettsiella*, and *Wolbachia*), or both, from the microbiome data of each firefly species affects the NST index value. We generated 95% confidence intervals via bootstrapping and tested NST variation between the unaltered microbiome of firefly species *vs.* that following removal of the aforementioned bacterial groups through pairwise, two-tailed Wilcoxon signed-rank statistical tests (Table S2).

#### 6.2. Alpha and beta diversity

To assess patterns of diversity, richness, evenness, dominance, and rarity in bacterial communities from fireflies, indices of alpha diversity were calculated on rarefied data with non-aggregated (ASV level) taxonomic tables, using the *microbiome* v1.26.0 R package (Lahti and Shetty 2017) (Table S4). To compare bacterial composition across firefly species, genera, subfamilies, and diverse factors, we used beta diversity metrics, first running ordinations on Bray-Curtis distance matrices computed from non-rarefied, proportional abundance data using nonmetric multidimensional scaling (NMDS) analyses computed with *phyloseq*, to infer similarities among fireflies from different taxa, diets, adult use of bioluminescence, regions, and tissues. To test how different factors were correlated with bacterial composition across fireflies, we then ran permutational multivariate analysis of variance (PERMANOVAs) on rarefied, taxon-aggregated data, with the *vegan* v2.6.4 R package (Oksanen et al. 2016), and pairwise PERMANOVAs as post hoc tests using the *pairwiseAdonis* v0.4.1 R package (Martinez 2020).

#### 6.3. Host relatedness, geographic distance, and microbiome dissimilarity

The impact of firefly host phylogeny on bacterial community dissimilarity was assessed using Mantel tests. We first generated a host phylogeny using five previously sequenced firefly genes (Catalan et al. 2022; Martin et al. 2017; Sander and Hall 2015; Stanger-Hall and Lloyd 2015; Stanger-Hall et al. 2007) (Figure S1; Supplementary Text 1). We then used the *ape* v5.8 R package (Paradis et al. 2004) to generate a phylogenetic distance matrix from our tree with the cophenetic.phylo() function. We next produced a matrix of geographic distance among collected firefly specimens calculated from latitude and longitude decimal variables at collection using the *sf* v1.0.17 R package (Pebesma 2018). As proxies for microbiome dissimilarity, we generated two pairwise distance matrices using the Bray-Curtis distance from rarefied relative bacterial abundance between firefly specimens and the Bray-Curtis binary distance from bacterial presence/absence (equivalent of Sørensen-Dice distance) between firefly specimens. Bacterial presence was defined as any bacterial taxon with at least 1% relative abundance in a specimen and absence as less than 1% relative abundance in that specimen. Because the number of pairwise comparisons was not computationally tractable, we applied a filter to our dataset, keeping only bacterial taxa that had more than 0.5% relative abundance in at least 5% of all firefly specimens. These microbiome dissimilarity matrices were compared to a matrix of phylogenetic distance and to geographic distances between firefly specimens through partial Mantel tests using the *vegan* R package, allowing us to control for host phylogeny and geography, respectively.

#### 6.4. Contribution of firefly ecological traits to bacterial abundance and diversity

We tested the influence of adult bioluminescence, diet, and body size of firefly species on bacterial abundance and diversity, as well as correlations between bacterial abundance and diversity metrics via generalized least square analyses with the *nlme* v3.1.164 R package (Pinheiro et al. 2024). To account for the effects of phylogenetic nonindependence in comparisons of firefly species-level covariates, we used phylogenetic generalized least square (PGLS) analyses, fitting a Brownian motion model of evolution. The firefly phylogeny was obtained as described in section 6.3 and Supplementary Text 1 (Figure S1). Bacterial titer and diversity indices were averaged per firefly species. We also averaged body size (i.e., total adult male body length) per firefly species or species groups using the ranges from Faust (2017), Fender (1970), Green (1956, 1957), and Lloyd (1969).

#### 6.5. Bacterial composition and differential abundance analysis

To test whether single bacteria vary in their abundance across fireflies and factors, we used the *MaAsLin2* v1.15.1 R package (Mallick et al. 2021), fitting our rarefied (or non-rarefied for tissue analyses) data to negative binomial distributions and controlling for random effects (see section 6.7. Statistical tests) in multiple generalized linear mixed models. To account for variables with unbalanced discrete levels (e.g., species with high *vs.* low number of specimens sampled), when more than two discrete levels were present in our fixed effect(s), we ran the same *MaAsLin2* models multiple times, changing the reference level each time (Table S5, S6, S7, and S9). We filtered out bacterial taxa detected in less than 1% of all firefly samples being tested, unless the test was run on groups with very low sample size (prevalence threshold lowered to 0.1% instead).

#### 6.6. Bacterial co-occurrence analysis

To investigate positive and negative co-occurrence patterns among bacteria in adult fireflies, we ran co-occurrence analyses, within firefly species that had more than eight adult individuals in our dataset (i.e., excluding *Lu. atra*, *Pn. consanguineus* group, *Pn. cookii*, and *Py. borealis*) and grouping all *Photuris* together. We used both the *cooccur* v1.3 and *CooccurrenceAffinity* v1.0 R packages (Griffith et al. 2016; Mainali and Slud 2022). A filter was applied before analyses to obtain a manageable number of bacterial taxa to be tested per firefly species. Bacterial taxa were considered present when they had at least 1% relative abundance (absent otherwise) in a specimen and were kept in these analyses when they had more than 10,000 reads—or 20,000 reads for the *Photuris—*in one or more samples for a given firefly species or species group.

#### 6.7. Statistical tests

In addition to the tests described in the previous sections, differences in alpha diversity and qPCR titers across hosts and diverse environmental factors were assessed using linear mixed models (LMM) with the *lme4* v1.1.35.5 R package (Bates et al. 2015), which allowed us to control for random effects that may otherwise bias our interpretation of statistical tests. Some key factors were always kept as random effects in our models because they had a strong influence on the microbiome of fireflies in our datasets. These random factors included the year of collection, the geographic location of sampling (either address, longitude, latitude, or region), and collection day or month of year (a rough proxy for firefly age in many species). When they were not among the fixed effects in our models, host sex (which included developmental stages) and species were used as random effects or covariates. The importance of other random effects in our models was previously tested through bidirectional model selection with likelihood ratio tests and comparing model’s AICs.

We tested the distribution of the residuals, percentage of the dispersion of the data, the presence of outliers, and zero-inflation using the *performance* v0.12.3 and *DHARMa* v0.4.6 R packages (Hartig 2018; Lüdecke et al. 2021). Our models were often flagged as poor fits due to zero-inflation, despite normalization and log10 transformation. When this was the case, we fitted our data to a negative binomial distribution in generalized LMMs (GLMMs) using *lme4*. We next performed pairwise post hoc tests using the *emmeans* v1.10.4 R package (Lenth 2022) to compute the estimated marginal means (Searle et al. 1980). In doing so, we applied an asymptotic degrees-of-freedom method, thus invoking *z*-statistics, which are computationally faster and recommended with larger sample sizes. A description of all LMMs, GLMMs, PERMANOVAs, pairwise PERMANOVAs, Mantel tests, OLS, and PGLS models is available in Table S3.

Additional files, including sequencing OTU and taxonomy tables, list of contaminant variants removed, multiple sequence alignments, photographs of insect specimens, multi-locus backbone phylogenetic trees, R script and input files, and additional supplementary figures, are available in our data repository at https://doi.org/10.6084/m9.figshare.28306616.

## Acknowledgements

We thank Meghan Catherwood, Susan Deering, Natalie Dyer, Hannah Holmes, Katie O’Connor, Nathan Peot, Sam Pring, Lauren Shaffer, Edith Simpson, Asher Timar, Jasmine Timar, and Morgan Ulrich for assistance with specimen collection. We are grateful to Moria Chambers who provided logistical support, laboratory space, and materials. Thanks also go to Johnathan Adamson and Kim Bowles from the University of Texas at Arlington and Stephanie Greenwald the Argonne National Laboratory who performed Illumina library preparation and sequencing.

## Author Contributions

BB designed the study, performed the dissections, analyzed the data, and wrote the initial manuscript. BB and SEL collected specimens. BB, SRN, and AR extracted DNA. BB and TJD performed DNA barcoding. BB and AR processed the amplicon sequencing data. BB, SEL, and AR contributed to review and editing of the manuscript. All authors read and approved the final version of the manuscript.

## Dada Availability

All 16S rRNA gene amplicon sequencing data have been deposited in the NCBI Short Read Archive under BioProject PRJNA1237523. The COI DNA barcoding sequences produced from Sanger sequencing are available from the NCBI GenBank database with accession numbers PV844642–PV844700. Supplementary data can be found in our data repository at: https://doi.org/10.6084/m9.figshare.28306616.

