## Supplementary material for "Species-specific bacterial associations emerge from stochastically assembled microbiomes in Northeastern American fireflies": Figure S

### Supplementary Figures S1–S24

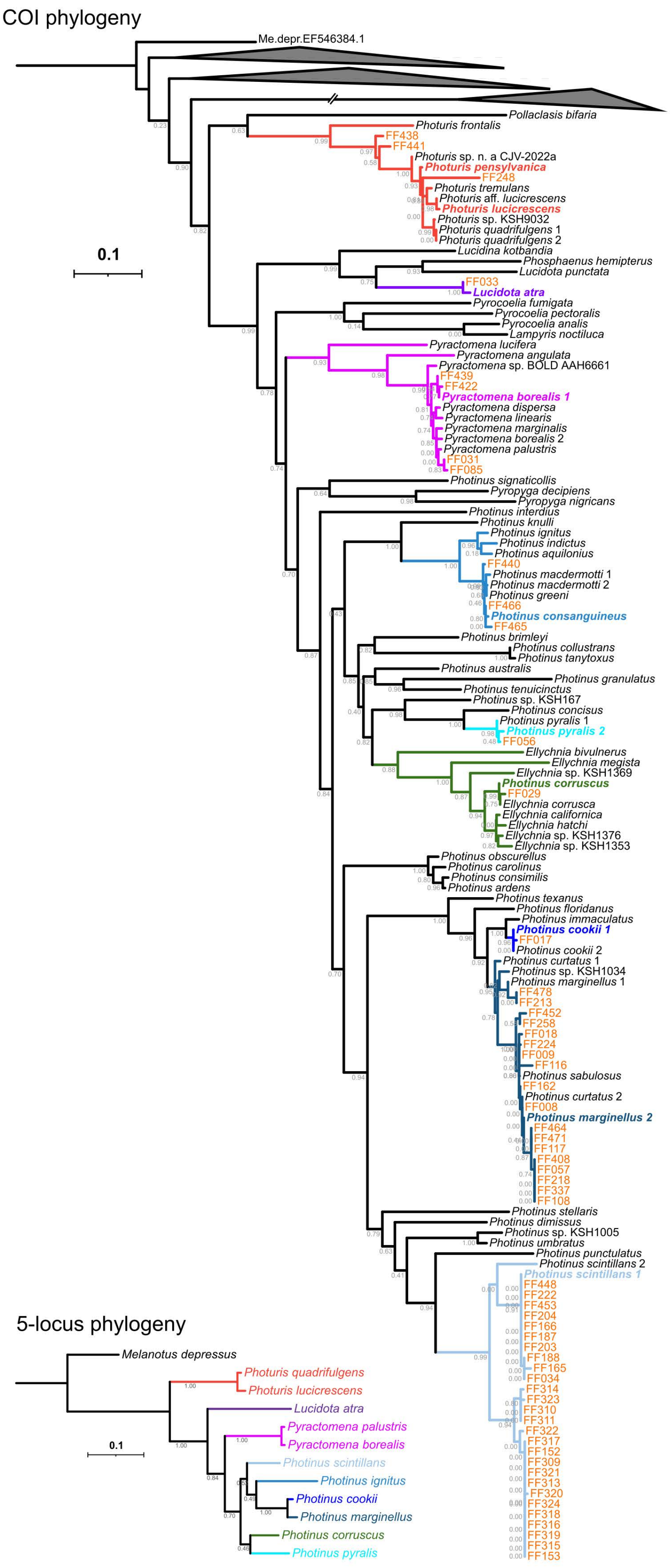
COI phylogeny
Me.depr.EF546384.1
Pollaclasis bifaria
Photuris frontalis
FF438
FF441
Photuris sp. n. a CJV-2022a
Photuris pensylvanica
FF248
Photuris tremulans
Photuris aff. lucicrescens
Photuris lucicrescens
Photuris sp. KSH9032
Photuris quadrifulgens 1
Photuris quadrifulgens 2
0.1
Lucidina kotbandia
Phosphaenus hemipterus
Lucidota punctata
FF033
Lucidota atra
Pyrocoelia fumigata
Pyrocoelia pectoralis
Pyrocoelia analis
Lampyris noctiluca
Pyractomena lucifera
Pyractomena angulata
Pyractomena sp. BOLD AAH6661
FF439
FF422
Pyractomena borealis 1
Pyractomena dispersa
Pyractomena linearis
Pyractomena marginalis
Pyractomena borealis 2
Pyractomena palustris
FF031
FF085
Photinus signaticollis
Pyropyga decipiens
Pyropyga nigricans
Photinus interdius
Photinus knulli
Photinus ignitus
Photinus indictus
Photinus aquilonius
FF440
Photinus macdermotti 1
Photinus macdermotti 2
Photinus greeni
FF466
Photinus consanguineus
FF465
Photinus brimleyi
Photinus collustrans
Photinus tanytoxus
Photinus australis
Photinus granulatus
Photinus tenuicinctus
Photinus sp. KSH167
Photinus concisus
Photinus pyralis 1
Photinus pyralis 2
FF056
Ellychnia bivulnerus
Ellychnia megista
Ellychnia sp. KSH1369
Photinus corruscus
FF029
Ellychnia corrusca
Ellychnia californica
Ellychnia hatchi
Ellychnia sp. KSH1376
Ellychnia sp. KSH1353
Photinus obscurellus
Photinus carolinus
Photinus consimilis
Photinus ardens
Photinus texanus
Photinus floridanus
Photinus immaculatus
Photinus cookii 1
FF017
Photinus cookii 2
Photinus curtatus 1
Photinus sp. KSH1034
Photinus marginellus 1
FF478
FF213
FF452
FF258
FF018
FF224
FF009
FF116
Photinus sabulosus
FF162
Photinus curtatus 2
FF008
Photinus marginellus 2
FF464
FF471
FF117
FF408
FF057
FF218
FF337
FF108
Photinus stellaris
Photinus dimissus
Photinus sp. KSH1005
Photinus umbratus
Photinus punctulatus
Photinus scintillans 2
Photinus scintillans 1
FF448
FF222
FF453
FF204
FF166
FF187
FF203
FF188
FF165
FF034
FF314
FF323
FF310
FF311
FF322
FF317
FF152
FF309
FF321
FF313
FF320
FF324
FF318
FF316
FF319
FF315
FF153
5-locus phylogeny
Melanotus depressus
Photuris quadrifulgens
Photuris lucicrescens
Lucidota atra
Pyractomena palustris
Pyractomena borealis
Photinus scintillans
Photinus ignitus
Photinus cookii
Photinus marginellus
Photinus corruscus
Photinus pyralis
0.1

**Figure S1: Host phylogenies**. The first tree depicts a maximum likelihood phylogeny of the cytochrome c oxidase subunit 1 (COI) gene among firefly species. The tree was rooted with a click beetle *Melatonus depressus* COI sequence (EF546384.1). The tree contains, in addition to reference firefly COI sequences downloaded from NCBI, COI barcoding sequences from our study, shown here with labels in orange font and identifier starting with "FF". Otherwise, branch and label colors correspond to the firefly clade color coding used throughout the study. The first three collapsed clades shown as gray triangles correspond to, from top to bottom, the *Luciolinae* subfamily, the *Lamprohizinae* subfamily, and the *Cheguevaria* genus. The second tree at the bottom focuses on the main firefly species and species groups represented in our study. It was built from a maximum likelihood 5-locus phylogenetic analysis and contains more robust distances among firefly taxa. This tree was used to calculate phylogenetic distances among hosts (see Figure S5). The tree scales represent the number of nucleotide substitutions per site. All bootstrap values are shown on the tree nodes.

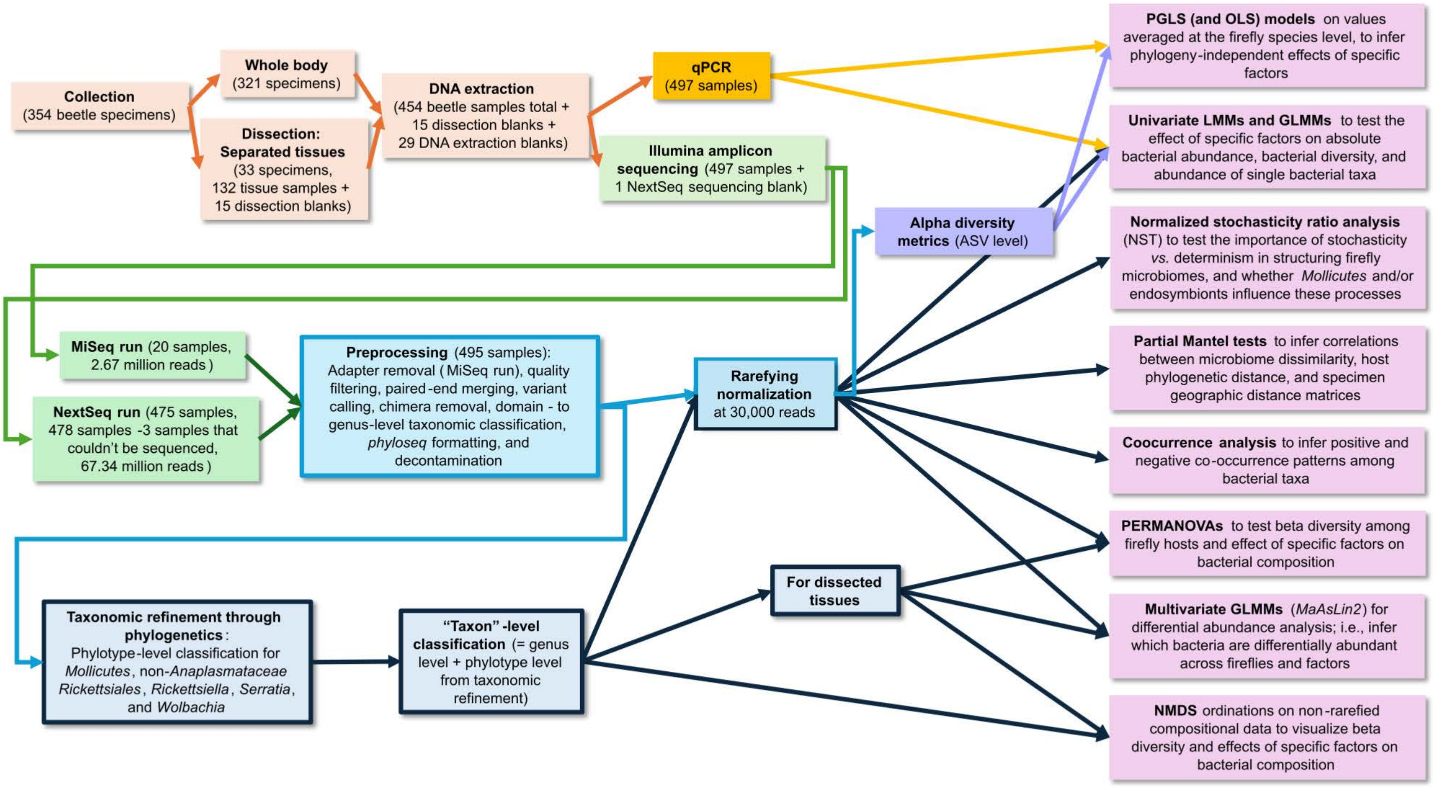


**Figure S2: Overview of the approaches used to study firefly microbiomes**. Following DNA extraction, samples were used in quantitative PCRs and Qubit DNA quantification to obtain bacterial titers and in Illumina amplicon sequencing of the 16S rRNA gene. The bioinformatic approach to process amplicon sequencing reads is given in rectangles with blue colors. Dark blue rectangles correspond to data processed through targeted taxonomic refinement (bacterial "taxon" level). The main statistical analyses performed are shown on the right in pink rectangles.

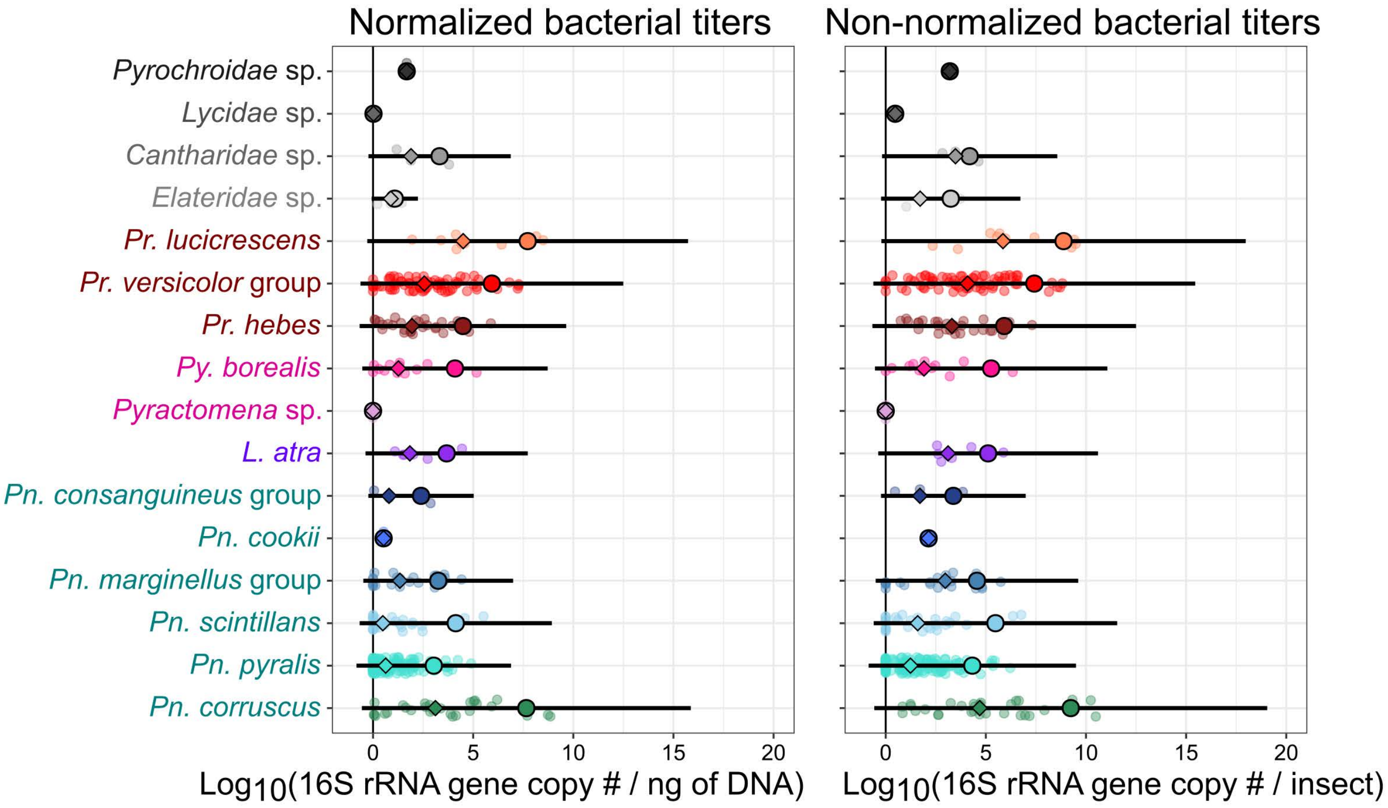


**Figure S3: Normalized and non-normalized absolute bacterial abundance across all firefly and related insect samples**. Values are expressed as the $\log_{10}$ of the number of 16S rRNA gene copies in a sample, measured through qPCR, minus the number of 16S rRNA gene copy in the DNA extraction blank sample from the same batch, and then, only for normalized, divided by the total DNA concentration in the sample, which was measured via Qubit. Balls, diamonds, and error bars represent the average, median and standard deviation, respectively. Raw qPCR data can be found in Table S10.

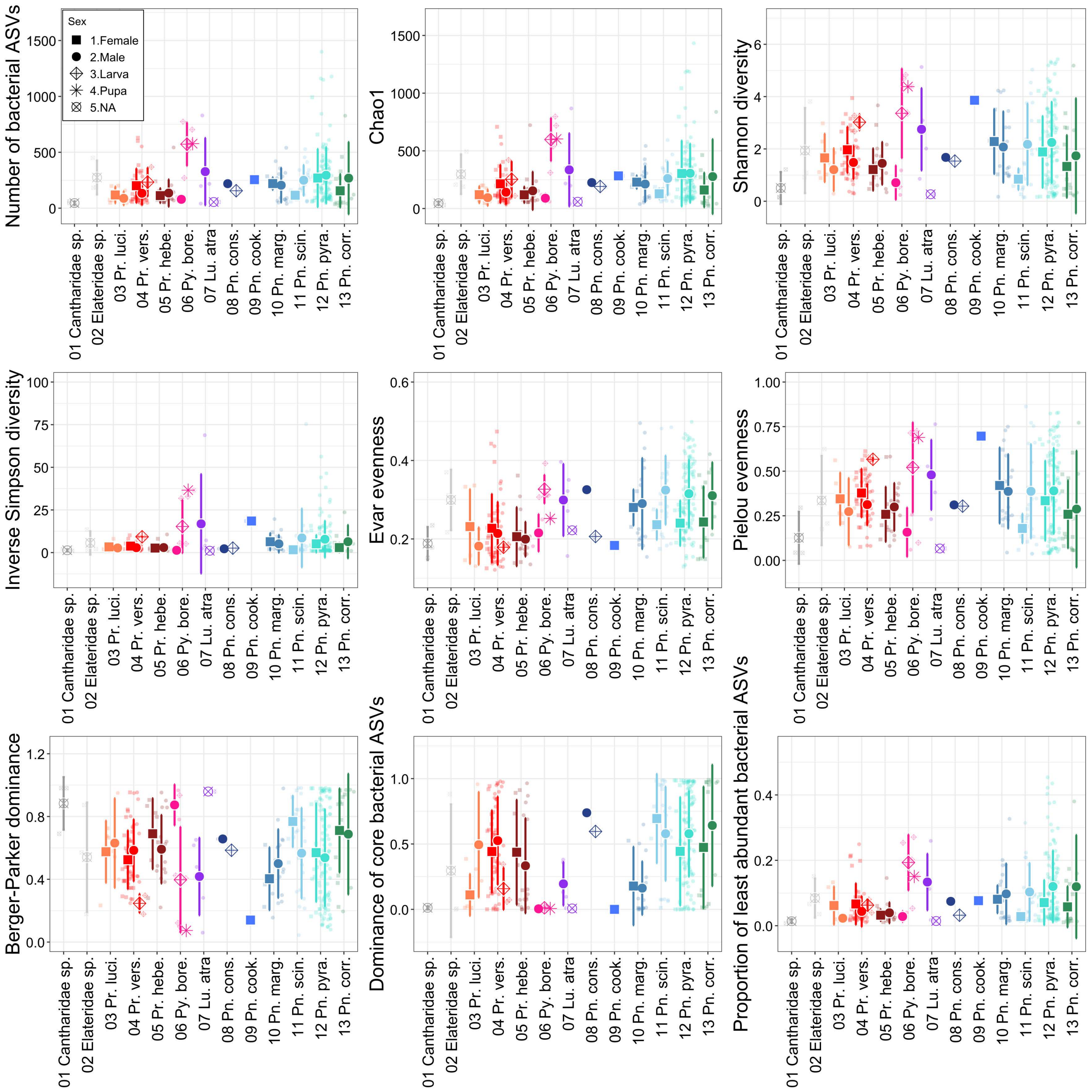


**Figure S4: Alpha diversity indices calculated across firefly species, stages, and sexes**. The indices include the ASV richness, the Chao1index, the Shannon diversity index, the inverse Simpson diversity index, the Smith and Wilson's Evar evenness index, the Pielou evenness index, the Berger-Parker dominance index, the dominance of core bacterial ASVs (those with detection level of 0.1% in over 20% of the samples), and the proportion of the least abundant bacterial ASVs (those below detection level of 0.2% determined separately in each sample) for each firefly species and across sexes and stages. Large size shapes and error bars represent the average and standard deviation, respectively, whereas the dots are the individual index values for each specimen. All alpha diversity metrics can be found in Table S4.

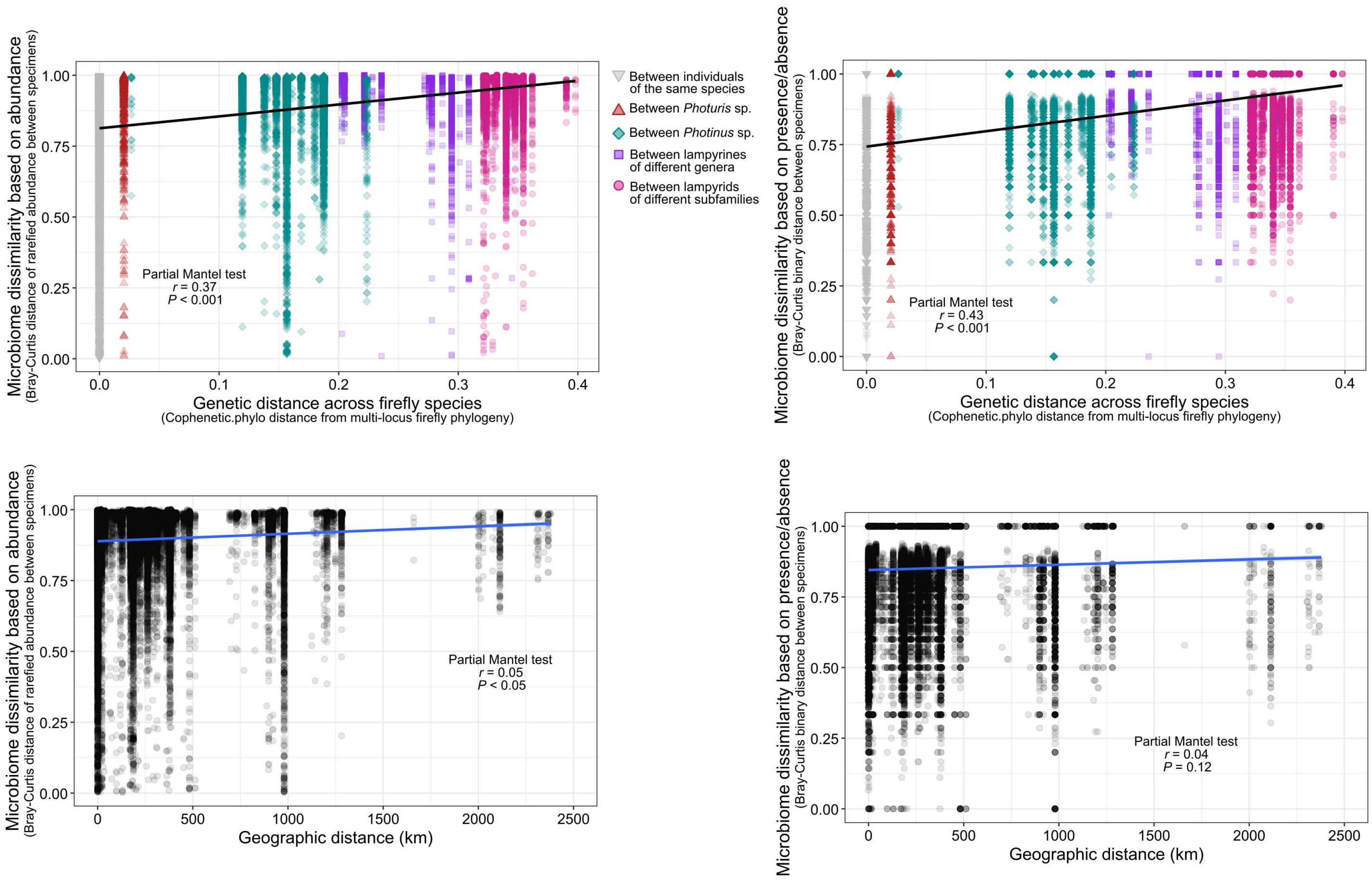


**Figure S5: Microbiome dissimilarity increases with genetic and geographic distances**. The graphs at the top show the positive correlation between pairwise phylogenetic distance across firefly species on the x-axis and pairwise microbiome dissimilarity across firefly specimens on the y-axis, calculated from relative rarefied abundance (graph on the left) and presence absence (graph on the right). Colors and shapes of data points represent the level of species comparisons: between individuals of the same species in down-pointing grey triangles, between species from the *Photuris* genus in up-pointing red triangles, between species from the *Photinus* genus in sea green diamonds, between species from the *Lampyrinae* subfamily in purple squares, and between species from different subfamilies in pink circles. The graphs at the bottom show the weak positive correlation between pairwise geographic distance across firefly collection sites on the x-axis (expressed in kilometers) and pairwise microbiome dissimilarity across firefly specimens on the y-axis, calculated from relative rarefied abundance (graph on the left) and presence absence (graph on the right).

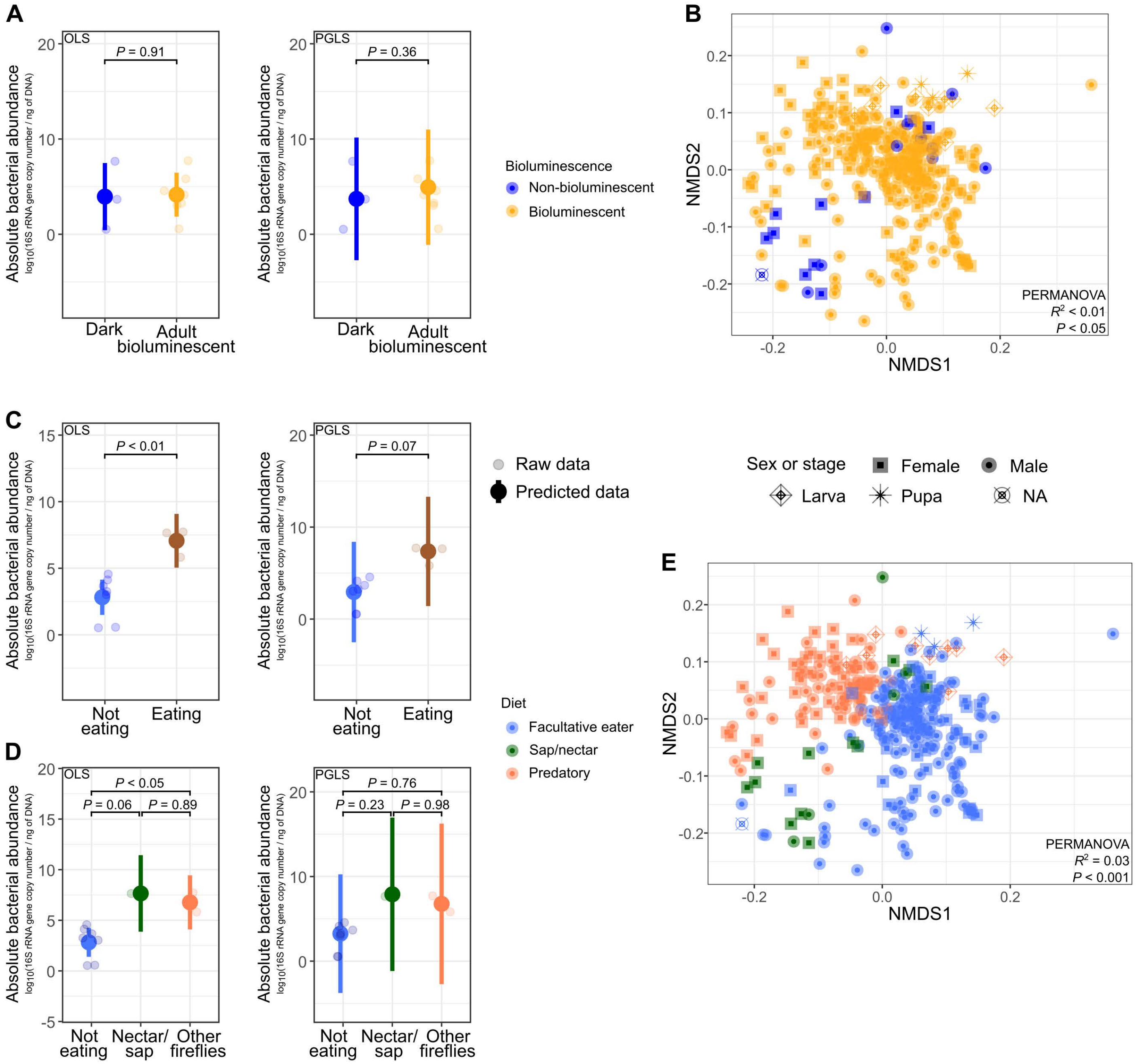


**Figure S6: Adult diet, not bioluminescence, influences the microbiome of fireflies**. **A.** Absolute bacterial abundance is not different between bioluminescent and non-bioluminescent species. The Y-axis shows absolute bacterial abundance (expressed as in Figure 1C) and against firefly species grouped as bioluminescent at the adult stage or not on the x-axis. All values were averaged per firefly species. Opaque balls and error bars correspond to predicted averages per group and 95% confidence intervals, respectively, obtained from ordinary least square (OLS) on the left graph and phylogenetic generalized least square (PGLS), which corrects for phylogenetic similarity biases due to the degree of genetic relatedness between firefly species, on the right graph. **B.** Non-metric multidimensional scaling (NMDS) analysis showing separation of the bacterial communities based on composition and relative abundance between bioluminescent and non-bioluminescent fireflies. Four outgroup specimens were removed from these analyses. The NMDS is derived from Bray-Curtis distances calculated on non-rarefied, compositional relative abundance of taxon-aggregated bacteria in firefly specimens. Permutational multivariate analyses of variance (PERMANOVA) were used for statistical testing of group similarities, computed on read depths rarefied at 30,000 reads. **C**. Absolute bacterial abundance (expressed as in Figure 1C) *vs.* firefly species grouped as eaters (*Photinus corruscus*, *Photuris hebes*, *Pr. lucicrescens*, and *Pr. versicolor* group) or non-eaters (all other species). **D**. Absolute bacterial abundance and bacterial richness *vs.* firefly species grouped based on their adult diet: nectar (*Pn. corruscus*), eating other fireflies (*Pr. hebes*, *Pr. lucicrescens*, and *Pr. versicolor* group), or not eating (all other species). **E**. NMDS analysis showing separation of the bacterial communities based on composition and relative abundance across firefly diets.

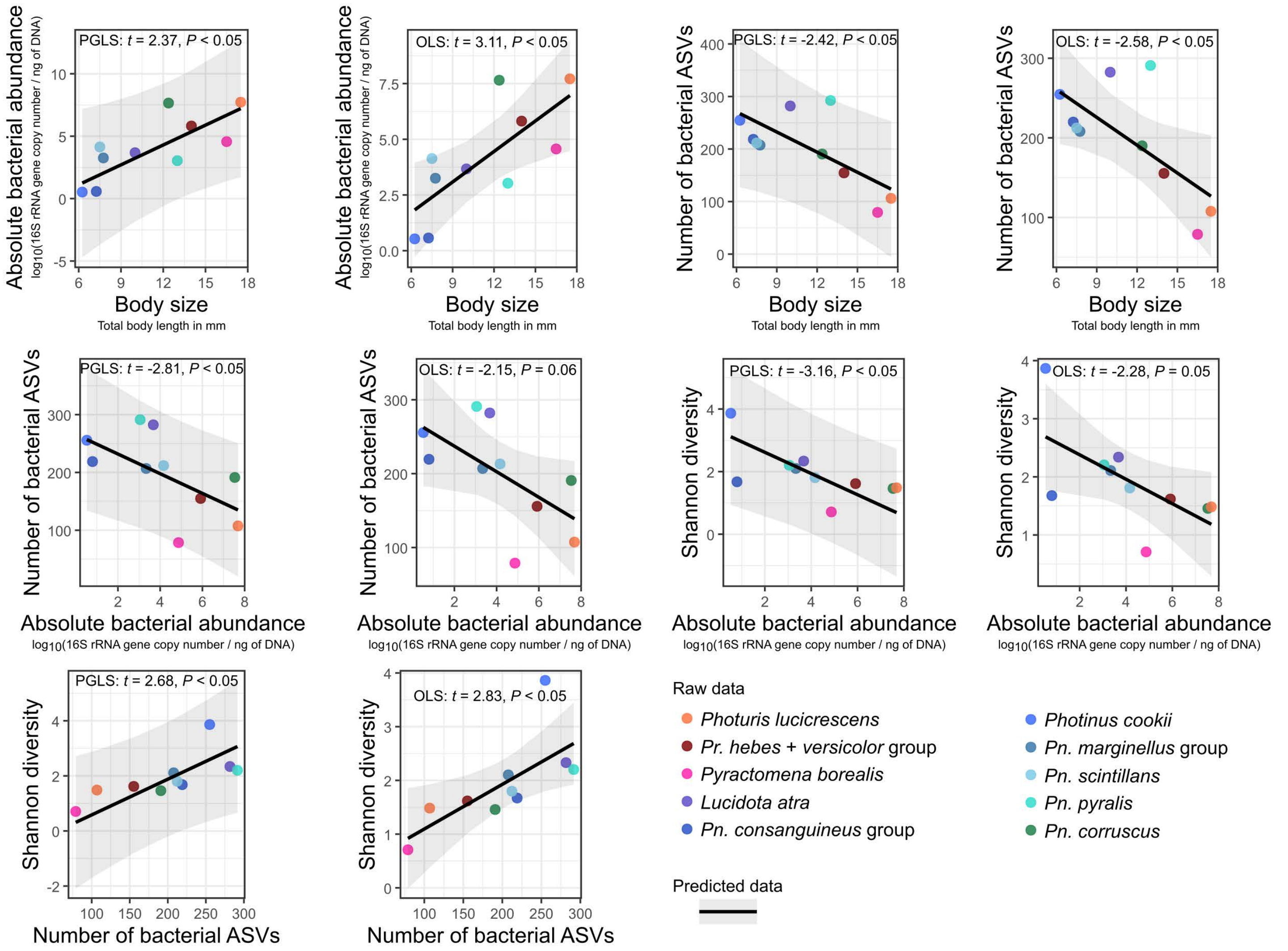


**Figure S7: Relationships between firefly body size, absolute bacterial abundance, richness, and diversity**. The first four graphs (from left to right) show the correlation of absolute bacterial abundance (expressed as in Figure 1C) and bacterial richness *vs.* total firefly body length. The other graphs show how absolute bacterial abundance, bacterial richness, and bacterial diversity correlate with one another in fireflies. All values were averaged per firefly species. Predicted correlation lines and 95% confidence interval shades were obtained from ordinary least square (OLS) and phylogenetic generalized least square (PGLS) models, which correct for phylogenetic similarity biases due to the degree of genetic relatedness between firefly species. Dot colors correspond to firefly species.

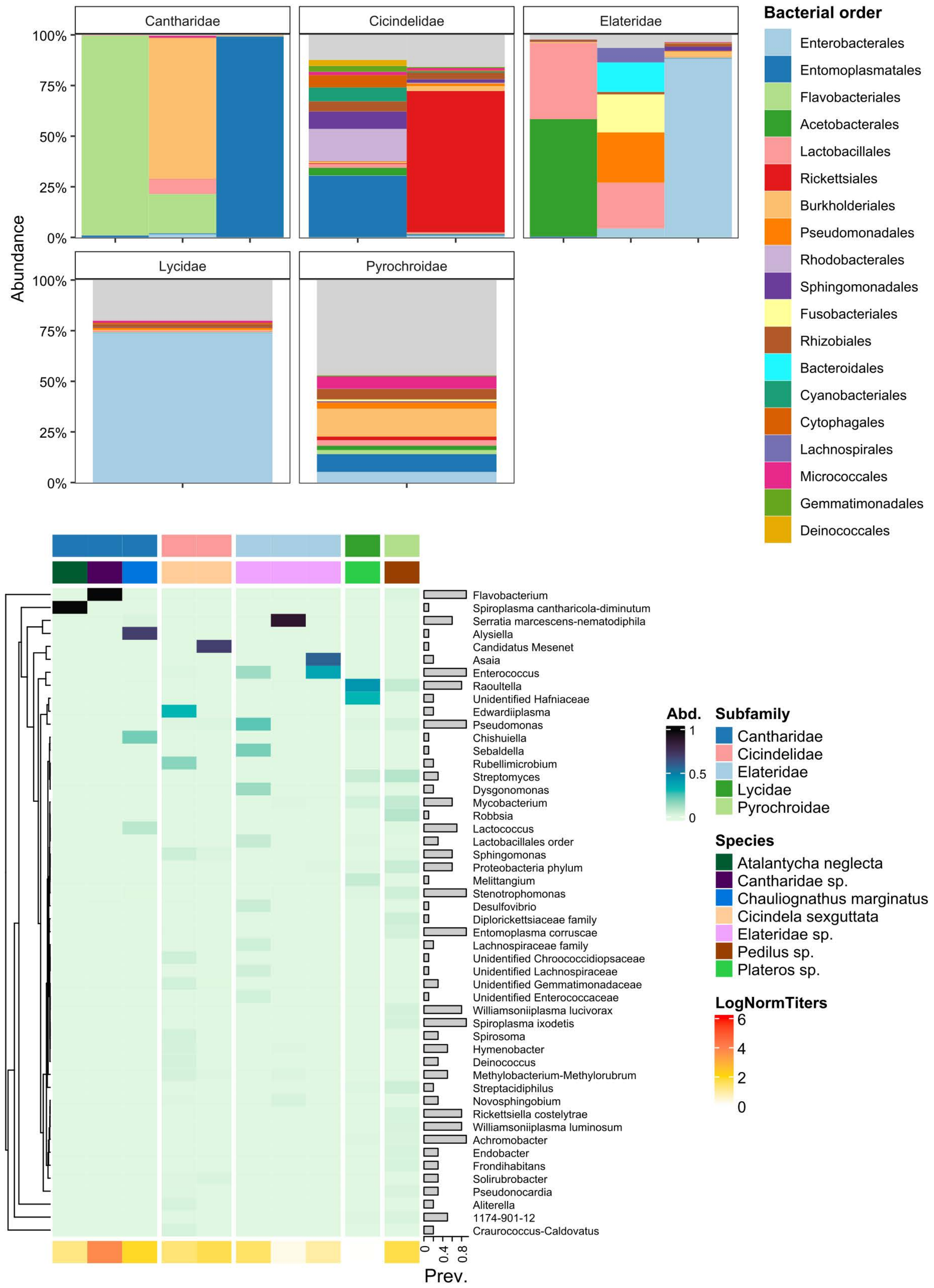


**Figure S8: Bacterial composition in non-lampyrid beetles**. The bar graphs at the top represent non-rarefied bacterial composition at the order level in five beetle groups. Abundance is the proportion of bacteria per specimen, obtained from compositional transformation of read depth. The composite heatmap at the bottom shows bacterial composition and relative abundance at the “taxon” level, along with total absolute bacterial abundance per specimen (expressed as normalized bacterial titer, see Figure 1C) and prevalence of single bacteria across these beetles. Note that the two tiger beetles (*Cicindelidae*) were already dead at the time of collection.

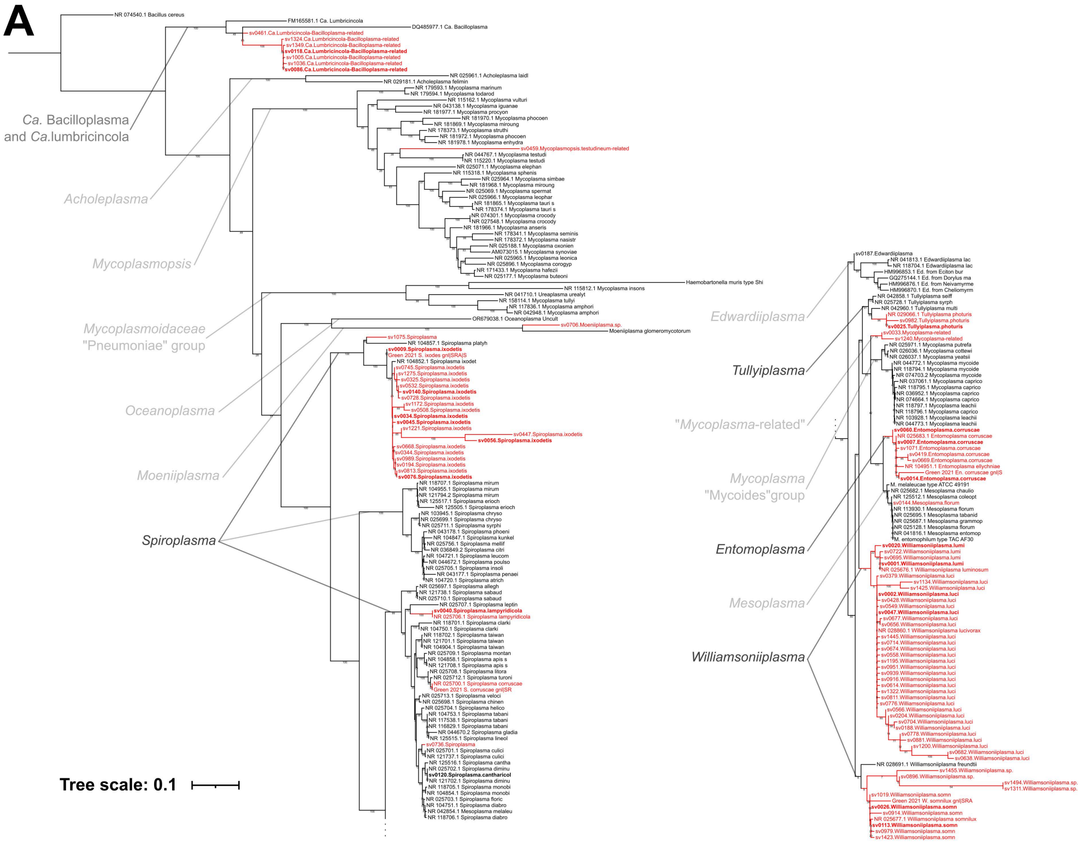


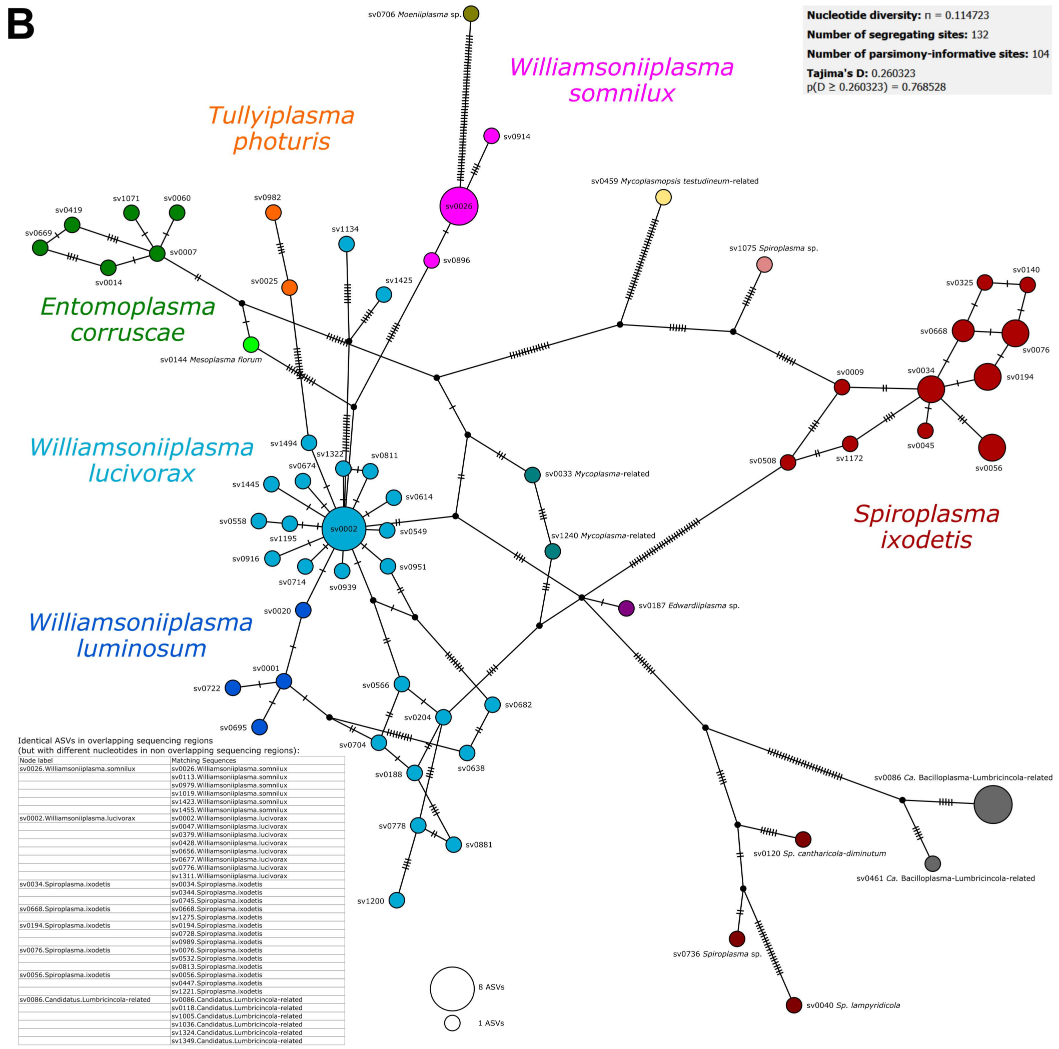


Identical ASVs in overlapping sequencing regions
(but with different nucleotides in non overlapping sequencing regions):

| Node label | Matching Sequences |
|---|---|
| sv0026.Williamsoniiplasma.somnilux | sv0026.Williamsoniiplasma.somnilux |
| | sv0113.Williamsoniiplasma.somnilux |
| | sv0979.Williamsoniiplasma.somnilux |
| | sv1019.Williamsoniiplasma.somnilux |
| | sv1423.Williamsoniiplasma.somnilux |
| | sv1455.Williamsoniiplasma.somnilux |
| sv0002.Williamsoniiplasma.lucivorax | sv0002.Williamsoniiplasma.lucivorax |
| | sv0047.Williamsoniiplasma.lucivorax |
| | sv0379.Williamsoniiplasma.lucivorax |
| | sv0428.Williamsoniiplasma.lucivorax |
| | sv0656.Williamsoniiplasma.lucivorax |
| | sv0677.Williamsoniiplasma.lucivorax |
| | sv0776.Williamsoniiplasma.lucivorax |
| | sv1311.Williamsoniiplasma.lucivorax |
| sv0034.Spiroplasma.ixodetis | sv0034.Spiroplasma.ixodetis |
| | sv0344.Spiroplasma.ixodetis |
| | sv0745.Spiroplasma.ixodetis |
| sv0668.Spiroplasma.ixodetis | sv0668.Spiroplasma.ixodetis |
| | sv1275.Spiroplasma.ixodetis |
| sv0194.Spiroplasma.ixodetis | sv0194.Spiroplasma.ixodetis |
| | sv0728.Spiroplasma.ixodetis |
| | sv0989.Spiroplasma.ixodetis |
| sv0076.Spiroplasma.ixodetis | sv0076.Spiroplasma.ixodetis |
| | sv0532.Spiroplasma.ixodetis |
| | sv0813.Spiroplasma.ixodetis |
| sv0056.Spiroplasma.ixodetis | sv0056.Spiroplasma.ixodetis |
| | sv0447.Spiroplasma.ixodetis |
| | sv1221.Spiroplasma.ixodetis |
| sv0086.Candidatus.Lumbricincola-related | sv0086.Candidatus.Lumbricincola-related |
| | sv0118.Candidatus.Lumbricincola-related |
| | sv1005.Candidatus.Lumbricincola-related |
| | sv1036.Candidatus.Lumbricincola-related |
| | sv1324.Candidatus.Lumbricincola-related |
| | sv1349.Candidatus.Lumbricincola-related |

**Figure S9: Classification of strains from the *Mollicutes***. **A**. Complete *Mollicutes* 16S rRNA phylogenetic tree. The tree was obtained from a maximum likelihood phylogenetic analysis on the 16S rRNA gene, rooted with a *Bacillus cereus* sequence (NR_074540.1), and contains 230 leaves of which 88 are 16S rRNA amplicon sequences from this study, four are amplicon sequences from another firefly microbiome study (Green et al. 2021), and four are amplicon sequences from another study on insect-associated *Mollicutes* (Funaro et al. 2011). All the other leaves are full or near-full length 16S rRNA gene sequences and were obtained either through NCBI searches to acquire reference sequence of mollicute species or through NCBI BLASTn runs to find the most similar sequences to our amplicon sequences. Red branches contain bacterial ASVs sampled from fireflies (from the present and previous studies). ASV labels in bold are among the most abundant in our dataset (SV identifier number lower than 151). Gray triangles correspond to collapsed monophyletic groups containing more than two non-firefly associated strains. Genus and species names are from Gupta et al. (2019). Genera with dark gray fonts are commonly sampled from fireflies. The tree scale represents the number of nucleotide substitutions per site. All bootstrap values are shown on the tree. **B**. Sequence dissimilarities among firefly-associated ASVs from the *Mollicutes*. The Templeton, Crandall, and Sing (TCS) network was produced via the PopART software using an alignment of all firefly-derived *Mollicutes* ASV sequences with an SV identifier below 1,500 (i.e., among the 1,500 most abundant ASVs in our dataset). Hatch marks between nodes represent the number of mutations between two ASVs. Population genetics statistics are shown on the upper right of the figure. Identical ASVs (when considering only overlaps across sequences that were obtained from the two sequencing strategies) are shown on the lower left of the figure.

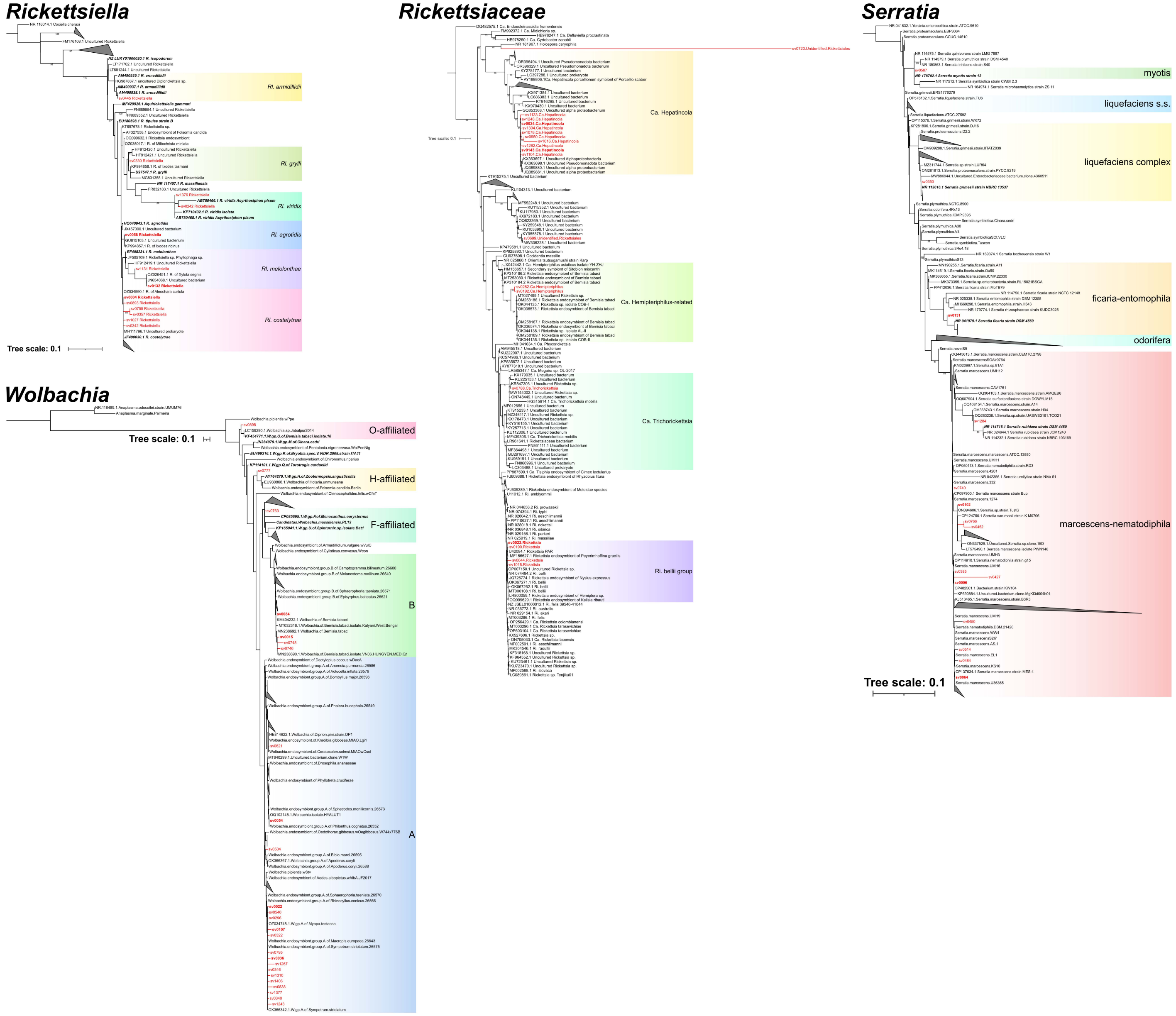


**Figure S10: Phylogenetic trees of four bacterial groups important to insects including sequences from fireflies**. All trees were obtained from a maximum likelihood phylogenetic analysis on the 16S rRNA gene except for the *Wolbachia* tree which was produced through a multi-locus phylogeny with fragment-insertion of 16S rRNA amplicon sequencing variants (ASVs) from our sequencing. For all trees, red branches contain bacterial ASVs sampled from fireflies (from the present study). ASV labels in bold are among the most abundant in our dataset (SV identifier number lower than 151). For the *Rickettsiella*, the *Rickettsiaceae*, and the *Serratia* trees, all the leaves that are not labeled as “sv” are full, near-full length, or on a few occasions, fragments of the 16S rRNA gene sequences that were obtained either through NCBI searches to acquire reference sequence of bacterial species or through NCBI BLASTn runs to find the most similar sequences to our amplicon sequences. The *Rickettsiella* tree was rooted with a *Coxiella cheraxi* sequence (NR_116014.1) and contains 63 leaves of which 13 are 16S rRNA amplicon sequences from this study. The *Rickettsiaceae* tree was rooted with a *Candidatus* Endoecteinascidia sequence (DQ482575.1) and contains 163 leaves of which 19 are 16S rRNA amplicon sequences from this study, eight from the *Ca.* Hepatincola genus, four from the *Rickettsia* genus, two from the *Ca.* Hemipteriphilus genus, one from the *Ca.* Trichorickettsia genus, and two unidentified *Rickettsiales*. Note that *Ca.* Hepatincola is not always classified as from the *Rickettsiaceae* family but was through our classifications using the SILVA database. The *Serratia* tree was rooted with a *Yersinia*

*enterocolitica* sequence (NR_041832.1) and contains 161 leaves of which 15 are 16S rRNA amplicon sequences from this study. For the *Wolbachia* tree, a backbone tree containing 160 leaves was first built from a maximum likelihood phylogenetic analysis based on a MAFFT alignment of seven genes (*coxA*, *fbpA*, *ftsZ*, *gatB*, *hcpA*, *recA*, and the 16S rRNA gene) from *Wolbachia* and *Anaplasma* (outgroup) genomes (backbone tree available in our Data Repository). Our 16S rRNA amplicon sequences from abundant ASVs in our study were then inserted into the backbone tree which was rooted at the two *Anaplasma* leaves (one from multi-locus another one from just the 16S rRNA gene (NR_118489.1)). The final tree contains 220 leaves of which 25 are 16S rRNA amplicon sequences from this study. The other leaves that were not part of the multi-locus, backbone tree are full or near-full length 16S rRNA gene sequences which were obtained through the same process as for the other trees. For all trees, gray triangles correspond to collapsed monophyletic groups containing more than two non-firefly associated strains. The tree scales represent the number of nucleotide substitutions per site. All bootstrap values are shown on the *Rickettsiella* tree, only bootstrap values above 70 are shown for the *Rickettsia* and *Serratia* trees, and for the *Wolbachia* tree, no bootstrap values were available for fragment-insertion trees (bootstrap support displayed on the backbone tree available in our Data Repository).

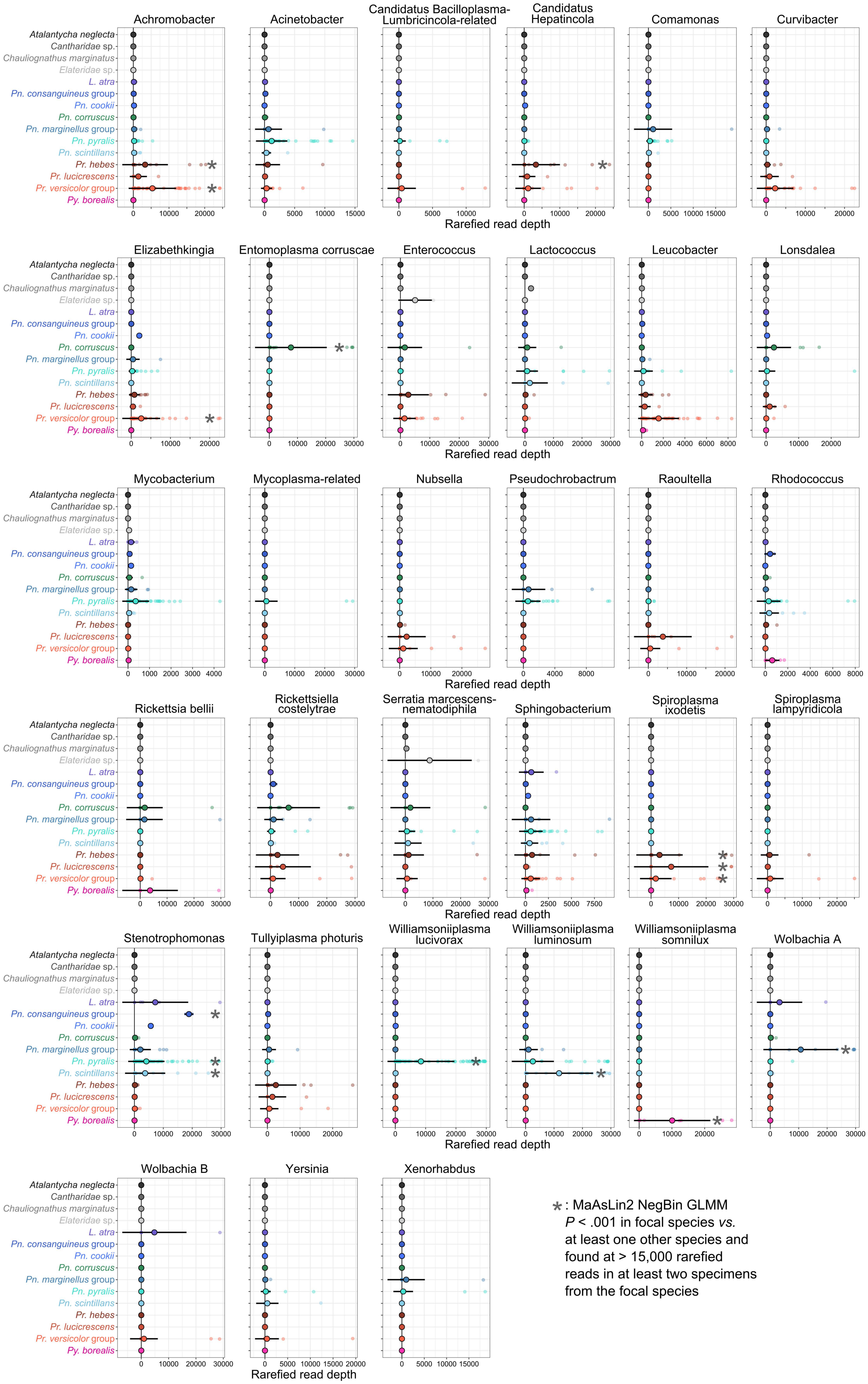

Achromobacter
Acinetobacter
Candidatus Bacilloplasma-Lumbricincola-related
Candidatus Hepatincola
Comamonas
Curvibacter
Elizabethkingia
Entomoplasma corruscae
Enterococcus
Lactococcus
Leucobacter
Lonsdalea
Mycobacterium
Mycoplasma-related
Nubsella
Pseudochrobactrum
Raoultella
Rhodococcus
Rickettsia bellii
Rickettsiella costelytrae
Serratia marcescens-nematodiphila
Sphingobacterium
Spiroplasma ixodetis
Spiroplasma lampyridicola
Stenotrophomonas
Tullyiplasma photuris
Williamsoniiplasma lucivorax
Williamsoniiplasma luminosum
Williamsoniiplasma somnilux
Wolbachia A
Wolbachia B
Yersinia
Xenorhabdus
Atalantycha neglecta
Cantharidae sp.
Chauliognathus marginatus
Elateridae sp.
L. atra
Pn. consanguineus group
Pn. cookii
Pn. corruscus
Pn. marginellus group
Pn. pyralis
Pn. scintillans
Pr. hebes
Pr. lucicrescens
Pr. versicolor group
Py. borealis
Rarefied read depth
*: MaAsLin2 NegBin GLMM
P < .001 in focal species vs.
at least one other species and
found at > 15,000 rarefied
reads in at least two specimens
from the focal species

**Figure S11: Abundance of 33 bacterial taxa across firefly species**. These bacteria are among the most abundant and prevalent in fireflies from our dataset. For each of them, rarefied read depth (at 30,000 reads) is shown on the x-axis. Firefly species are on the y-axis. Balls and error bars represent the per species average and standard deviation, respectively. The asterisks indicate when a bacterium was considered as a characteristic bacterial taxon in a given firefly species. To account for prevalence, abundance, and other random effects, we classified bacterial taxa as characteristic of a firefly species if (1) *MaAsLin2* multiple generalized linear mixed models indicated that they were significantly more abundant in a given species than in at least another one with a $P < 0.001$, and (2) they were found with an abundance of over 15,000 rarefied read depth in at least two specimens from the focal species. A summary of the *MaAsLin2* results is available in Table S5.

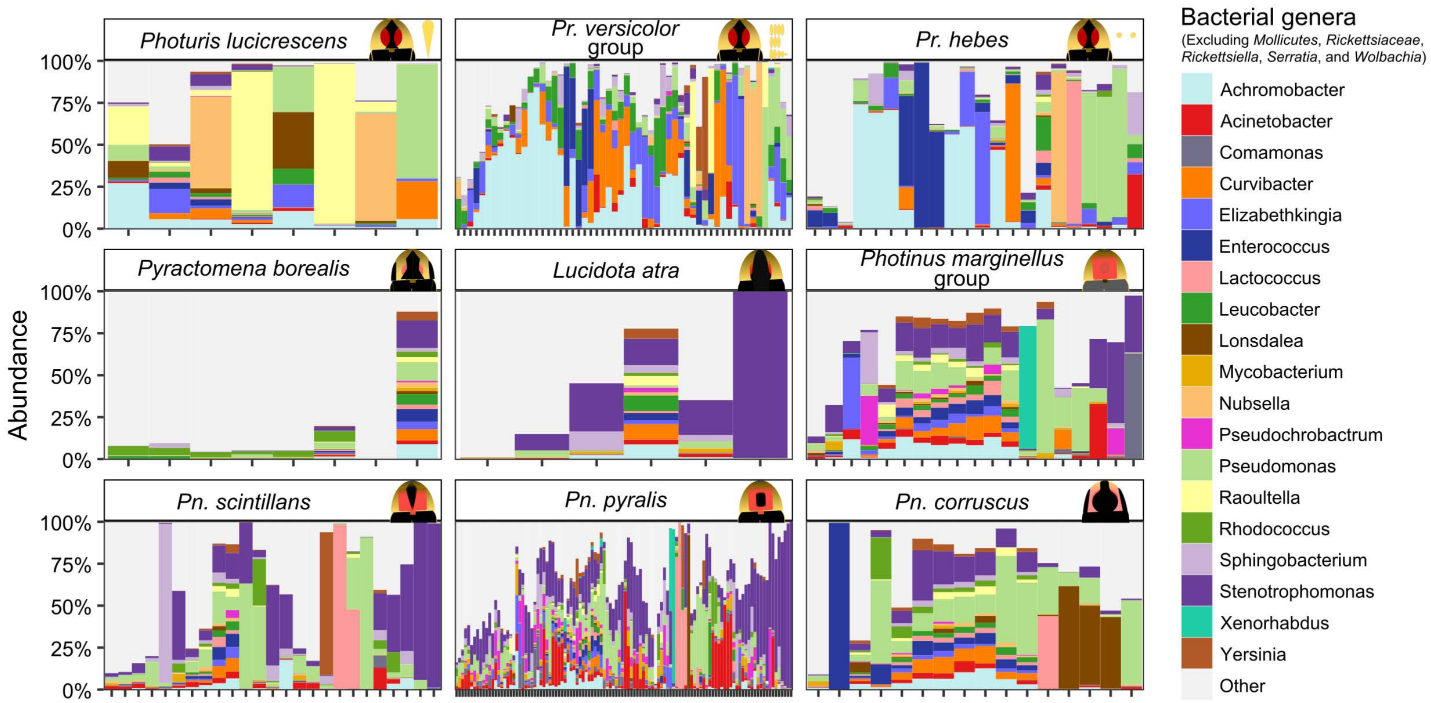


**Figure S12: Composition of bacterial taxa across firefly species after excluding mollicutes and putative endosymbionts**. Only firefly species with more than three specimens in our dataset post rarefying are shown. The relative rarefied abundance of the most abundant bacteria is shown as different colors whereas the abundance of all other bacteria is in light grey. Bacteria classified as from the *Candidatus* Hepatincola, *Mollicutes*, *Rickettsia*, *Rickettsiella*, *Serratia*, and *Wolbachia* were excluded from the dataset before generating these graphs. Bacterial colors here approximately match those in Figure 6 and Figure S15.

#### *Photinus corruscus*

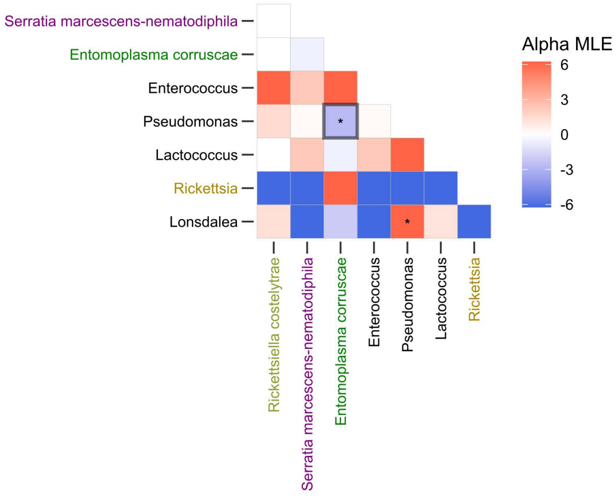


Filtered taxa with rarefied abundance > 10,000 reads in one or more samples
Presence = bacterial taxon with at least 1% relative abundance in a specimen

□ : result supported by coocur R package analysis

#### *Photinus marginellus* group

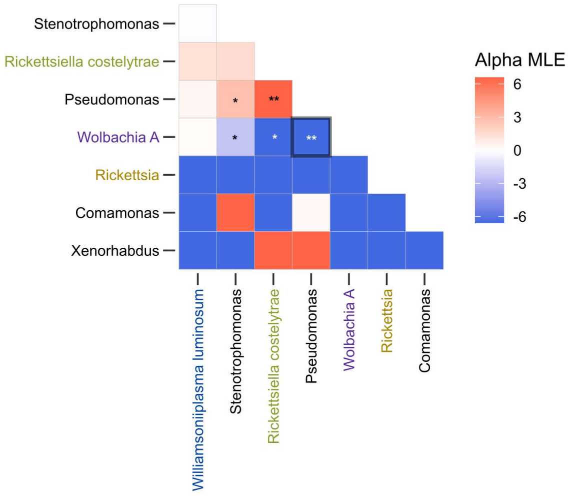


Filtered taxa with rarefied abundance > 10,000 reads in one or more samples
Presence = bacterial taxon with at least 1% relative abundance in a specimen

□ : result supported by coocur R package analysis

#### *Photinus pyralis*

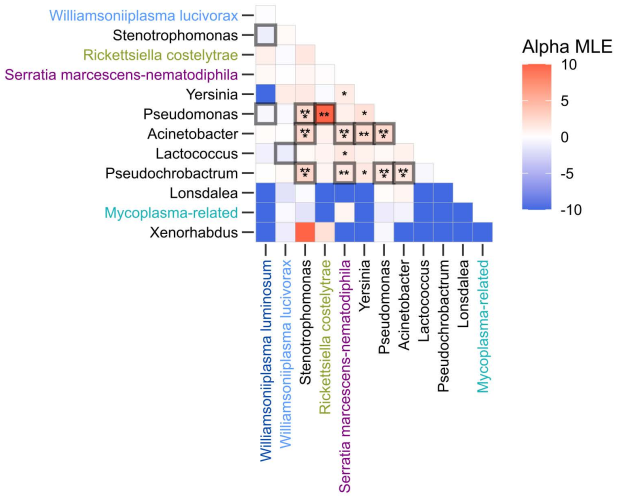


Filtered taxa with rarefied abundance > 10,000 reads in one or more samples
Presence = bacterial taxon with at least 1% relative abundance in a specimen

□ : result supported by coocur R package analysis

#### *Photinus scintillans*

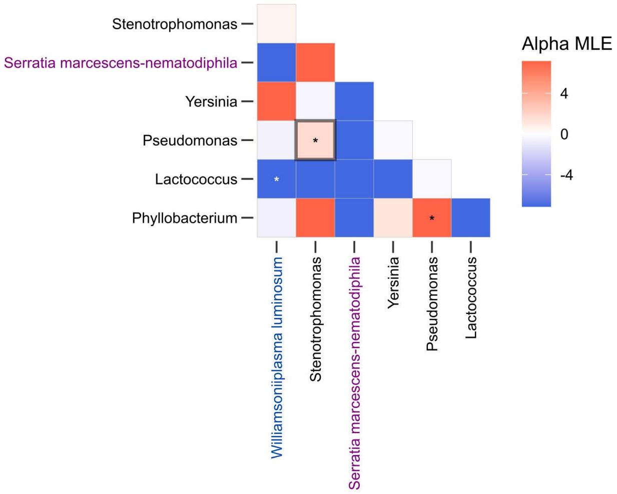


Filtered taxa with rarefied abundance > 10,000 reads in one or more samples
Presence = bacterial taxon with at least 1% relative abundance in a specimen

□ : result supported by coocur R package analysis

#### *Photuris*

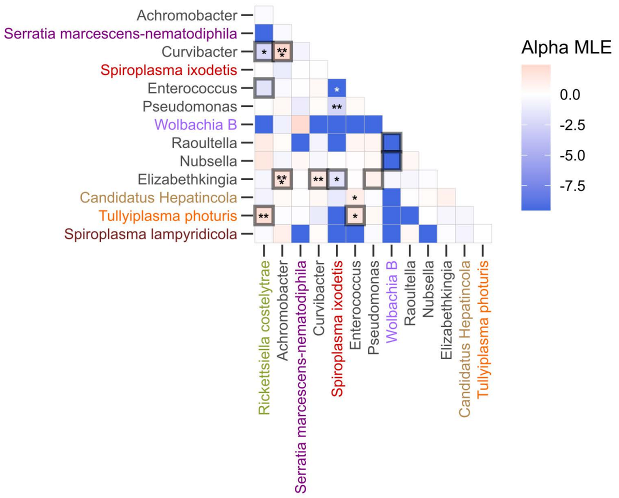


Filtered taxa with rarefied abundance > 20,000 reads in one or more samples
Presence = bacterial taxon with at least 1% relative abundance in a specimen

□ : result supported by coocur R package analysis

**Figure S13: Positive, negative, and random co-occurrence among bacterial taxa associated with fireflies**. The heatmaps were generated with the *CooccurrenceAffinity* R package. Red colors correspond to positive co-occurrence, blue color is for negative co-occurrence, and white for random co-occurrence. The darkest colors, very high or very low alpha MLE values (see Mainali et al., 2022) are for bacterial taxa that almost always or never co-occur, respectively. Dark borders indicate a positive or negative co-occurrence pattern supported by inferences from the *cooccur* R package. Asterisks correspond to different levels of statistical significance in the positive or negative co-occurrence obtained from *CooccurrenceAffinity*, with ".": $P < 0.1$, "*": $P < 0.05$, "**": $P < 0.01$, "***": $P < 0.001$.

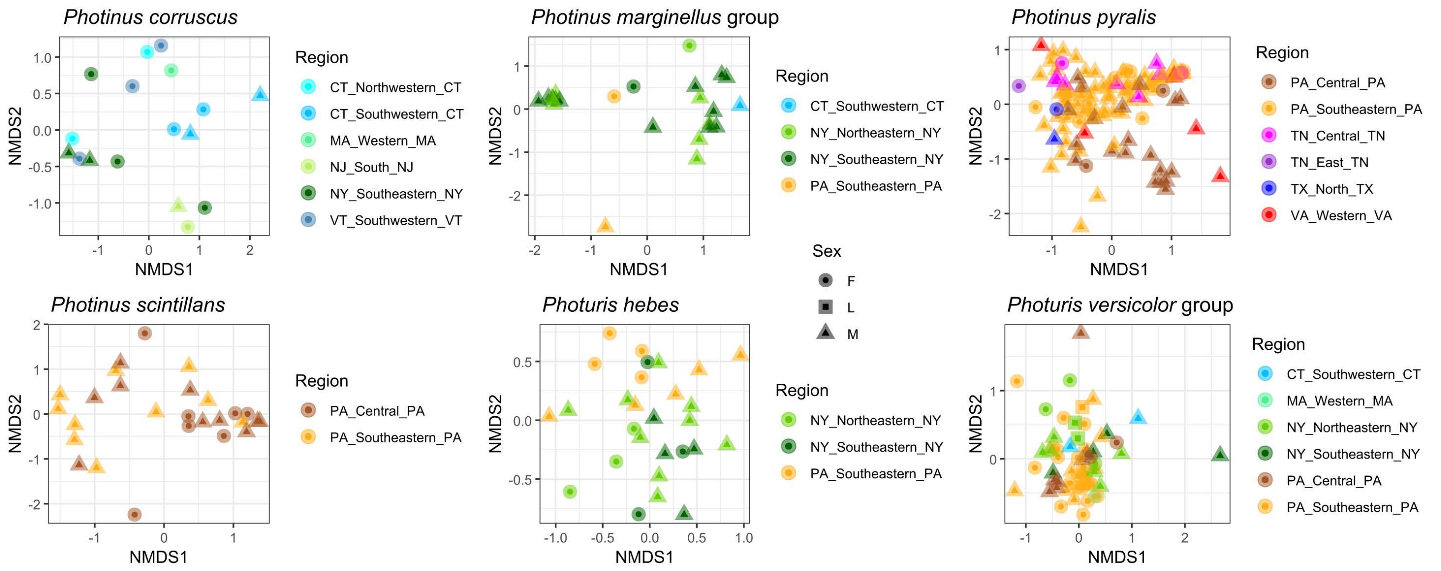


**Figure S14: Microbiome divergence across fireflies from different regions**. Non-metric multidimensional scaling (NMDS) analyses showing the separation of the bacterial communities based on composition and relative abundance across regions for four firefly species and two species groups. All plots are derived from Bray-Curtis distances calculated on non-rarefied, compositional relative abundance of taxon-aggregated bacteria in firefly specimens. Results from *MaAsLin2* multiple generalized linear mixed models, inferring bacterial differential abundance, can be found in Table S6 and results from PERMANOVAs and pairwise PERMANOVAs comparing regional microbiomes per species can be found in Table S3.

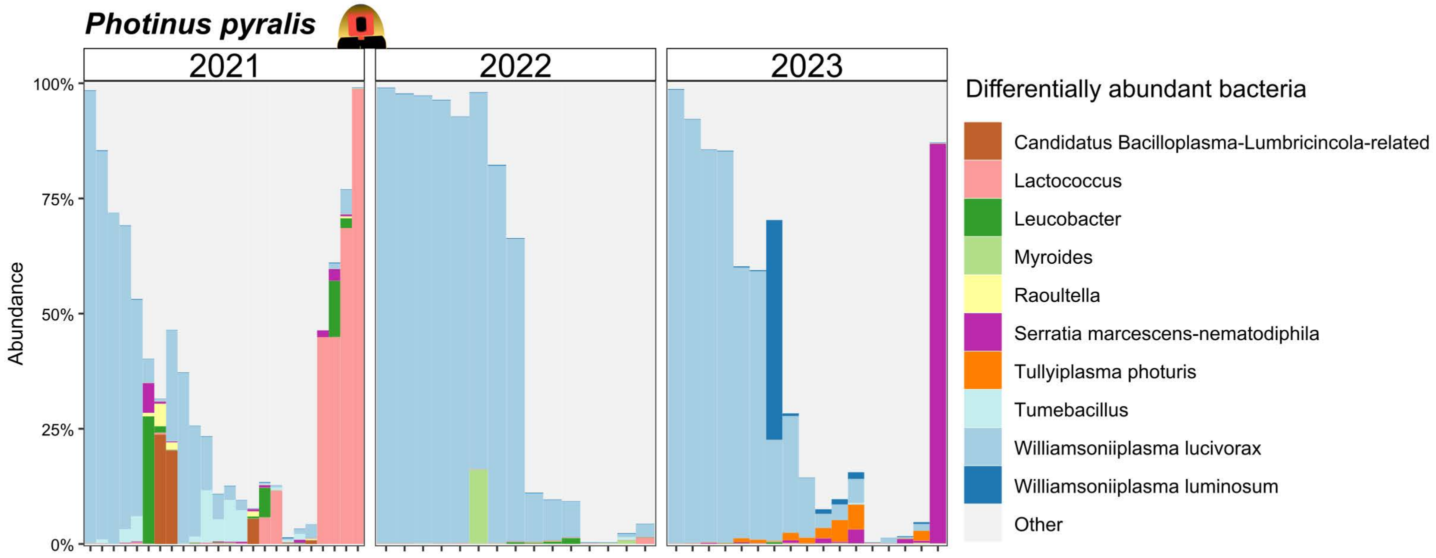


**Figure S15: Most differentially abundant bacteria across the years in a firefly species collected in a single location**. *Photinus pyralis* fireflies were collected in Philadelphia Fairmont Park in the month of July 2021, 2022, and 2023. Bacterial colors here approximately match those in Figure 6 and Figure S12.

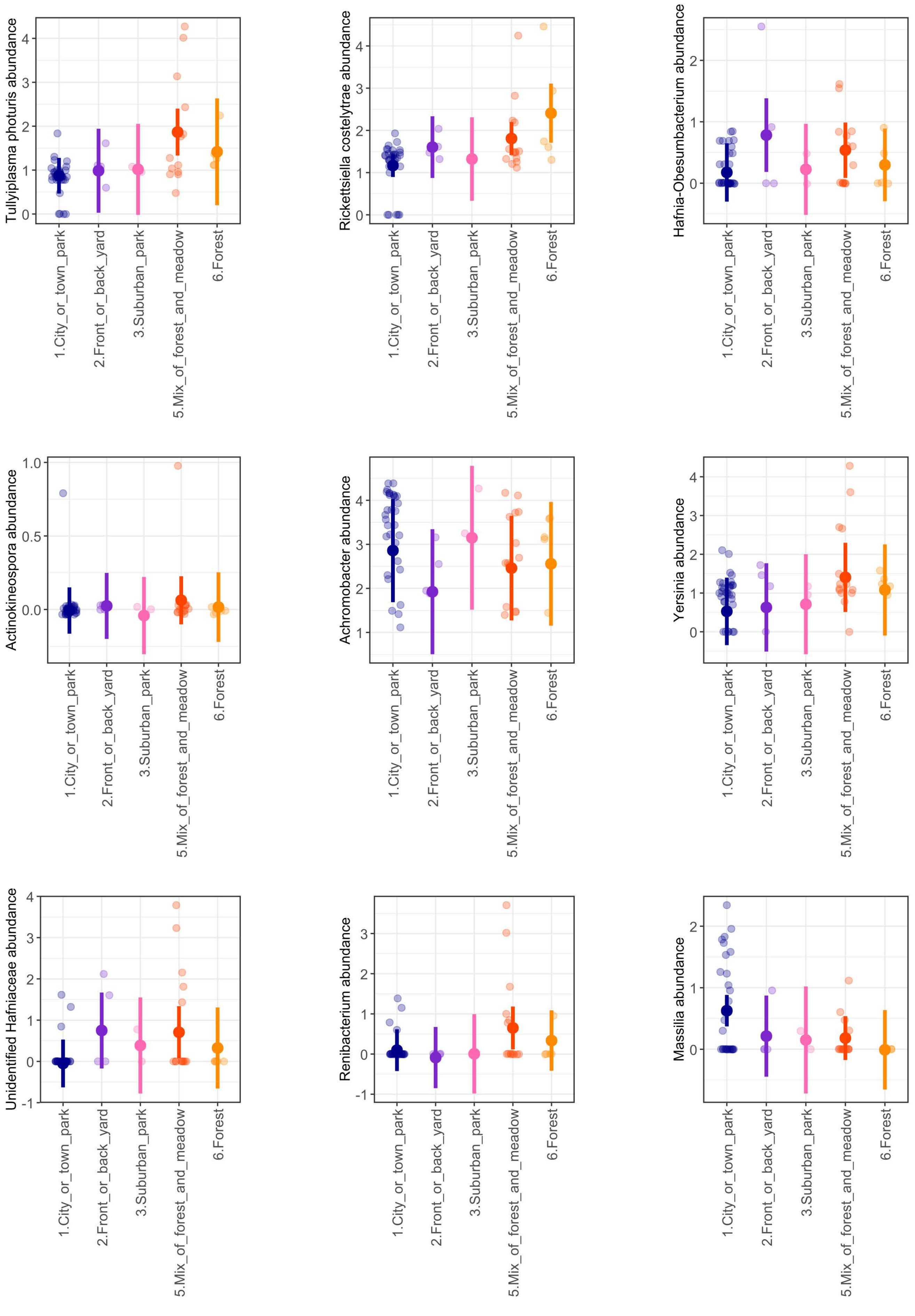
Tullyiplasma photuris abundance
Rickettsiella costelytrae abundance
Hafnia-Obesumbacterium abundance
Actinokineospora abundance
Achromobacter abundance
Yersinia abundance
Unidentified Hafniaceae abundance
Renibacterium abundance
Massilia abundance
1.City_or_town_park
2.Front_or_back_yard
3.Suburban_park
5.Mix_of_forest_and_meadow
6.Forest

**Figure S16: Significantly differentially abundant bacterial taxa across habitat types in unidentified species from the *Photuris versicolor* group**. These bacteria are among those with the most significantly different abundance across habitat in this firefly species group, as assessed through *MaAsLin2* multiple generalized linear mixed models (GLMMs). For each of these bacteria, a univariate GLMM was run to test whether its rarefied abundance differs across these fireflies' habitats, when controlling for sex type, region, day of year, and year of capture. The most consistent results were observed for *Rickettsiella costelytrae*. Results from *MaAsLin2* runs can be found in Table S7. Bacterial abundance is expressed as $\log_{10}$ of rarefied abundance.

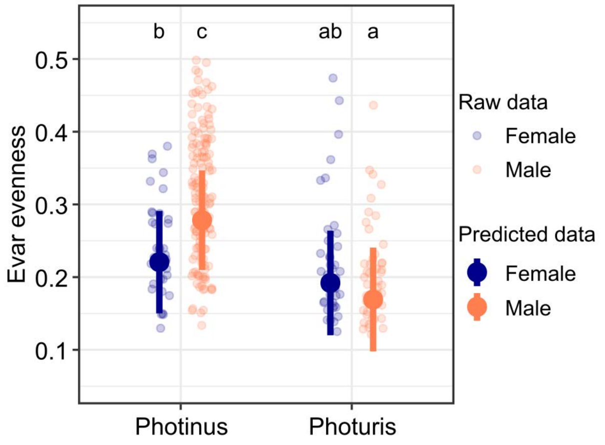


**Figure S17: Male *Photinus* microbiomes are more even than females'**. A univariate generalized linear mixed model was used to test whether the index of Evar evenness is different between females and males in *Photinus* and *Photuris* fireflies, when controlling for sampling location, day of year, year of capture, and sequencing type. Significant differences are shown with lowercase English letters.

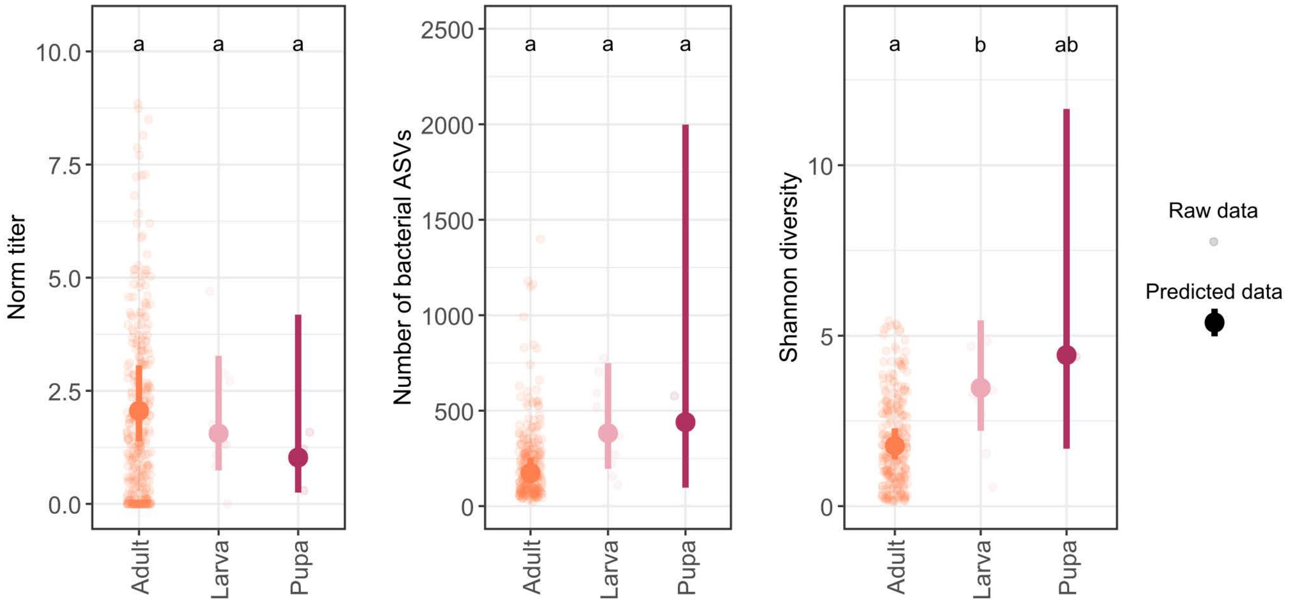


**Figure S18: Bacterial absolute abundance and diversity in adults, larvae, and pupae**. This figure is similar to Figure 7F, except that the juvenile category was broken down into larvae and pupae. Univariate generalized linear mixed models (GLMMs) were used to test whether bacterial titer, bacterial richness, and the Shannon diversity index are different between adults, larvae, and pupae, using firefly genus as a covariate and controlling for region, day of year, and year of capture. Bacterial titer is expressed as in Figure 1C. Predicted data with 95% confidence intervals were obtained from the GLMMs.

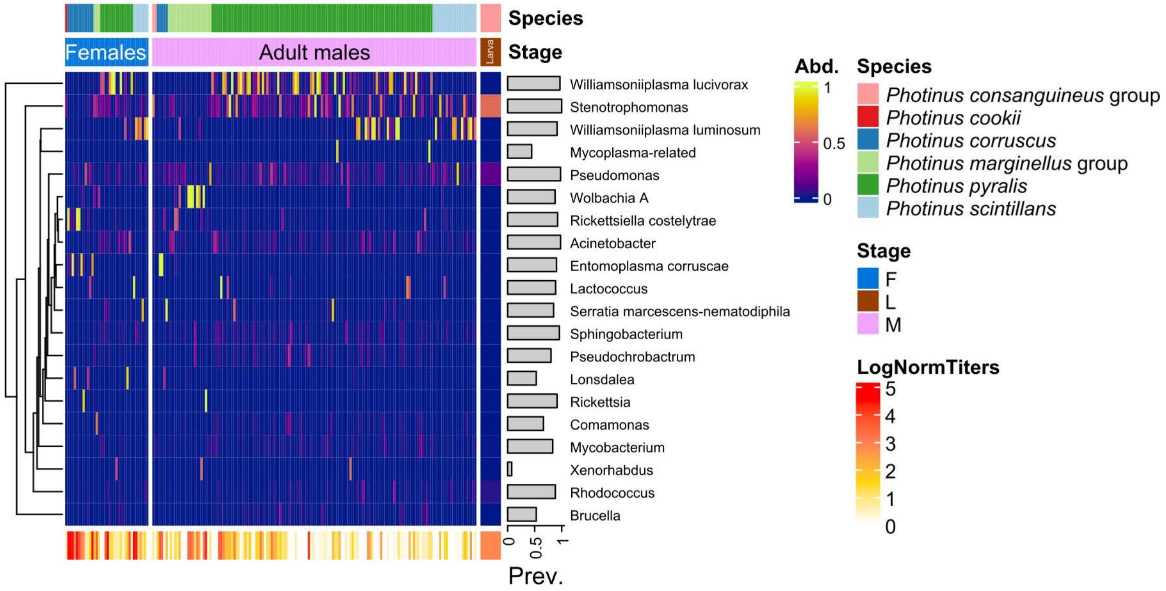
Species
Females
Adult males
Larva
Stage
Williamsoniiplasma lucivorax
Stenotrophomonas
Williamsoniiplasma luminosum
Mycoplasma-related
Pseudomonas
Wolbachia A
Rickettsiella costelytrae
Acinetobacter
Entomoplasma corruscae
Lactococcus
Serratia marcescens-nematodiphila
Sphingobacterium
Pseudochrobactrum
Lonsdalea
Rickettsia
Comamonas
Mycobacterium
Xenorhabdus
Rhodococcus
Brucella
0
0.5
1
Prev.
Abd.
1
0.5
0
Species
Photinus consanguineus group
Photinus cookii
Photinus corruscus
Photinus marginellus group
Photinus pyralis
Photinus scintillans
Stage
F
L
M
LogNormTiters
5
4
3
2
1
0

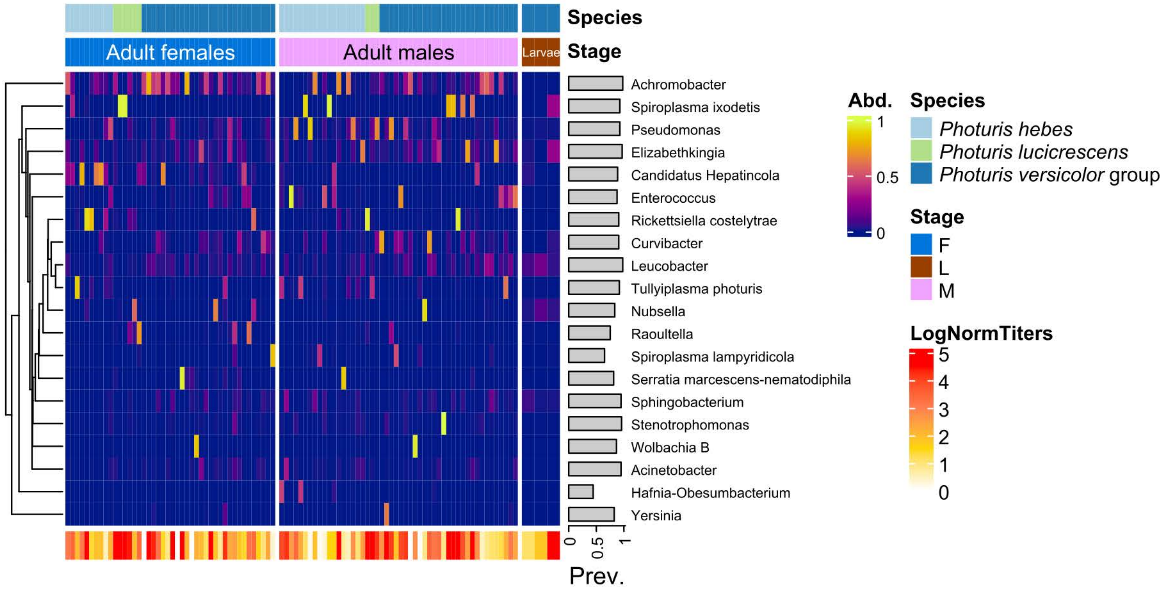
Species
Adult females
Adult males
Larvae
Stage
Achromobacter
Spiroplasma ixodetis
Pseudomonas
Elizabethkingia
Candidatus Hepatincola
Enterococcus
Rickettsiella costelytrae
Curvibacter
Leucobacter
Tullyiplasma photuris
Nubsella
Raoultella
Spiroplasma lampyridicola
Serratia marcescens-nematodiphila
Sphingobacterium
Stenotrophomonas
Wolbachia B
Acinetobacter
Hafnia-Obesumbacterium
Yersinia
0
0.5
1
Prev.
Abd.
1
0.5
0
Species
Photuris hebes
Photuris lucicrescens
Photuris versicolor group
Stage
F
L
M
LogNormTiters
5
4
3
2
1
0

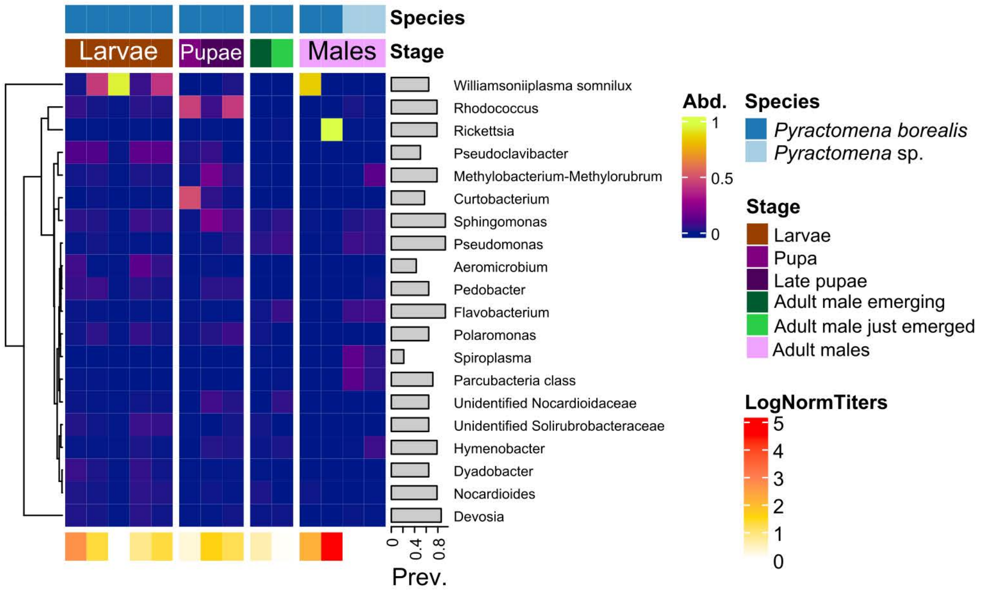
Species
Larvae
Pupae
Males
Stage
Williamsoniiplasma somnilux
Rhodococcus
Rickettsia
Pseudoclavibacter
Methylobacterium-Methylorubrum
Curtobacterium
Sphingomonas
Pseudomonas
Aeromicrobium
Pedobacter
Flavobacterium
Polaromonas
Spiroplasma
Parcubacteria class
Unidentified Nocardioidaceae
Unidentified Solirubrobacteraceae
Hymenobacter
Dyadobacter
Nocardioides
Devosia
0
0.4
0.8
Prev.
Abd.
1
0.5
0
Species
Pyractomena borealis
Pyractomena sp.
Stage
Larvae
Pupa
Late pupae
Adult male emerging
Adult male just emerged
Adult males
LogNormTiters
5
4
3
2
1
0

**Figure S19: Bacterial composition in larvae and adults of three firefly species**. The composite heatmaps show bacterial composition and relative abundance for the *Photinus*, *Photuris*, and *Pyractomena* genera. Relative abundance values were obtained from compositional transformation of non-rarefied read depth, at the "taxon" level, along with total absolute bacterial abundance per specimen (expressed as normalized bacterial titer, see Figure 1C) and prevalence of single bacteria across these firefly specimens.

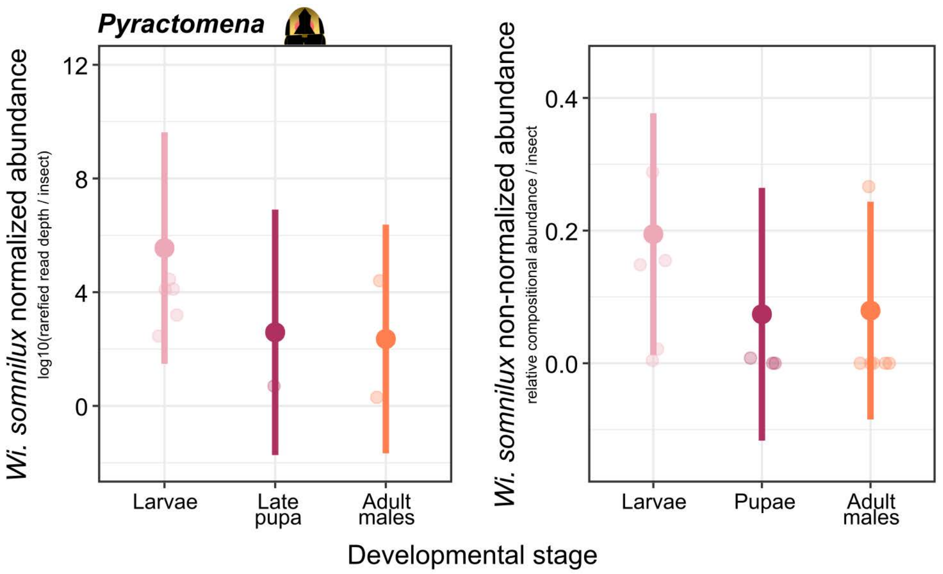


**Figure S20: Abundance of *Williamsoniiplasma somnilux* in *Pyractomena* fireflies is higher in larvae than in adults and pupae**. *Wi. somnilux* abundance is expressed as the $\log_{10}$ of rarefied read depth in the first panel and as the relative compositional non-rarefied abundance in the second panel. A univariate generalized linear mixed modelx was used to test whether *Wi. somnilux* abundance is influenced by developmental stage in *Pyractomena* fireflies, when controlling for species, region, day of year, and year of collection.

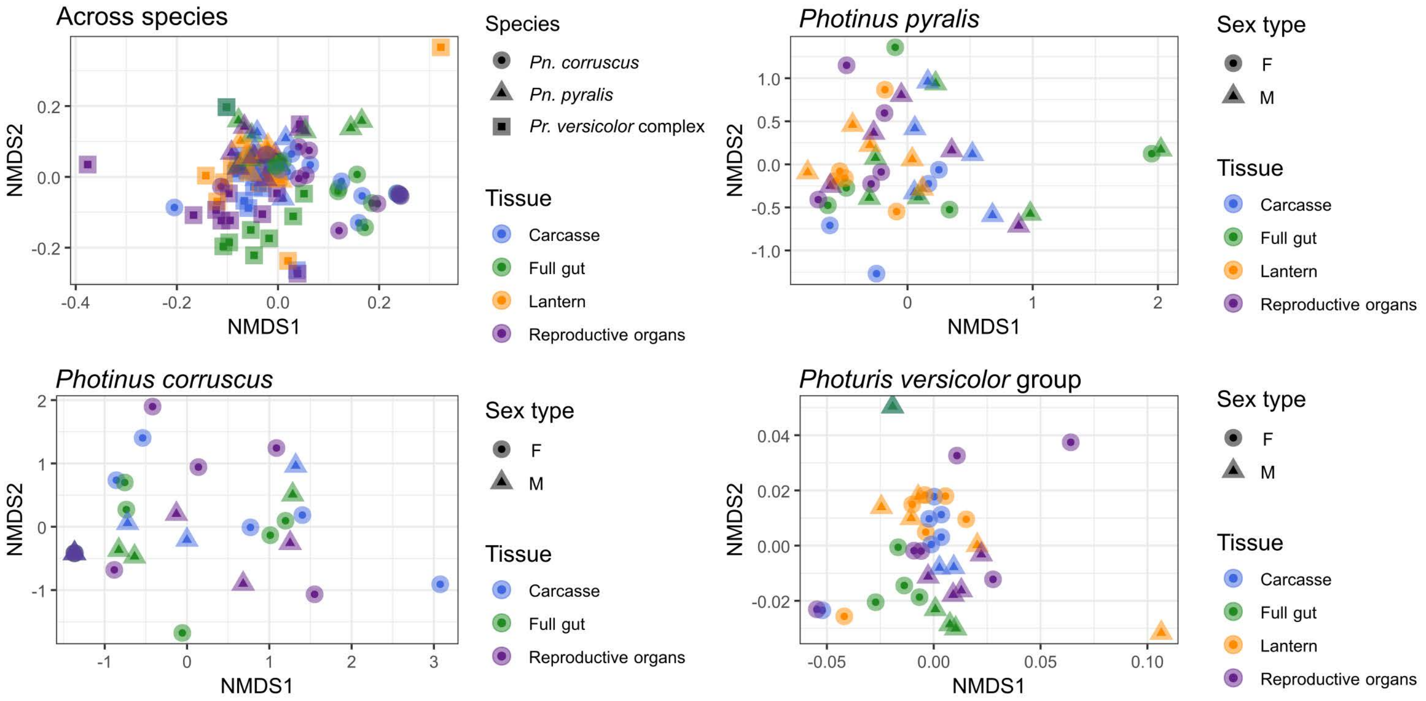


**Figure S21: Microbiome divergence across firefly tissues**. Non-metric multidimensional scaling (NMDS) analyses showing the separation of the bacterial communities based on composition and relative abundance across tissues for three firefly species and species group. All plots are derived from Bray-Curtis distances calculated on non-rarefied, compositional relative abundance (i.e., "proportional abundance") of taxon-aggregated bacteria in firefly specimens. One outlier was removed from the first and second graphs.

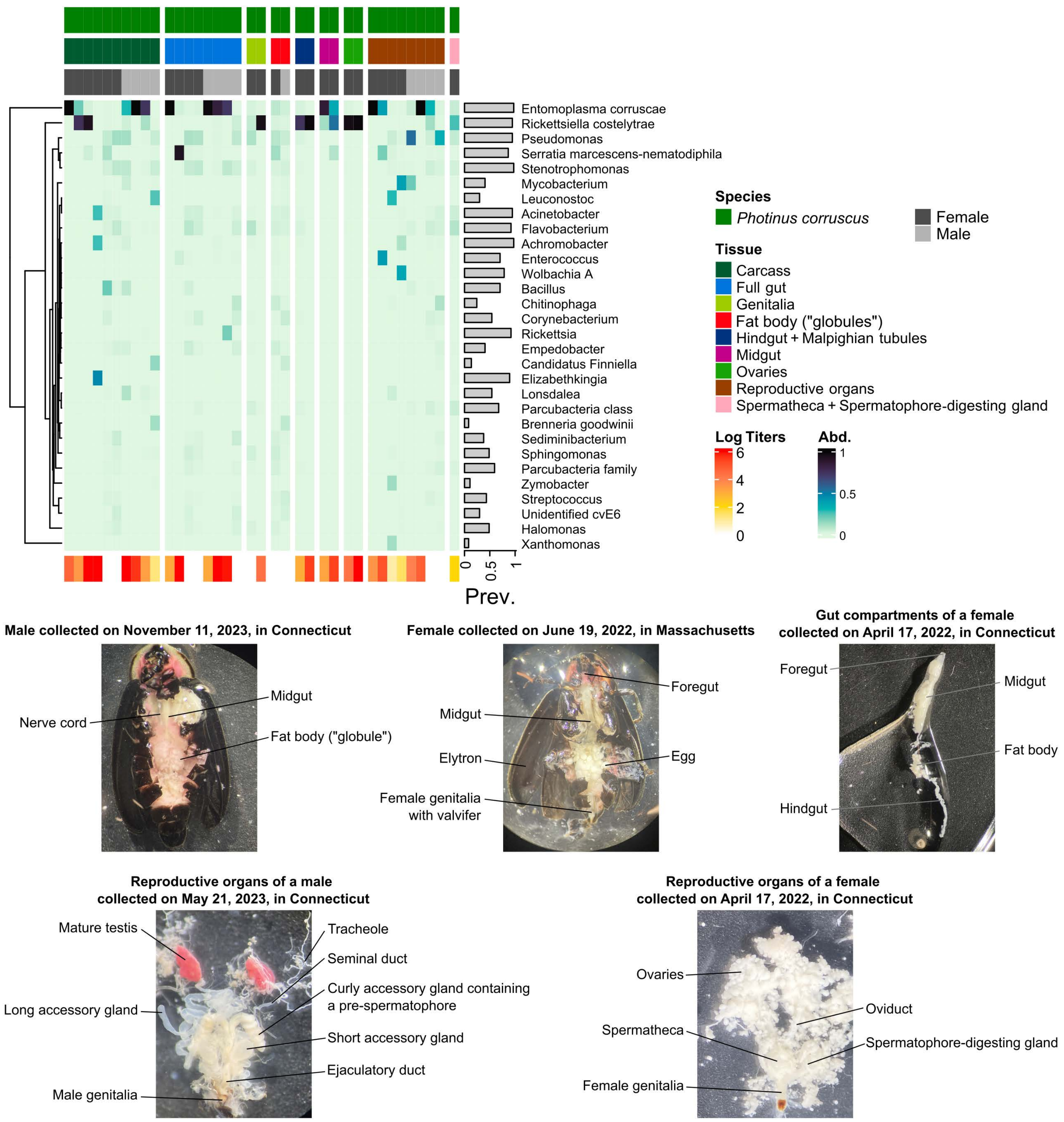


**Figure S22: Bacterial composition in different tissues of *Photinus corruscus* fireflies**. The composite heatmap at the top shows bacterial composition and relative abundance, obtained from compositional transformation of non-rarefied read depth, at the "taxon" level, along with total absolute bacterial abundance per specimen (expressed as normalized bacterial titer, see Figure 1C) and prevalence of single bacteria across specimen tissues. Photographs of dissections with anatomical annotations are shown at the bottom.

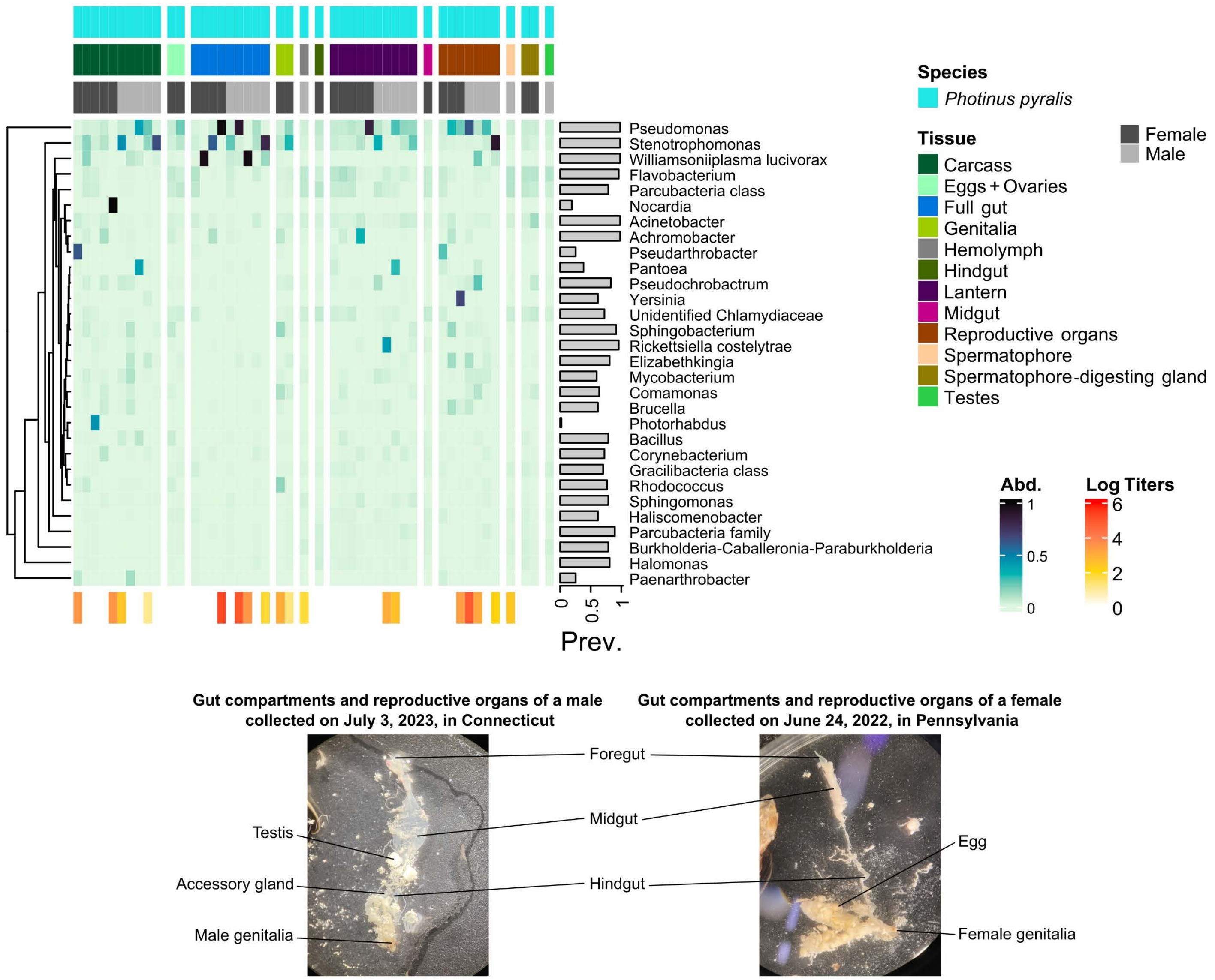


**Figure S23: Bacterial composition in different tissues of *Photinus pyralis* fireflies**. The composite heatmap at the top shows bacterial composition and relative abundance, obtained from compositional transformation of non-rarefied read depth, at the "taxon" level, along with total absolute bacterial abundance per specimen (expressed as normalized bacterial titer, see Figure 1C) and prevalence of single bacteria across specimen tissues. Photographs of dissected organs with anatomical annotations are shown at the bottom.

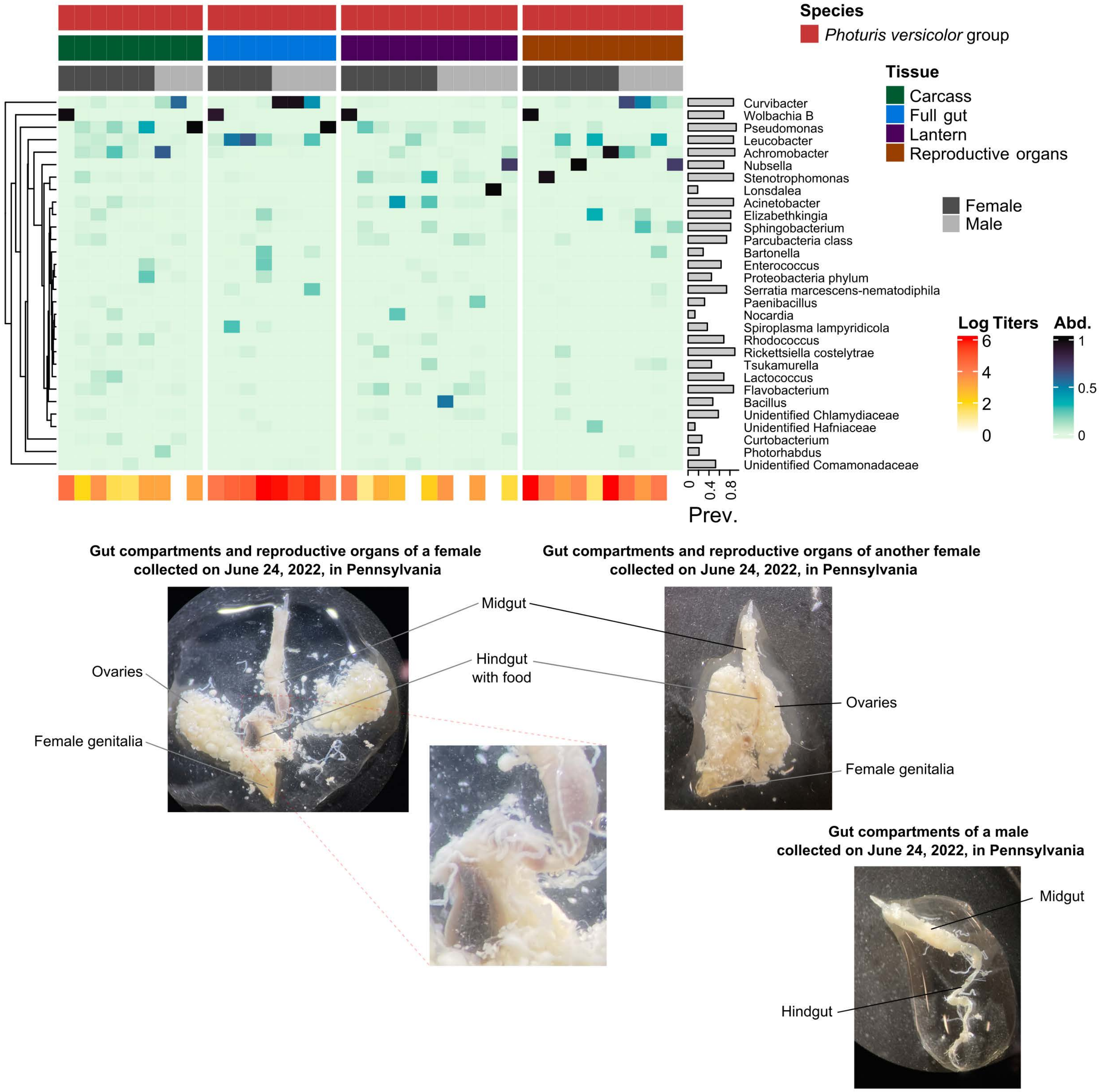


**Figure S24: Bacterial composition in different tissues of unidentified species from the *Photuris versicolor* group**. The composite heatmap at the top shows bacterial composition and relative abundance, obtained from compositional transformation of non-rarefied read depth, at the "taxon" level, along with total absolute bacterial abundance per specimen (expressed as normalized bacterial titer, see Figure 1C) and prevalence of single bacteria across specimen tissues. Photographs of mouth parts and dissections with anatomical annotations are shown at the bottom.
