## Supplementary Text 1 for "Species-specific bacterial associations emerge from stochastically assembled microbiomes in Northeastern American fireflies"

**Table of Content**

### Methods

#### Firefly DNA barcoding.

We used barcoding to improve our identification of *Photinus* species and species groups and juveniles. Separating the *Photinus consanguineus* group, *Pn. marginellus* group, and *Pn. scintillans* was almost impossible solely based on morphology and flash patterns (aedagus dissection was not considered since it would have impacted our microbiome sequencing and analyses). A large number of the barcoded specimens (*n* = 27) were found to be from the *Pn. scintillans* species. In addition to these *Photinus* and juvenile identifications, we added one representative firefly specimen for each species or species group for which we had confidence in our preliminary identification to assess the robustness of the COI DNA barcoding method.

We first used, for 59 specimens, the LCO1490 (5’-GGTCAACAAATCATAAAGATATTGG-3’) and HCO2198 (5’-TAAACTTCAGGGTGACCAAAAAATCA-3’) primers (Folmer et al., 1994). For 43 specimens that could not be amplified with these first primers, we then used the C1-J-1718m (5’-GGAGGCTTCGGAAATTGATTAGTTCC-3’) and C1-J-1751ff (5’-GGGGCTCCTGATATAGCTTTTCC-3’) as forward primers, and C1-J-2183 (5’-CCAAAAAATCAAAATAAATGTTG-3’) as the reverse primer from (Simon et al., 1994; Stanger-Hall et al., 2007). From these specimens’ DNA samples, 37 still did not amplify. We thus used the degenerate primers Sauron‐S878 (5’-GGDRCWGGWTGAACWGTWTAYCCNCC-3’) as the forward and jgHCO2198 (5’-TAIACYTCIGGRTGICCRAARAAYCA-3’) as the reverse primer, designed by (Geller et al., 2013; Rennstam Rubbmark et al., 2018).

Following Sanger sequencing of our PCR products from Eurofins (Lancaster, PA, USA), we cleaned and combined forward and reverse sequences using the Geneious Prime software version 2024.0.2 (Biomatters, New Zealand), ran multiple sequence alignment with other firefly COI sequences downloaded from NCBI using the Clustal Omega aligner (Sievers et al., 2011), and performed maximum likelihood phylogenetic analyses with SeaView v5.0.5 (Gouy et al., 2009).

PCR recipes and thermocycler conditions for COI barcoding were as follows. For the first PCR using the LCO1480–HCO2198 primers, we prepared PCR mixes of 25 µL total per sample, combining 2.5 µL of DNA template with 7.07 µL of sterile water, 2.14 µL of the LCO1480 primer (5 µM), 2.14 µL of the HCO2198 primer (5 µM), 0.17 µL of OneTaq DNA polymerase (5 U/μL; New England Biolabs, Ipswich, MA, USA), 9.23 µL of 5× OneTaq Standard Reaction Buffer (NEB), 1.75 µL of dNTPs (10 mM), and from 0 to 0.60 µL (in a volume gradient, adjusting the volume of sterile water accordingly) of MgCl_2_ (25 mM; NEB). The thermocycling program consisted of an initial denaturation at 94 °C (3 minutes), followed by 35 cycles at 95 °C (denaturation, 30 seconds), 50–52 °C (annealing, 30 seconds), 72 °C (extension, 1 minute), and a final extension step at 72 °C for 1 minute.

For the second and third PCR runs, using the C1-J-1718m/C1-J-2183 and the C1-J-1751ff/C1-J-2183 primer pairs, we similarly prepared PCR mixes of 25 µL total per sample, combining 1.25 µL of DNA template with 11.40 µL of sterile water, 2.11 µL of the forward primer (5 µM; C1-J-1718m or C1-J-1751ff), 2.11 µL of the reverse primer (5 µM; C1-J-2183), 0.13 µL of OneTaq DNA polymerase (5 U/μL; NEB), 7.00 µL of 5× OneTaq Standard Reaction Buffer (NEB), and 1.00 µL of dNTPs (10 mM). The thermocycling program consisted of an initial denaturation at 95 °C (3 minutes), followed by 35 cycles at 95 °C (denaturation, 30 seconds), 50 °C (annealing, 1 minute), 72 °C (extension, 1 minute), and a final extension step at 72 °C for 1 minute.

For the fourth PCR run, using the Sauron‐S878 and jgHCO2198 primers, we prepared PCR mixes of 20 µL total per sample, combining 3 µL of DNA template with 5.44 µL of sterile water, 2.00 µL of the Sauron‐S878 primer (5 µM), 2.00 µL of the jgHCO2198 primer (5 µM), 0.10 µL of OneTaq DNA polymerase (5 U/μL; NEB), 2.00 µL of dNTPs (10 mM), 4.00 µL of 5× of OneTaq Standard Reaction Buffer (NEB), 0.96 µL of MgCl_2_ (25 mM; NEB), and 0.50 µL bovine serum albumin (10 mg/mL). The thermocycling program consisted of an initial denaturation at 95 °C (3 minutes), followed by 35 cycles at 95 °C (denaturation, 20 seconds), 50 °C (annealing, 30 seconds), 72 °C (extension, 1 minute), and a final extension step at 72 °C for 3 minutes.

All PCR reactions were run on a BioRad C1000 Touch Thermal Cycler (Bio-Rad Laboratories Inc., Hercules, CA, USA). For each PCR reaction, a blank negative control containing sterile water instead of template DNA was included. Gel electrophoresis was used to verify correct amplification through 1.5% agarose gels and mixing 4 µL of the PCR products with 1 µL of 20× Gel Red solution. DNA barcoding sequences were submitted to NCBI as GenBank accession IDs PV844642-PV844700.

#### Notes on firefly species identifications.

While two *Photuris* species from the *Photuris versicolor* group could be identified with certainty thanks to flash patterns and morphology (Faust, 2017; McDermott, 1967), all the others were kept at the species group level, and are referred to as “unidentified species from the *Photuris versicolor* group” in the main text. Similarly, related species within each of the *Photinus consanguineus* and *Pn. marginellus* groups can only be confidently separated into species through the examination of the aedeagus (but see Lloyd, (1967) for potential hybridization of *Pn. marginellus* and *Pn. curtatus* in some areas of their range). We thus use the “*Photinus consanguineus* group” and “*Photinus marginellus* group” terminologies, respectively. The identification of some *Pyractomena* species can also be difficult. Two *Pyractomena* specimens could not be identified with certainty to the species level, even with COI barcoding (grouping with *Py. palustris*; Figure S1), but they were both found to be very similar to *Py. sinuata* and an unnamed *Pyractomena* species, referred to as the “Hudson Valley Firefly” or “near *sinuata*” by Lloyd, (2018). We use the term “*Pyractomena* sp.” in the main text, but acknowledge that they may be from these rare species.

#### Bacterial titer quantification through quantitative PCR.

For each sample that was run in quantitative PCR (qPCR) reactions, 4 µL of template DNA was mixed with 16 µL of a qPCR master mix that contained 10 µL of PowerUp SYBR Green Master Mix (Applied Biosystems), 2 µL of each of the two primers at 5 µM concentration, and 2 µL of sterile PCR water. The qPCR program consisted of uracil-DNA glycosylases activation at 50 °C for 2 minutes; initial denaturation at 94 °C for 3 minutes; and 40 cycles of 94 °C for 45 seconds, 50 °C for 60 seconds, 72 °C for 60 seconds, and a plate read at elongation. Melting curve analysis was applied at the end of these 40 cycles, using the default Dissociation Stage method implemented by the 7300 Real-Time PCR System (rapid heating to 95 °C for 15 seconds, followed by cooling at 60 °C for 60 seconds, reheating at 95°C for 15 seconds, and recooling at 60 °C for 15 seconds).

To obtain absolute bacterial titers measured in 16S rRNA gene copy numbers, we used, in each qPCR plate, a standard curve built on a ten-fold dilution series of a DNA sample with known 16S rRNA gene copy numbers. To produce diluted standards, we extracted DNA from a Lep1A1 *Caballeronia* symbiont originally isolated from a *Leptoglossus zonatus* leaf-footed bug (Hunter et al., 2022). From this extracted DNA, we ran a conventional PCR using the aforementioned qPCR primers and PCR reagents, replacing SYBR Green with OneTaq DNA polymerase, OneTaq Reaction Buffer, and dNTPs (see barcoding section above). Following DNA quantification with Qubit and calculation of the total 16S rRNA gene copy numbers in PCR products, we performed ten-fold serial dilutions from 10^8^ to 10^1^ 16S rRNA gene copies per µL, which were used in the qPCR standard curves.

Bacterial titer values were corrected using blank samples. We first subtracted the titer in each sample by the titer in its corresponding DNA extraction blank. For dissected tissues, we also subtracted this corrected value per sample by its corresponding corrected dissection blank. Each sample was run in two or three separate technical replicates, consisting of runs on separate qPCR plates. PCR efficiency ranged from 51.1% to 94.3% per plate (average = 78.1%, standard deviation (SD) = 8.9%, median = 78.4%). We calculated the average of blank-corrected titers across plate replicates and used these values in downstream statistical analyses.

#### Illumina library preparation and sequencing.

For the first batch of amplicon sequencing, DNA was amplified with the universal bacterial primers 341F (5’-CCTACGGGNGGCWGCAG-3’) and 785R (5’-GACTACHVGGGTATCTAATCC-3’), focusing on the V3-V4 hypervariable region of the 16S rRNA gene (Klindworth et al., 2012). Each primer included an overhang adapter to enable downstream addition of sample indexes, as per Illumina’s 16S Metagenomic Sequencing Library Preparation guide. Samples were amplified in a volume of 20 µL with final concentrations of 0.5 µM forward primer, 0.5 µM reverse primer, and 1× Q5 Master Mix (NEB). The thermocycler program was: denaturation at 98 °C for 1 minute followed by 35 cycles of denaturation at 98 °C for 10 seconds, primer annealing at 58 °C for 20 seconds, and extension at 72 °C for 30 seconds, with a final extension of 72 °C for 2 minutes. We cleaned the PCR products with magnetic beads (Rohland & Reich, 2012). To differentiate the amplicons from each sample, a second short PCR was used to attach a unique pair of 8-nucleotide barcodes to the forward and reverse ends of each amplicon (Hamady et al., 2008). The volume of this second PCR was also 20 µL. We amplified 2.5 µL of the cleaned PCR product with 0.5 µM of each barcoded primer and 1× Q5 master mix. Thermocycler conditions were: denaturation at 98 °C for 1 minute followed by 8 cycles of denaturation at 98 °C for 10 seconds, primer annealing at 55 °C for 20 seconds, and extension at 72 °C for 30 seconds, followed by a final extension of 72 °C for 2 minutes. PCR products were again cleaned with magnetic beads, quantified fluorometrically with Qubit DS DNA HS assays (Invitrogen, Waltham, MA, USA), and an equal mass of each sample was bidirectionally sequenced on a 600 cycle paired-end run on an Illumina Mi-Seq platform (Illumina, San Diego, CA, USA).

For the second batch, PCR amplicon libraries targeting the 16S rRNA gene were first produced using a barcoded bacterial universal primer set adapted for Illumina sequencing (Caporaso et al., 2012). Specifically, the V4 region of the 16S rRNA gene was PCR amplified with primers 515F–806R that included sequencer adapter sequences used in the Illumina flowcell (1, 2, 3, 4, 5). The forward amplification primer also contained a twelve base pair barcode sequence (Caporaso et al., 2012; Walters et al., 2015) that supported pooling of up to 2,167 different samples in each lane (1, 2, 3, 4, 5). Each 25 µL PCR reaction contained 9.5 µL of PCR sterile water, 12.5 µL of AccuStart II PCR ToughMix (QuantaBio, Beverly, MA, USA), 1 µL Golay barcode tagged forward primer at 200 pM, 1 µL reverse primer at 200 pM, and 1 µL of template DNA. The conditions for PCR were as follows: 94 °C for 3 minutes to denature the DNA, with 35 cycles at 94 °C for 45 seconds, 50 °C for 60 seconds, and 72 °C for 90 seconds, with a final extension of 10 min at 72 °C. Amplicons were then quantified using PicoGreen (Invitrogen) on an Infinite 200 PRO plate reader (Tecan, Morrisville, NC, USA). Once quantified, volumes of each of the products were pooled into a single tube so that each amplicon was represented in equimolar amounts. This pool was then cleaned up using AMPure XP Beads (Beckman Coulter, Brea, CA, USA) and quantified using a Qubit fluorometer (Invitrogen). After quantification, the molarity of the pool was determined and diluted down to 2 nM, denatured, and diluted to a final concentration of 6.75 pM with a 10% PhiX spike.

#### Pre-processing of sequencing data.

For the MiSeq sequencing run, priming sites and poor-quality bases were removed from the 5’ and 3’ ends of the sequences using the cutadapt v3.2 program (Martin, 2011). Because primer pads were used during the NextSeq 2000 sequencing run, no trimming of poor-quality bases at the 5’ and 3’ ends of the sequences was needed. For each sequencing run, we then discarded all reads that contained any unassigned bases (“Ns”). We also removed reads that had an expected error score greater than 3 for the forward reads and 8 for the reverse reads in the MiSeq run, and 2 for the NextSeq run both forward and reverse reads, using the filterAndTrim() function from the *DADA2* v1.32.0 R package (Callahan et al., 2016). We truncated the remaining reads at the first instance of a quality score less than 2. For the MiSeq run, 2.67 million reads total were sequenced across 20 samples for this project (22.45 million reads total, including data from other unpublished projects), averaging 133,718 reads per sample (± 40,377 SD; Table S11). For the NextSeq run, 67.34 million reads total were sequenced across 478 samples for this project (85.43 million reads total, including data from other unpublished projects), averaging 141,780 reads per sample (± 79,453 SD). Upon quality filtering, 95.75% (76.74% for the MiSeq run and 96.55% for the NextSeq run) of these reads were retained per sample, on average. We used the derepFastq(), dada(), and mergePairs() functions from *DADA2* to merge paired ends and infer the bacterial strains present, generating amplicon sequence variants (ASVs). Data from the two sequencing runs were then merged by sequence using *DADA2*’s mergeSequenceTables() function. We performed *de novo* chimera checking and removal on the combined data with the removeBimeraDenovo() *DADA2* function. Taxonomic classification was performed from kingdom to genus levels using the RDP classifier implemented in the assignTaxonomy() *DADA2* function with a modified SILVA SSU v138.1 training dataset (Quast et al., 2013; Wang et al., 2007), which included recent emended descriptions of the genus *Lactobacillus* (Schmidt, 2023; Zheng et al., 2020). As a result, 90.2% of our ASVs were assigned a phylum and 48.1% were assigned a genus. Filtered amplicon sequences were exported in fasta format using the write.fasta() function from the *seqinr* v4.2.36 R package (Charif & Lobry, 2007) and are available in our data repository (<https://figshare.com/s/94f2f48346754f8df88a>).

The ASV count table, the taxonomic classification table, and the metadata information (including bacterial titer and DNA concentration per sample) were converted into a phyloseq object that contained 99,979 ASVs, using the phyloseq() function from the *phyloseq* v1.48.0 and *speedyseq* v0.5.3.9021 R packages (McLaren, 2020; McMurdie & Holmes, 2013). For contaminant removal, we first deleted ASVs that were annotated as chloroplasts, mitochondria, or were unidentified at the phylum level. We then used the *decontam* v1.24.0 R package (Davis et al., 2018) with the stringent “either” frequency or prevalence method that identifies and removes contaminant ASVs based on frequency variation with DNA concentration (concentrations of zero were replaced by half of the minimum value = 0.0005 ng/µL) and prevalence in the extraction and sequencing blanks. With this method, 1,555 ASVs were flagged as contaminants and removed, resulting in 53,165 ASVs being retained in our dataset.

Finally, our data were rarefied to 30,000 sequences per sample, using the rarefy_even_depth() function, with the “replace = FALSE” argument, from the *phyloseq* R package, to control for among-sample differences in sequencing depth (Schloss, 2023, 2024; Weiss et al., 2017). Visual inspection of rarefaction curves, built with the *phyloseq* and *plyr* v1.8.9 R packages (Wickham, 2011), indicated that observed diversity had plateaued in the samples by this depth. To verify the relevance of rarefying our data, we first used the comparePhyloseq() function from the *biomeUtils* v0.22 R package (Shetty, 2021) to inspect loss in sample and read counts. Of our 619 samples that passed through the previous pre-processing steps (464 non-blank samples for this project only), 107 were removed during the rarefying process, reducing our ASV count to 44,488 (by 16.3%), with 281 rarefied samples from whole body specimens used in downstream analyses, accounting for 31,994 ASVs total. Additionally, we calculated the rarefaction efficiency index (REI) to assess loss in statistical power induced by rarefying, using the REI() R function (Hong et al., 2022). The average REI across host species in our dataset was 0.87 (SD = 0.16; median = 0.94), which suggests that there was limited loss in efficiency due to rarefying (close to 1). Because rarefying would remove important samples in our dissected tissue dataset, we instead used compositional normalization, computed with the tax_transform() function from the *microViz* v0.12.4 R package (Barnett et al., 2021), which converts read depth per bacteria into proportions (from 0 to 1) of total read depth per sample. Similarly, rarefied data were not used for our multivariate analyses based on Bray-Curtis distances, since compositional normalization has been shown to be a more robust normalization method for clustering on Bray-Curtis distance matrices (McMurdie & Holmes, 2014).

#### Taxonomic annotation refinement through phylogenetic analyses.

The 16S rRNA gene sequences (both our ASVs and NCBI-downloaded sequences) were not enough to obtain a well resolved *Wolbachia* phylogeny, as the supergroup A (Lo et al., 2002) was not monophyletic, separated into two groups. We thus used the 16S rRNA fragment insertion in a multi-locus tree (approach described in Ravenscraft et al. (2024)) to build a tree with higher resolution. Briefly, we first constructed a multi-locus tree, using IMG/M-ER (Chen et al., 2019) to download sequences of seven genes (*coxA*, *fbpA*, *ftsZ*, *gatB*, *hcpA*, *recA*, and the 16S rRNA gene; Ren et al., 2020) from *Wolbachia* genomes, Geneious Prime to manipulate sequences and concatenate them, MAFFT (Katoh et al., 2002) to run alignments, RAxML (Stamatakis, 2014) to run the phylogenetic analyses with partition blocks, and QIIME2 SEPP (Bolyen et al., 2019; Mirarab et al., 2012) with the fragment-insertion function (Janssen et al., 2018) to add our 16S rRNA ASV sequences to the backbone multi-locus tree. Thanks to this method, we obtained a seven-locus *Wolbachia* tree with higher bootstrap support overall and a monophyletic supergroup A (Figure S10).

Similar to *Wolbachia*’s phylogenetic analysis, our 16S rRNA sequences (both our ASVs and NCBI-downloaded sequences) were not enough for *Serratia* either, but the situation appeared to be worse. When running BLASTs of ASVs, hits were at 100% similarity and coverage to subjects from both *Serratia marcescens* and *Se. nematodiphila*. The same method as for *Wolbachia* was thus used to improve our tree topology and clade differentiation, with six genes selected to run a multi-locus *Serratia* phylogeny (*fumC*, *gyrB*, *phoA*, *pflB*/*bssA*, and *recA*; Alvaro et al., 2024; Bourdin et al., 2021). However, the backbone, multi-locus tree showed non-monophyletic placement for many *Serratia* species, still including polyphyletic groups for both *Se. marcescens* and *Se. nematodiphila*. Since the backbone, multi-locus phylogeny was not well resolved, we used autoMLST (Alanjary et al., 2019) to attempt to build a better multi-locus tree with a higher number of loci (×86). Focusing our efforts on improving the placement of *Se. nematodiphila*, a symbiont of entomopathogenic nematodes (Zhang et al., 2009), which could indicate nematode infection in our fireflies, we selected 17 genomes from only this species. Using autoMLST we identified MLSTs, aligned them to sequences from closest genomes from NCBI, and built the maximum likelihood tree with bootstrapping. While the bootstrap values were very high, and the topology and position of most *Serratia* species matched previous phylogenomic studies on this genus (Williams et al., 2022), *Se. nematodiphila* still appeared polyphyletic, nested within the *Se. marcescens* species. The branch length was also very short, indicating that *Se. nematodiphila* genomes are not much different from *Se. marcescens*’ ones. We thus returned to the 16S rRNA gene phylogeny (Figure S10), and used this one for *Serratia* phylotype assignment, mentioning species groups, like the “nematodiphila-marcescens” group, instead of separating them into two distinct phylotypes.

We note that some taxa are named after a reference species because of how close they are to this reference, but not necessarily nested within the species group. It is the case of *Spiroplasma* sv0009 which was considered as from the *Spiroplasma ixodetes* species (Figure S9) (Tully et al., 1995). Results from a PopART Templeton, Crandall, and Sing (TCS) network alignment run (Leigh & Bryant, 2015) using *Mollicutes* ASVs also concur with this placement, this ASV being very close to other *Sp. ixodetis* ASVs sister to the *Sp. ixodetis* reference sequence. While our 16S rRNA gene phylogenies show low bootstrap support for some taxon placements, most concur with previous research using multi-locus phylogenetic analyses (Gupta et al., 2019). Some *Williamsoniiplasma* ASVs had long branches on our 16S rRNA gene phylogeny, but we could classify them as from *Wi. lucivorax* or *Wi. somnilux* thanks to results from the PopART TCS network analysis, revealing one or less substitution between these ASVs and the ASVs which we confidently identified through phylogenetics.

We used the tax_agg() function from the *microViz* R package to aggregate taxonomy table at different taxonomic levels. Finally, we ran NCBI BlastN to annotate several abundant ASVs (those with an ASVsv ID number lower than 201) that were unidentified at the genus level through SILVA.

#### Bioinformatic analyses.

##### *Normalized stochasticity ratio analysis.*

To calculate the normalized stochasticity ratio (NST index), we used the tNST() function from the *NST* R package v3.1.9 (Ning et al., 2019), with the Bray-Curtis distance, 999 randomizations, and the default null model including simulated species occurrence frequency proportional to observed data and simulated richness in each sample maintained the same as observed data (null.model = “PF”). To test how NST index values within firefly species change between a full microbiome and a microbiome in which we removed *in silico* specific bacterial groups (*Mollicutes*, endosymbionts, or both *Mollicutes* and endosymbionts), we used the nst.boot() function from *NST* to perform bootstrapping (with 999 random draws) and run pairwise, two-tailed Wilcoxon signed-rank statistical tests. The output bootstrap values were extracted to calculate the 95% confidence intervals.

##### *Alpha and beta diversity.*

Indices of alpha diversity were calculated on rarefied data with non-aggregated (ASV level) taxonomic tables, using the alpha() function from the *microbiome* R package (Lahti & Shetty, 2017) (Table S4). For beta diversity, we first ran ordinations on Bray-Curtis distance matrices computed from non-rarefied, compositional data using the ordinate() and plot_ordination() functions from *phyloseq*. Because the percentage of explained variation on the first two axes was low on our principal coordinate (PCoA) analyses, we chose to run nonmetric multidimensional scaling (NMDS) analyses instead. We then ran permutational multivariate analysis of variance (PERMANOVAs) on rarefied, taxon-aggregated data, with the adonis2() function from the *vegan* v2.6.4 R package (Oksanen et al., 2016), and pairwise PERMANOVAs as post hoc tests using the pairwise.adonis() function from the *pairwiseAdonis* v0.4.1 R package (Martinez, 2020).

##### *Host phylogeny, geographic distances, microbiome dissimilarity, and Mantel tests.*

We build a multi-locus tree from a list of five genes**,** including the 16S rRNA, carbamoyl‐phosphate synthetase 2 (CAD), COI, UV opsin, and wingless (wg) genes (Figure S1). In this phylogeny, we grouped both *Pr. hebes* and unidentified species from the *Pr. versicolor* group under *Pr. quadrifulgens*, which are likely very close genetically (Lower et al., 2017). The *Pn ignitus* and *Pn. marginellus* species were used as representative of the *Pn. consanguineus* and *Pn. marginellus* groups. A click beetle (*Melatonus depressus*) COI gene was used as an outgroup. Gene nucleotide sequences were aligned using MUSCLE (Edgar, 2004) and concatenated in SeaView. Phylogenetic analysis was run with RAxML Blackbox on CIPRES with 999 bootstrap iterations. While the tree topology was identical to previous multi-locus phylogenies (Figure S1; Lower et al., 2017; Stanger-Hall et al., 2007), because of the small number of species and all the genes were not available for every firefly species, some minor differences in the topology of the tree can be noted when compared to the more recent phylogenetic analyses (e.g., earlier divergence of the branch containing *Lucidota atra* than *Pyractomena*, position of *Pn. scintillans* and *Pn. ignitus*) (Catalan et al., 2022; Höhna et al., 2024; Martin et al., 2019). Nevertheless, because we only used this phylogeny to generate genetic distance, which was calculated from branch length, the tree topology was not expected to bias our downstream analyses.

We generated a geographic distance matrix between firefly specimens using the st_as_sf() function from the *sf* v1.0.17 R package (Pebesma, 2018) to convert latitude and longitude decimal variables into an sf object and the st_distance() function to compute geographic distances in meters between collection points. The vegdist() function from *vegan* was used to generate the distance matrices, with the argument “binary = TRUE” (Bray-Curtis binary distance) or “binary = FALSE” (Bray-Curtis relative abundance distance). Bacterial presence was defined as any bacterial taxon with at least 1% relative abundance in a specimen and absence as less than 1% relative abundance in that specimen. Because the number of pairwise comparisons that we had to compute to generate pairwise matrix distances was excessively high, we applied a filter to our dataset, keeping only bacterial taxa that had more than 0.5% relative abundance in at least 5% of all firefly specimens.

These microbiome dissimilarity matrices were compared to a matrix of genetic distance and to geographic distances between firefly specimens through Mantel tests using the mantel() function from *vegan* with the pearson method and 9,999 permutations. We also performed partial Mantel tests using the mantel.partial() function from the same package, to determine the correlation between microbiome dissimilarity and host phylogeny while controlling for geography and the correlation between microbiome dissimilarity and geography while controlling for host phylogeny.

##### *Phylogenetic generalized least square models.*

To account for the effects of phylogenetic nonindependence in comparisons of firefly species-level covariates, we used phylogenetic generalized least square (PGLS) analyses, in addition to ordinary least square (OLS) analyses, with the argument correlation = corBrownian() to fit a Brownian motion model of evolution.

##### *Bacterial composition and differential abundance analysis.*

We used bargraphs, horizontal dotplots, and heatmaps to visualize bacterial composition and abundance across fireflies. The comp_barplot() function from the *microViz* R package, along with the *ggplot2* v3.5.1 package (Wickham, 2016), were used to generate barplots, representing relative abundance of main bacterial taxa across firefly species. The tax_select() function from *microViz* was used to rank specific bacteria to be drawn first in our barplots. To visualize in detail the rarefied abundance of abundant single bacterial taxa across firefly species, we used *ggplot2* to generate dotplots through custom functions. To visualize bacterial composition, titers, and prevalence in single graphs across non-firefly insects, juveniles, and different tissues, we generated composite heatmaps using the comp_heatmap() function from *microViz*.

To visualize differential bacterial abundance at multiple taxonomic ranks between firefly subfamilies, we used the taxatree_models(), taxatree_models2stats(), and taxatree_plots() functions from *microViz*. Because this function only runs linear models, we manually added results from *MaAsLin2* (Mallick et al., 2021) runs (as pink crosses) with taxa aggregated at the taxon and genus levels.

##### *Linear mixed models.*

We used the lmer() function from the *lme4* v1.1.35.5 R package (Bates et al., 2015) to fit linear mixed models and the glmer.nb() function from *lme4* to fit our data to a negative binomial distribution in generalized linear mixed models. Predicted means and 95% confidence intervals were visualized using the *ggeffects* v1.7.1 and *ggplot2* R packages (Lüdecke, 2018; Wickham, 2016).

### Results

#### Analysis of potential contaminants.

Among prevalent bacteria in fireflies, some may be contaminant or transient. We ran *MaAsLin2* multiple GLMMs within firefly species or genera to test whether there were negative correlations between the relative abundance of single bacteria and the specimen’s total absolute bacterial abundance, which may signal potential contaminant (Salter et al., 2014). Significant negative correlation corresponds to bacteria that were only relatively abundant in specimens with low total bacterial abundance. We found that in several firefly species some bacteria were only detected at high relative abundance in specimens that had low total bacterial abundance. These included *Brevibacterium*, *Flavobacterium*, *Halomonas*, *Methylobacterium*/*Methylorubrum*, *Salipaludibacillus*, *Sphingomonas*, and *Streptococcus*, with a negative correlation in four out of six species and species groups tested (Table S12). Using these negative correlations, we also found that some bacteria that were likely important in some firefly species, as mentioned in the main text, may be contaminant in other firefly species. This was the case for *Acinetobacter* in both *Pn. corruscus* and *Pn. marginellus* group, *Nubsella* in *Pn. corruscus*, *Pn. marginellus* group, and *Pn. scintillans*, *Wi. lucivorax* in *Pyractomena*, and *Wi. somnilux* in *Pn. marginellus* group.

#### Phylogenetic notes and strain composition.

Mollicute strains detected in our firefly specimens included *En. corruscae* (with its reference 16S rRNA sequence found here to be almost indistinguishable from that of *En. ellychniae*, which we thus refer to as another *En. corruscae* strain), *Spiroplasma ixodetis*, *Sp. lampyridicola*, *Tullyiplasma photuris*, *Williamsoniiplasma lucivorax*, *Wi. luminosum*, and *Wi. somnilux*. Only one previously isolated firefly-associated mollicute taxon was not represented in our dataset: *Sp. corruscae* (Hackett et al., 1996). One ASV (sv0736) was found to be somewhat related to *Sp. corruscae*, but did not form a monophyletic group with this species, instead grouping with *Sp. culicicola*. We also found a polytomy that included a new clade formed by two sister ASVs (referred to as “*Mycoplasma*-related” here), the *Mycoplasma* clade, and the clade including *Mesoplasma* and *Entomoplasma*. These *Mycoplasma*-related ASVs were only abundant in *Pn. pyralis* from the state of Virginia (USA) (Figure 6B). In addition, another ASV from the *Edwardiiplasma* genus was abundant in one of our tiger beetles (Figure S8 and S9).

Also detected in our dataset were mollicute strains from clades that have never been collected in fireflies before but have been detected in other invertebrates. These include ASVs most closely related to *Candidatus* Bacilloplasma and *Ca.* Lumbricincola (Kostanjšek et al., 2007; Nechitaylo et al., 2009) found in crustaceans and earthworms, *Moeniiplasma glomeromycotorum* (Naito et al., 2017) found in arbuscular mycorrhizal fungi, and *Mesoplasma* (Matteau et al., 2024; Tully et al., 1994) found in bees, beetles, and flies.

Regarding potential endosymbionts (i.e., symbionts residing inside the body or cells of another organism), most *Wolbachia* supergroup A strains from fireflies appeared closely related to those sampled from *Sympetrum striolatum* dragonflies, *Macropis europaea* bees, and *Myopa testacea* flies, while those from the supergroup B were closely related to *Wolbachia* strains from *Bemisia tabaci* whiteflies (Figure S10). For the *Rickettsia* genus, all four ASVs identified via the SILVA database and included in our phylogeny nested within the *Ri. bellii* group that contains strains associated with *Acyrthosiphon pisum* pea aphids. Otherwise, in the *Rickettsiaceae*-affiliated, ten ASVs were classified as *Ca.* Hepatincola and were related to *Ca.* Hepatincola porcellionum, an isopod symbiont (Dittmer et al., 2023). ASVs from other *Rickettsiaceae* clades such as *Ca.* Hemipteriphilus (*Bemisia tabaci* endosymbiont; Bing et al., (2013)), *Ca.* Trichorickettsia (paramecia and planarian endosymbiont; Modeo et al., (2020)), and two unclassified groups, were detected in our dataset. Among the *Rickettsiella* ASVs, at least six of them, including the most prevalent and abundant strains found in fireflies, grouped with *Rl. costelytrae* and *Rl. pyronotae* strains, which were sequenced from scarab beetle larvae (Leclerque et al., 2012). We assigned this group the taxon name “*Rickettsiella costelytrae*”.

#### Bacterial titer, alpha diversity, and beta diversity across firefly microbiomes.

At the species level, we found that *Pr. lucicrescens* had the highest median bacterial titer of all firefly species sampled (3.2 x 10^4^ copies of 16S rRNA gene per ng of DNA), with levels significantly higher than several bioluminescent *Photinus* species, including those in the *Pn. marginellus* group (median: 2.1 x 10^1^ copies of 16S rRNA gene per ng of DNA; emmeans: *z*-ratio = 4.29, *P* < 0.01), *Pn. pyralis* (median: 3.3 x 10^0^ copies of 16S rRNA gene per ng of DNA; emmeans: *z*-ratio = 7.76, *P* < 0.0001), and *Pn. scintillans* (median: 2.1 x 10^0^ copies of 16S rRNA gene per ng of DNA; emmeans: *z*-ratio = 5.55, *P* < 0.0001). In addition to *Pr. lucicrescens*, other species from the *Pr. versicolor* group, either from the *Pr. hebes* species or unidentified at the species level, had moderately high bacterial titers (median: 8.6 x 10^1^ and 4.0 x 10^2^ copies of 16S rRNA gene per ng of DNA, respectively), which were significantly higher than for *Pn. pyralis* titers (emmeans: *z*-ratio > 4.10, *P* < 0.01).

While the Shannon diversity index did not differ among microbiomes of firefly species, the inverse Simpson diversity index, the Evar index of evenness (Smith & Wilson, 1996), and the proportion of the least abundant bacterial ASVs were slightly but significantly higher in *Photinus* firefly microbiomes than in *Photuris* fireflies (emmeans: *z*-ratio > 3.52, *P* < 0.05;Figure S4; Table S4), suggesting a more even representation of the bacterial strains in *Photinus* compared to *Photuris*, mostly driven by low abundance bacteria.

In the *Lampyrinae* subfamily, when comparing across genera, we found that bacterial composition differed across *Photinus*, *Photuris*, and *Pyractomena* (pairwise adonis: *R*^2^ > 0.01, *F* > 1.83, FDR *P*-adj. < 0.05). Bacterial composition was not different between members of the *Lucidota* genus and *Photinus* fireflies (pairwise adonis: *R*^2^ = 0.01, *F* = 1.50, FDR *P*-adj = 0.10). This result appears to be driven at the firefly species level, by the microbiome similarity between *L. atra* and either *Pn. consanguineus* group, *Pn. cookii*, and *Pn. marginellus* group (pairwise adonis: *R*^2^ < 0.17, *F* < 1.26, FDR *P*-adj > 0.22). Even though *Wolbachia* was a genus characteristic of the microbiome of both *L. atra* and *Pn. marginellus* group, after removing likely endosymbionts from the dataset (i.e., *Rickettsia*, *Rickettsiella*, and *Wolbachia*), microbiome composition was still not significantly different between *L. atra* and either of these two *Photinus* species (pairwise adonis: *R*^2^ < 0.17, *F* < 1.21, FDR *P*-adj > 0.28). Nevertheless, bacterial community composition was significantly different across most species (considering only species that had replicate individuals in our dataset, i.e., excluding *Pn. cookii*, pairwise adonis: *R*^2^ > 0.01, *F* > 1.97, FDR *P*-adj < 0.05).

Species from the *Photuris versicolor* species group differed less in their microbiome composition with only minimal, yet significant, differences between unidentified species from the *Pr. versicolor* group and *Pr. hebes*, and between the *Pr. versicolor* group and *Pr. lucicrescens* (pairwise adonis: *R*^2^ > 0.02, *F* > 1.96, FDR *P*-adj < 0.05; Figure 3D). *Pr. hebes* and *Pr. lucicrescens* appeared to share similar bacterial communities (pairwise adonis: *R*^2^ = 0.03, *F* = 0.95, FDR *P*-adj = 0.50), despite significantly different absolute bacterial abundances (emmeans: *z*-ratio = 4.18, *P* < 0.01; Figure 1C).

#### Influence of species’ ecology and life history traits.

##### *Firefly microbiomes do not vary between bioluminescent and non-bioluminescent species.*

We assessed whether bacterial density, richness, and composition differed between bioluminescent and non-bioluminescent species. Since phylogenetic relatedness was a main factor influencing bacterial composition (Figure S5), we used statistical tests that account for fireflies’ degree of relatedness. We found no significant difference in total bacterial density and bacterial richness between bioluminescent and non-bioluminescent species, whether we controlled for phylogenetic relatedness or not (PGLS and OLS: *F* < 1.62, *P* > 0.23; Figure S6). Similarly, bacterial composition changed only marginally between bioluminescent and non-bioluminescent species when considering only adult specimens (PERMANOVA with bioluminescence nested within genus within subfamily and species nested within genera within subfamily as covariate: *R*^2^ < 0.01, *F*_1,252_ = 1.32, *P* = 0.043), and when taking juveniles into account (all bioluminescent) (PERMANOVA with bioluminescence nested within genus within subfamily and sex/stage nested within species within genera within subfamily as covariate: *R*^2^ < 0.01, *F*_1,265_ = 1.32, *P* < 0.05). Based on *MaAsLin2* multiple GLMM analysis, no bacterial taxa significantly differed in their relative abundance between bioluminescent and non-bioluminescent species. These results suggest that firefly microbiomes do not vary between bioluminescent and non-bioluminescent species.

##### *Dietary effects are confounded with host genetic relatedness.*

We next tested whether the microbiome changed between firefly species that eat or are not known to eat as adults, and across adult diets. We found that, when not controlling for phylogenetic distance between firefly species, bacterial density was higher in species that eat as adults compared to those not known to eat as adults (OLS: *F* = 16.42, *P* < 0.01; Figure S6), whereas bacterial richness did not vary. When analyzing different diets, predatory fireflies harbored a higher bacterial density than non-eating fireflies (OLS emmeans: *t* = 3.08, *P* < 0.05), while the sap/nectar feeder category was represented by only one species making it difficult to formally compare this group to others, although it showed similar total bacterial density to predatory fireflies. However, the significance of this effect was drastically reduced when controlling for phylogenetic distance between firefly species, with bacterial density only being marginally significantly higher in species eating as adults compared to non-eating ones (PGLS: *F* = 4.42, *P* = 0.07).

Bacterial composition was different between eating and non-eating firefly species (PERMANOVA with adult eating/non-eating factor nested within genus within subfamily and species nested within genera within subfamily as covariate: *R*^2^ = 0.02, *F*_1,253_ = 6.76, *P* < 0.001) and across firefly diets (PERMANOVA with adult eating/non-eating factor nested within genus within subfamily and species nested within genera within subfamily as covariate: *R*^2^ = 0.02, *F*_1,253_ = 6.76, *P* < 0.001; Figure S6), with the difference being the strongest between non-eating and predatory fireflies (pairwise adonis: *R*^2^ = 0.08, *F* = 22.81, FDR *P*-adj < 0.001). Although we controlled for host species in our *MaAsLin2* models, results from differentially abundant bacterial taxa across dietary groups recapitulated our results for the interspecies comparisons (described in the main text), notably where sap/nectar drinkers, the *Pn. corruscus*, harbored high abundance of *Entomoplasma corruscae*; non-eaters, which included other lampyrines, harbored high abundance of *Stenotrophomonas*; and predatory *Photuris* fireflies harbored high abundance of *Achromobacter* (MaAsLin2: coef > 3.10, FDR *P*-adj < 0.0001; Table S7). These results suggest that while fireflies eating as adults may harbor a higher bacterial density and that firefly species diets may influence the composition of their microbiomes, the degree of relatedness is a major factor influencing firefly microbiomes that needs to be accounted for.

##### *Bigger fireflies harbor higher bacterial density but lower bacterial diversity.*

Because host body size is known to influence bacterial abundance, richness, and diversity in the microbiome (Godon et al., 2016, p. 20; Kwong et al., 2017), with bigger hosts providing larger habitats to microbes, we assessed whether bacterial density and diversity varied with the average size of firefly species. We found that bigger firefly species had higher absolute bacterial abundance (PGLS: *t* = 2.37, *P* < 0.05; Figure S7), while bacterial richness followed the opposite trend (PGLS: *t* = -2.42, *P* < 0.05). With a body length of 17.5 cm on average, the *Pr. lucicrescens* adults were thus more likely to harbor high densities of bacteria than fireflies with a smaller body size, but lower bacterial richness.

#### Negative correlations between bacterial density and diversity.

We found a weak negative correlation between absolute bacterial abundance and bacterial richness and diversity (OLS and PGLS: *t* < -2.15, *P* < 0.06; Figure S7). This result highlights a possibly important contrast across firefly species, suggesting that these insects either harbor a low density of diverse bacteria, as for *Lucidota atra* and most *Photinus* species, or a bacterial community dominated by a few strains with high absolute abundance, as in *Pn. corruscus*, *Photuris* species, and *Py. borealis*.

#### Notes on bacterial composition in fireflies and related beetles.

It is important to note that a much larger number of *Pn. pyralis* specimens was represented in our dataset (Figure 1A), and bacterial prevalence and relative abundance at the family, subfamily, and genus levels may be biased in favor of this overrepresented species.

*Lucidota atra* fireflies could be associated with *Wolbachia* from either the supergroup A or B (Figure 4A; Figure S11). The microbiome of the less sampled *Pn. cookii* and *Pn. consanguineus* group was not dominated by mollicutes, but had a high abundance of *Stenotrophomonas* (Figure S11).

Few of the abundant firefly-associated bacterial taxa were retrieved in non-lampyrid elateroid beetles (Figure S8). Compared to close relatives, such as click beetles (*Elateridae*) and soldier beetles (*Cantharidae*), and when looking at the bacterial genus level, fireflies hosted significantly higher abundance of *Elizabethkingia* (*Flavobacteriales*), *Leucobacter* (*Micrococcales*), *Nubsella* (*Sphingobacteriales*), *Sphingobacterium* (*Sphingobacteriales*), *Stenotrophomonas* (*Xanthomonadales*), *Williamsoniiplasma* (*Mollicutes*), and *Wolbachia* (*Rickettsiales*) (MaAsLin2: coef > 3.78, FDR *P*-adj < 0.01; Figure S11), while *Alysiella* (*Burkholderiales*) and *Flavobacterium* (*Flavobacteriales*) were more abundant in non-firefly elateroid beetles compared to fireflies (MaAsLin2: coef > 4.59, FDR *P*-adj < 0.05; Figure S8). At the bacterial taxon level, aside from the previously mentioned genera, *Spiroplasma* *ixodetis* and *Wolbachia* supergroup B appeared to be enriched in fireflies compared to click and soldier beetles (MaAsLin2: coef > 4.10, FDR *P*-adj < 0.01).

#### Bacteria positively and negatively co-occur in fireflies.

Positive or negative co-occurrence among members of a community can indicate potential competitive or cooperative interactions. From our co-occurrence analysis, we found that some bacteria were more likely to be found on their own in fireflies, having low α MLE affinity index values (i.e., maximum likelihood estimate log odds ratio α of conditional occurrence probability; Mainali et al., (2022)). These included bacteria that were rarely found in the same firefly specimen as other abundant bacteria, such as *Rickettsia bellii* in *Pn. corruscus* and *Pn. marginellus* group; *Wolbachia* supergroup A in *Pn. marginellus* group; *Lactococcus* and *Serratia marcescens-nematodiphila* in *Pn. scintillans*; *Lonsdalea* and the *Mycoplasma*-related group in *Pn. pyralis*; and *Sp. ixodetis*, *Sp. lampyridicola*, and *Wolbachia* B in *Photuris* fireflies (Figure S13).

We also found some positive co-occurrence patterns within firefly species. In *Pn. pyralis* *Acinetobacter*, *Pseudochrobactrum*, *Pseudomonas*, *Se. marcescens-nematodiphila*, and *Stenotrophomonas* appeared to be commonly coexisting in the same hosts (Figure S13). Interestingly, in both *Pn. pyralis* and *Pn. marginellus* group, *Pseudomonas* were significantly positively associated with *Stenotrophomonas* and *Rl. costelytrae*. In *Photuris* fireflies, we found a tripartite significant positive co-occurrence of *Achromobacter*, *Curvibacter*, and *Elizabethkingia*.

#### Effects of geography, urbanization, and temperature on the microbiome of fireflies.

##### *Geographic location affects the microbiome of fireflies.*

In *Photinus corruscus*, bacterial titers were generally low in lower latitudes, such as southwestern Connecticut (swCT) (median: 4.0 x 10^0^ copies of 16S rRNA gene per ng of DNA) and South New Jersey (sNJ) (median: 3.6 x 10^1^ copies of 16S rRNA gene per ng of DNA), but could reach much higher levels in northern latitudes like in western Massachusetts (wMA) (median: 8.4 x 10^5^ copies of 16S rRNA gene per ng of DNA) and southwestern Vermont (swVT) (median: 2.5 x 10^7^ copies of 16S rRNA gene per ng of DNA) (GLMM Wald test on normalized titer across decimal latitudes: *Χ*^2^ = 326.39, *P* < 0.0001). *Rl. costelytrae*, was abundant in *Pn. corruscus* specimens from nwCT, swCT, swVT, and wMA, but virtually absent from specimens collected in seNY and sNJ (seNY or sNJ *vs.* nwCT or wMA, MaAsLin2: coef > 6.07, FDR *P*-adj < 0.001), suggesting a potential preference of *Rl. costelytrae* for easternmost states. *Williamsoniiplsma luminosum* and *Wi. lucivorax* appeared to be virtually absent from the very few *Pn. pyralis* specimens collected in East Tennessee (eTN) (*n* = 1) and North Texas (*n* = 1).

Microbiome differences across *Photuris versicolor* group specimens from different regions was mainly driven by the strong difference in microbiome composition between specimens from central Pennsylvania (cPA) and southeastern PA (sePA) (pairwise adonis: *R*^2^ = 0.07, *F* = 3.06, FDR *P*-adj. < 0.01; Figure 6C), although confounding factors may have biased our results, including habitat types (e.g., almost all of the specimens collected in sePA were collected in the city). Bacterial titers were particularly high in specimens from cPA (median: 6.0 x 10^4^ copies of 16S rRNA gene per ng of DNA *vs.* 3.4 x 10^2^ in specimens from other regions), especially compared to those from sePA (emmeans: *z*-ratio = 4.13, *P* < 0.001).

##### *Bacterial composition varies across habitat types in unidentified species from the* Pr. versicolor *group.*

Bacteria, such as members of the *Hafniaceae* family and *Yersinia* were more abundant in habitats that combine both meadows and forests compared to the city (MaAsLin2: coef > 3.86, FDR *P*-adj < 0.05; Figure S16).

##### *Temperature slightly affects the microbiome of* Pn. pyralis*.*

The microbiome of *Pn. pyralis* appeared to change with the temperature at specimen collection, both in terms of absolute abundance (GLMM Wald test on normalized titer across decimal temperatures: *Χ*^2^ = 5.17, *P* < 0.05), with higher bacterial titers found at higher temperatures, and in composition (PERMANOVA: *R*^2^ = 0.02, *F* = 2.48, *P* < 0.05). However, we did not find any bacterial taxon detected in more than 1% of all *Pn. pyralis* specimens that had an abundance changing with temperature, implying a contribution of only rare bacteria.

#### Alpha diversity and bacterial composition in juveniles.

When looking at alpha diversity, comparing larvae to adults across all fireflies, we found that they had a marginally higher bacterial richness (emmeans: *z*-ratio = 2.19, *P* = 0.072; Figure S18) and significantly higher diversity (emmeans: *z*-ratio = 2.70, *P* < 0.05). Larvae were also different from adults in terms of bacterial composition (pairwise adonis: *R*^2^ = 0.04, *F* = 3.60, FDR *P*-adj < 0.001).

Alpha diversity in the microbiomes of *Pyractomena borealis* appear to change through development. While larvae from this species harbored marginally more bacterial ASVs than their adult conspecifics (median: 593 *versus* 79, respectively; emmeans: *z*-ratio = 3.65, *P* = 0.058; Figure S4), the dominance index of Berger-Parker was much lower in the microbiome of our single *Py. borealis* pupa that passed rarefying compared to two adult males (0.07 *vs.* 0.87 on average, respectively), but this difference was not significant, likely due to the small sample size. This suggests a higher bacterial richness in larvae and a more even representation of bacteria in pupae, potentially linked to the low, but quantifiable, bacterial density in this developmental stage (median: 1.9 x 10^1^ copies of 16S rRNA gene per ng of DNA *vs.* 7.8 x 10^1^ in adult males; Figure S18).

Our *Photinus consanguineus* group larva had a high abundance of *Stenotrophomonas*, *Pseudomonas*, and *Rhodococcus* bacteria, but not *Williamsoniiplasma* spp. (Figure S19). In *Pyractomena*, *Acinetobacter*, *Brevundimonas*, unidentified *Microbacteriaceae*, *Rhodococcus*, and *Spirosoma* had a significantly higher abundance in juveniles than in adult males (MaAsLin2: coef > 1.99, FDR *P*-adj < 0.05).

#### Notes on tissue-specific analyses.

We found that bacterial abundance and composition were different across firefly tissues, with species-specific compositional and density patterns (Figure 8A–8E). Because reproductive organs can have high amounts of host DNA, we used raw bacterial titers for our analyses rather than titers normalized by total sample DNA concentration. Nevertheless, compared to models fitting with normalized titers, using raw titers did not drastically alter the results, except for the *Pr. versicolor* group where differences of bacterial abundance across tissues were slightly amplified. In terms of composition, only non-rarefied data was used here because rarefying removed key samples that included potentially impactful bacteria such as *Wolbachia*. Instead, we used proportion-normalized relative abundance data (i.e., compositional).

In *Pn. corruscus*, while the highest bacterial titers were seen in the carcass, they were not significantly higher than in the gut and reproductive organs where titers remained relatively high (emmeans: *t*-ratio < 1.76, *P* > 0.18). Using *MaAsLin2* analyses, we found a slightly higher abundance of *Brachybacterium* in the gut than in reproductive organs (MaAsLin2: coef = 0.81, FDR *P*-adj < 0.0001; Table S9), while unidentified bacteria from the *Rhizobiales* order and *Wolbachia* from the supergroup A were more abundant in reproductive organs than in the carcass (MaAsLin2: coef > 3.87, FDR *P*-adj < 0.05). We also note that *Rickettsia bellii* was detected in the gut of one *Pn. corruscus* male, whereas *Serratia marcescens-nematodiphila* was detected at high relative abundance in the gut of a female. As previously observed (Rooney & Lewis, 2000), when we dissected two *Pn. corruscus* specimens that were collected in November (a female and a male), we noticed numerous enlarged fat body structures surrounding the gut (Figure S22). However, a very limited amount of bacteria was detected in these structures.

We note that bacterial taxa characteristic of multiple firefly species, such as *Acinetobacter*, *Achromobacter*, and *En. corruscae* were detected in most tissues of several *Pn. pyralis* specimens (as well as other dissected firefly species), but at very low abundance, raising the possibility of reads being detected from cross contamination or sequencing errors.

### Discussion

#### Geographic variation in mollicutes.

Although they were not always present, characteristic bacterial symbionts from the *Mollicutes* (or *Wolbachia* in *Photinus marginellus* complex) could be detected at high abundance in specimens from different regions. For example, in our study, *Williamsoniiplasma lucivorax* was the most abundant bacterial taxon in at least one *Pn. pyralis* specimen collected from central Pennsylvania (PA), southeastern PA, central Tennessee, and western Virginia, USA (Figure 6B). Other studies have also documented the presence of this bacterial species in *Pn. pyralis* specimens collected from the state of Maryland (Hackett et al., 1992). Additionally, in our US-based study, despite being more abundant in less urbanized habitats, the mollicute *Tullyiplasma photuris* could be detected in *Photuris* fireflies from all types of habitats (Figure S16), and was also sampled from *Photuris* sp. fireflies in Ecuador (Hackett et al., 1992).

#### Sex-specific microbiome differences in *Photinus*.

We found curious differences between female and male microbiomes of the *Pn. pyralis* and *Pn. scintillans* species. While the bacterial composition and richness did not differ between sexes (Figure 3A–3F; Figure S4), females had a much higher bacterial density than males in these two species (Figure 7E). Similar to larvae, females from these two *Photinus* species experience more contact with their microbe-rich environment than males, spending more time on the soil or on the vegetation, where they may acquire more environmental bacteria than male conspecifics. This might especially be the case for the *Pn. scintillans* females which are brachypterous (i.e., lack developed wings) and were found to have an even higher bacterial density than *Pn. scintillans* males as compared to *Pn. pyralis* females and males. Like most other fireflies, females from these two species also receive nutrient-rich spermatophores from males (Al-Wathiqui et al., 2016; Faust et al., 2019). It is unclear, however, whether bacteria can be horizontally transmitted during mating and whether spermatophore transfers are associated with higher bacterial loads in females. It is also not known whether the repeatedly evolved extreme sexual dimorphism in fireflies (South et al., 2011), characterized by complete female paedomorphosis (i.e., morphologically similar to larvae), enforces drastically different microbiomes between sexes.

#### Hypothetical functions of firefly-associated bacteria.

In addition to the mollicutes being common and abundant microbial associates of fireflies, we found that specimens from the *Pn. marginellus* group were frequently infected with *Wolbachia*. The functional role of these bacteria in fireflies is still unknown, but as shown in a multitude of insects, non-random symbiotic associations are linked to net fitness advantages (for beneficial symbionts) or disadvantages (for parasitic symbionts) to the host.

Here, we found that two *Spiroplasma* species (*Mollicutes*), *Sp. ixodetis* and *Sp. lampyridicola*, from two separate clades (ixodetes and apis), commonly associate with fireflies from the *Pr. versicolor* group. Previous studies have shown that *Pn. corruscus* fireflies also commonly harbor *Sp.* *corruscae* (Green et al., 2021; Hackett et al., 1992, 1996; Wedincamp et al., 1996). Spiroplasmas have been implicated in diverse functional roles as insect associates, regularly found to be commensals, pathogens, and reproductive manipulators (Bastian et al., 2012; Duron et al., 2008; Hurst et al., 1999; Pollmann et al., 2022; Williamson & Poulson, 1979), but occasionally serving defensive functions (Awuoche et al., 2025; Jaenike et al., 2010; Paredes et al., 2016; Xie et al., 2010). The functional roles of these spiroplasmas, other mollicutes, and other bacteria associated with fireflies have never been assessed and remain completely unknown. Based on bacterial characteristics in culture and genomic data, we present here speculations as to what their impact on firefly physiology and fitness could be. Importantly, while none of the firefly specimens in our study showed signs of illness at collection, it is also possible that the mollicutes detected in fireflies are parasitic.

First, many of the firefly-associated mollicutes—including *Entomoplasma ellychniae*, *Sp. corruscae*, *Sp. ixodetis*, *Sp. lampyridicola*, *Williamsoniiplasma* spp., but not *En. corruscae* and *T. photuris*—require sterol in their environment to survive (Hackett et al., 1996; Stevens et al., 1997; Tully et al., 1989, 1994, 1995; Williamson et al., 1990). Coincidentally, like other insects, fireflies cannot synthesize sterols but require this compound in order to produce their anti-predator toxin lucibufagin and diverse lipid-containing molecules (Behmer & Nes, 2003; Eisner et al., 1978). Fireflies most likely obtain sterol from their diet as larvae, when they prey on cholesterol-rich invertebrates. Sterol-requiring mollicutes may thus take advantage of the high cholesterol intakes in firefly guts at the larval stage, interfering with the assimilation of this nutrient by the host. If that is the case, firefly-associated mollicutes would thus behave as parasites, competing with their host for a resource with a concentration that may fluctuate in firefly guts over time. But it has been shown that mollicutes can have low impacts on insect hosts, persisting at low abundance while occasionally benefitting from excess host cholesterol intake (Paredes et al., 2016). Such host-microbe competition for lipids has been documented between the mollicute *Sp. poulsonii* and its *Dr. melanogaster* host (Herren et al., 2014), the former becoming protective by depleting resources otherwise available to other parasites (Awuoche et al., 2025; Paredes et al., 2016). Studies quantifying hemolymph lipids in fireflies infected or not by mollicutes and other parasites should be conducted to test this possibility.

A second hypothetical function is that mollicutes may synthesize defensive toxins for their host. As previously hypothesized (Smedley et al., 2017), they may utilize diet-derived sterols to participate in the synthesis of steroids such as the toxin lucibufagin and several insect hormones. While its anti-predatory role has been demonstrated (Eisner et al., 1978, 1997; Gronquist et al., 2006; Smedley et al., 2017; Tyler et al., 2008), lucibufagin may also have antimicrobial activity, since similar toxins from toads deter trypanosome protozoans (Rodriguez et al., 2021). The production of defensive compounds by symbionts has been demonstrated in a few other insects, including beetles (Kellner, 2002). In addition, one related, yet often overlooked, symbiont-mediated defensive mechanism is the reduction of appeal or palatability to predators. Some fireflies are less palatable than others, as suggested by a study showing that adults from a few lampyrine species are attacked and bitten by *Photuris* but not further eaten (Lewis et al., 2012). The defensive role of firefly-associated microbes should be assessed in future studies.

Third, many insects with nutrient-poor diets benefit from nutrient-providing symbionts (Douglas, 2009). Since several lampyrine fireflies are considered to be facultative eaters as adults with limited food intake, their diet might be nutritionally unbalanced at this stage. However, our results do not support this hypothesis since we found that fireflies that do not eat as adults tend to have a reduced total bacterial load compared to eaters (Figure S6). The examination of the lampyrine-associated mollicute draft genomes (Lo et al., 2018) suggest limited capabilities to synthesize key nutrients, such as amino acids and B-vitamins, despite being able to catabolize arginine and produce ammonia, ATP, and carbon dioxide. In contrast to the lampyrines, predatory photurine fireflies harbored a high density of bacteria in the gut (Figure 8E). In these fireflies, mollicutes could conceivably be involved in digestive or detoxification functions. With their exclusively predatory diet, the larval stage might also benefit from digestive symbionts able to process prey materials. Future studies should investigate the role of gut bacteria in firefly digestion, detoxification, and nutrition at diverse life stages.

Fourth, reproductive manipulation is a common behavior of arthropod-associated endosymbionts, including *Rickettsia*, *Rickettsiella*, *Wolbachia*, and some *Spiroplasma* strains (Duron et al., 2008; Rosenwald et al., 2020; Werren et al., 2008). Here we found that the specimens from the *Pn. marginellus* group collected in different regions of the Northeastern US were frequent hosts of *Wolbachia* from the diverse supergroup A (Figure 5A and 6B). Strains from this endosymbiont supergroup infect a variety of insects and often use cytoplasmic incompatibility to manipulate the reproduction of their hosts (Liu et al., 2023), which include beetles (Giordano et al., 1997). Reproductive manipulation performed by maternally inherited endosymbionts often leads to sex ratio distortion toward females (Cordaux & Gilbert, 2025). While the sex ratio is believed to be male biased in synchronous *Photinus* species (López-Palafox et al., 2020), it is not known whether it is biased in *Pn. marginellus* group and whether *Wolbachia* might contribute to sex ratio distortion in fireflies. This high *Wolbachia* A prevalence in this firefly species group is to be contrasted with the more diffuse and random infections of other endosymbionts in other firefly species (Figure 5A). Future research should investigate whether endosymbionts act as reproductive manipulators in fireflies.
